# FisB relies on homo-oligomerization and lipid-binding to catalyze membrane fission in bacteria

**DOI:** 10.1101/2020.09.25.313023

**Authors:** Ane Landajuela, Martha Braun, Christopher D. A. Rodrigues, Alejandro Martínez-Calvo, Thierry Doan, Florian Horenkamp, Anna Andronicos, Vladimir Shteyn, Nathan D. Williams, Chenxiang Lin, Ned S. Wingreen, David Z. Rudner, Erdem Karatekin

## Abstract

Little is known about mechanisms of membrane fission in bacteria despite their requirement for cytokinesis. The only known dedicated membrane fission machinery in bacteria, FisB, is expressed during sporulation in *Bacillus subtilis* and is required to release the developing spore into the mother cell cytoplasm. Here we characterized the requirements for FisB-mediated membrane fission. FisB forms mobile clusters of ∼12 molecules that give way to an immobile cluster at the engulfment pole containing ∼40 proteins at the time of membrane fission. Analysis of FisB mutants revealed that binding to acidic lipids and homo-oligomerization are both critical for targeting FisB to the engulfment pole and membrane fission. Experiments using artificial membranes and filamentous cells suggest FisB does not have an intrinsic ability to sense or induce membrane curvature but can bridge membranes. Finally, modeling suggests homo-oligomerization and trans interactions with membranes are sufficient to explain FisB accumulation at the membrane neck that connects the engulfment membrane to the rest of the mother cell membrane during late stages of engulfment. Together, our results show that FisB is a robust and unusual membrane fission protein that relies on homo-oligomerization, lipid-binding and the unique membrane topology generated during engulfment for localization and membrane scission, but surprisingly, not on lipid microdomains, negative-curvature lipids, or curvature-sensing.

## INTRODUCTION

Membrane fission is a fundamental process required for endocytosis^1^, membrane trafficking^2^, enveloped virus budding^3^, phagocytosis^4^, cell division^5^ and sporulation^6-8^. During membrane fission, an initially continuous membrane divides into two separate ones. This process requires dynamic localization of specialized proteins, which generate the work required to merge membranes^9-13^. Dynamin^14^ and the endosomal sorting complex required for transport III (ESCRT-III) catalyze many eukaryotic membrane fission reactions^15^. Both fission machineries bind acidic lipids, assemble into oligomers, and use hydrolysis of a nucleoside triphosphate (GTP or ATP) to achieve membrane fission. However, membrane fission can also be achieved by friction^16^, stress accumulated at a boundary between lipid domains^17^, forces generated by the acto-myosin network^18-21^ or protein crowding^22^. By contrast, very little is known about membrane fission in bacteria, even though they rely on membrane fission for every division cycle.

We previously found that fission protein B (FisB) is required for the final membrane fission event during sporulation in *B. subtilis*^23^. When nutrients are scarce, bacteria in the orders *Bacillales* and *Clostridiales* initiate a developmental program that results in the production of highly resistant endospores^24^. Sporulation starts with an asymmetric cell division that generates a larger mother cell and a smaller forespore (Figure 1A). The mother cell membranes then engulf the forespore in a process similar to phagocytosis. At the end of engulfment, the leading membrane edge forms a small pore. Fission of this membrane neck connecting the engulfment membrane to the rest of the mother cell membrane releases the forespore, now surrounded by two membranes, into the mother cell cytoplasm (Figure 1A,B). At this late stage the mother nurtures the forespore as it prepares for dormancy. Once mature, the mother cell releases the spore into the environment through lysis. Spores can withstand heat, radiation, drought, antibiotics, and other environmental assaults for many years^25-28^. Under favorable conditions, the spore will germinate and restart the vegetative life cycle.

**Figure 1.**
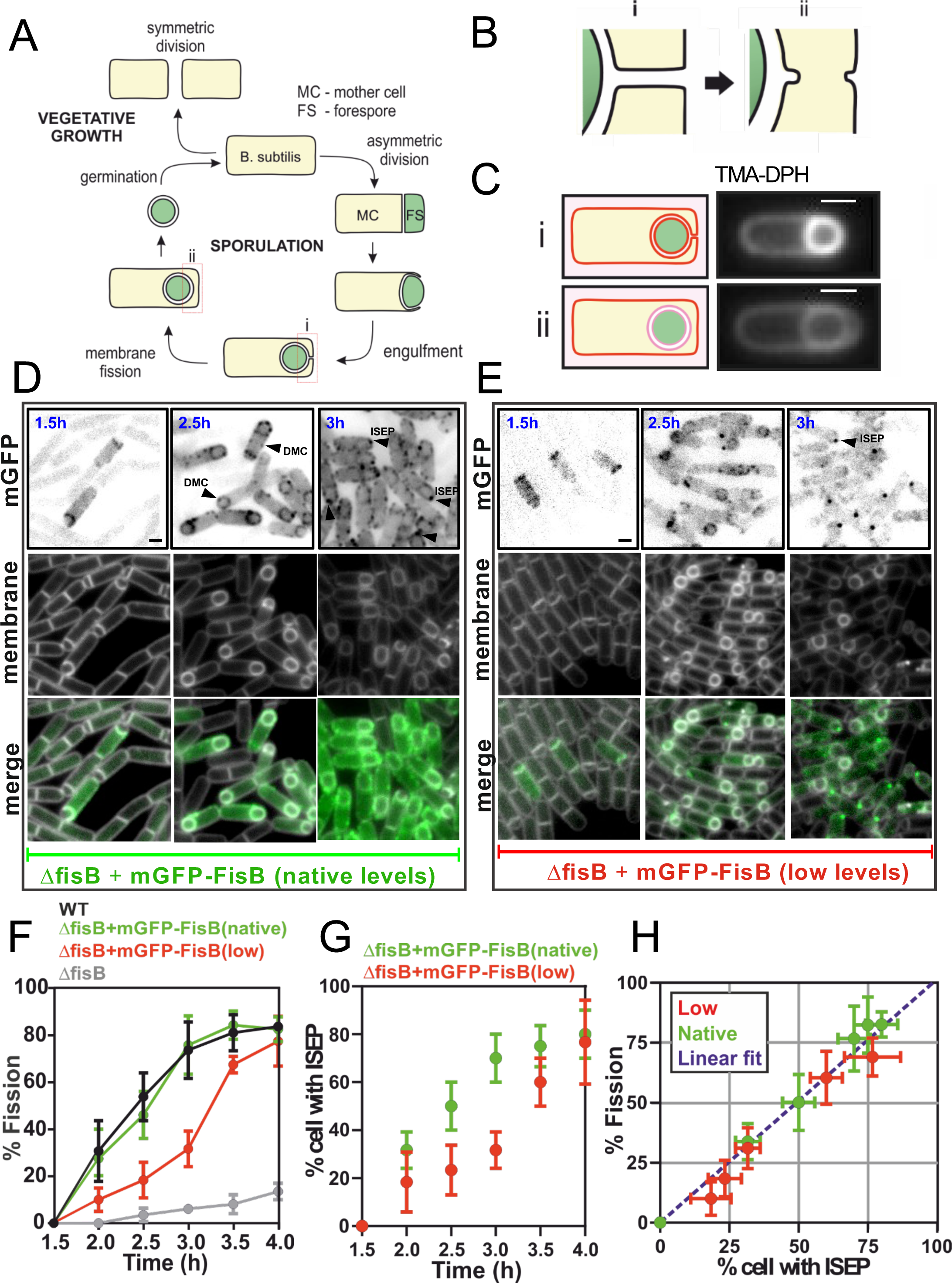
Membrane fission during sporulation is nearly always accompanied by accumulation of a FisB cluster at the fission site. **A**. Vegetatively growing cells enter sporulation when nutrients become scarce. Asymmetric division creates a forespore (FS) and a mother cell (MC). The MC engulfs the FC in a phagocytosis-like event. At the end of engulfment, a membrane neck connects the engulfment membrane to the rest of the MC (i). Fission of the neck (ii) releases the FS, now surrounded by two membranes, into the MC cytoplasm. Once the forespore becomes a mature spore, the MC lyses to release it. **B**. The membrane fission step shown in more detail. **C**. Detection of membrane fission. The lipophilic dye TMA-DPH does not fluoresce in the aqueous solution and crosses membranes poorly. If membrane fission has not yet taken place, the dye has access to the engulfment, FS and MC membranes, thus shows intense labeling where these membranes are adjacent to one another (i). If fission has already taken place, the dye labels internal membranes poorly (ii). **D.** Images show mGFP-FisB (strain BAM003, native expression level) at indicated times during sporulation. Membranes were visualized with TMA-DPH. Examples of sporulating cells with mGFP-FisB enriched at the septum (1.5 h), forming dim mobile cluster (DMC; 2 h) and with a discrete mGFP-FisB focus at the cell pole (intense spot at engulfment pole, ISEP, 3 h) are highlighted with arrowheads. **E.** Similar to D, but using a strain (BAL003) that expresses mGFP-FisB at lower levels in a *ΔfisB* background. **F.** Time course of membrane fission for wild-type cells, *ΔfisB* cells, or *ΔfisB* cells complemented with mGFP-FisB expressed at native (BAM003) or low levels (BAL003). Lower expression of mGFP-FisB leads to a delay in membrane fission kinetics. **G.** The percentage of cells with an intense spot at the engulfment pole (ISEP) for low and native level expression of mGFP-FisB as a function of time into sporulation. **H**. Correlation between percentage of cells that have undergone fission and percentage of cells having an ISEP for all time points shown in F and G. The fitted dashed line passing through the origin has slope 1.06 (*R*^2^ = 0.9). Scale bars represent 1 μm.

Conserved among endospore-forming bacteria, FisB is a mother-cell transmembrane protein expressed under the control of the transcription factor, σ^E^, after asymmetric division^29^. In sporulating cells lacking FisB, engulfment proceeds normally but the final membrane fission event, detected using a lipophilic dye, is impaired^23^ (Figure 1C,F and S1 Appendix Fig. 1A). During engulfment, FisB fused to a fluorescent protein forms dim, mobile clusters in the engulfment membrane (Figure 1D,E, Movie 1). When the engulfing membranes reach the cell pole, approximately 3 hours (t = 3h) after the onset of sporulation, a cluster of FisB molecules accumulates at the pole forming a more intense, immobile focus, where and when fission occurs (Figure 1D,E, Movie 2).

We had previously reported^23^ that FisB interacts with cardiolipin (CL), a lipid enriched at cell poles^30-32^ whose levels increase during sporulation^33^ and is implicated in membrane fusion^34-36^ and fission reactions^37^. In addition, CL was reported to act as a landmark for the polar recruitment of the proline transporter ProP, and the mechanosensitive channel MscSm^38,39^. Thus, it seemed plausible that CL could act as a landmark to recruit FisB to the membrane fission site and facilitate membrane fission. Apart from this hypothesis, no information has been available about how FisB localizes to the membrane fission site and how it may drive membrane scission.

Here, we determined the requirements for FisB’s sub-cellular localization and membrane fission during sporulation. Using quantitative analysis, we find small clusters of ∼12 FisB molecules diffuse around the mother cell membrane and ∼40 copies of FisB accumulate at the fission site as an immobile cluster to mediate membrane fission. When FisB expression was lowered, ∼6 copies of FisB were sufficient to drive membrane fission, but fission took longer. Unexpectedly, FisB dynamics and membrane fission are independent of both CL and phosphatidylethanolamine (PE), another lipid implicated in membrane fusion and fission. We found FisB binds phosphatidylglycerol (PG) with comparable affinity as CL, after adjusting for charge density. Thus, we suspect that, as a more abundant lipid in the cell, PG can substitute for CL to bind FisB. We tested other factors that may be important for the sub-cellular localization of FisB and membrane fission. We found FisB dynamics are independent of flotillins, which organize bacterial membranes into functional membrane microdomains^40^, cell wall synthesis machinery, and proton or voltage gradients across the membrane. Using mutagenesis, we show that both FisB oligomerization and binding to acidic lipids are required for proper localization and membrane fission. *B. subtilis ΔfisB* cells were partially complemented by *C. perfringens* FisB, despite only ∼23% identity between the two proteins, suggesting a common localization and membrane fission mechanism based on a few conserved biophysical properties. The membrane neck that eventually undergoes fission and where FisB accumulates is the most highly curved membrane region in the late stages of engulfment. Thus, FisB could potentially localize at the membrane neck due to a preference for highly curved membrane regions. We tested this possibility in experiments with both artificial giant unilamellar vesicles (GUVs) and live cells. Surprisingly, these experiments failed to reveal any intrinsic affinity of FisB for highly curved membranes. However, we found that FisB bridges membranes and accumulates at membrane adhesion sites. Using modeling, we found that self-oligomerization of FisB, coupled with its ability to bridge negatively charged membranes is sufficient to explain its localization to the membrane neck. Thus, proteins can localize to highly curved membrane regions through mechanisms independent of intrinsic curvature sensitivity. Together, these results suggest FisB-FisB and FisB-lipid interactions, combined with the unique membrane topology generated at the engulfment pole during sporulation, provide a simple mechanism to recruit FisB to mediate membrane fission independent of other factors.

## RESULTS

### Membrane fission occurs in the presence of a cluster of FisB molecules

To correlate FisB dynamics with membrane fission, we devised a labeling strategy that allowed us to monitor both simultaneously, using a modified version of a fission assay developed previously^41^. In this assay, synchronous sporulation is induced by placing *B. subtilis* cells in a nutrient-poor medium. At different time points after the nutrient downshift, aliquots are harvested from the culture, stained with the lipophilic membrane dye FM4-64, mounted on an agar pad, and imaged using fluorescence microscopy. The dye is virtually non-fluorescent in the medium, and it cannot cross the cell membrane.

Thus, before fission, FM4-64 labels the outer leaflet of both the mother cell and the forespore membranes. After fission, only the outer leaflet of the mother cell is labeled (S1 Appendix Figure 1B). Because post-fission cells and cells that never entered sporulation are labeled identically, in addition to FM4-64, a fluorescent protein is expressed in the forespore under the control of the forespore-specific transcription factor σ^F^ to distinguish between the two cell types^42^ (S1 Appendix Figure 1B). This makes it challenging to monitor FisB dynamics simultaneously, which requires a third channel. As an alternative, we used another lipophilic dye, TMA-DPH, that has partial access to internal membranes but can distinguish between pre- and post-fission stages without need for a forespore reporter^23^ (Figure 1C and S1 Appendix Fig. 1D-G). Using TMA-DPH as the fission reporter, we quantified the percentage of cells that have undergone fission as a function of time, for wild-type, *fisB* knock-out (*ΔfisB*, strain BDR1083, see S1 Appendix Table 2 for strains used), and *ΔfisB* cells complemented with FisB fused to monomeric EGFP (mGFP-FisB, strain BAM003) as shown in Figure 1D and 1F. These kinetic measurements reproduced results obtained using FM4-64 (S1 Appendix Figure 1C). Thus, TMA-DPH can be used as a faithful reporter of membrane fission, leaving a second channel for monitoring dynamics of FisB fused to a fluorescent reporter.

In the experiments of Figure 1D and 1F, we simultaneously monitored dynamics of mGFP-FisB and membrane fission. We found that membrane fission is almost always accompanied by an intense, immobile mGFP-FisB signal at the engulfment pole (Figure 1D, time= 3hr into sporulation). This intense <u>s</u>pot at the <u>e</u>ngulfment <u>p</u>ole (ISEP) is distinct from the dimmer, mobile clusters (DMC) that appear at earlier times elsewhere (Figure 1D). By 3 h into sporulation, around 70 % of the cells expressing mGFP-FisB at native levels had an ISEP (Figure 1G), a number that was close to the percentage of cells that had undergone fission by then (Figure 1F). Scoring individual cells, we found >90% (212/235) of cells that had undergone membrane fission also had an ISEP.

We also monitored membrane fission and mGFP-FisB signals in a strain with lower FisB expression. Here, lower FisB expression is achieved by reducing the spacing between the ribosome binding site (RBS) and the ATG start codon^43^. In this strain (BAL003), there was an initial delay in the fraction of cells that had undergone fission, but fission accelerated after t=3 h to reach near wild-type levels at around t=4h (Figure 1E,F). The fraction of cells with an ISEP followed a similar pattern (Figure 1G). The fraction of cells that had undergone fission at a given time was strongly correlated with the fraction of cells with an ISEP at that time (Figure 1H). Scoring individual cells, we found >93% (258/277) of cells that had undergone membrane fission had an ISEP. We conclude that membrane fission occurs in the presence of a large immobile cluster of FisB molecules at the site of fission.

### About 40 FisB molecules accumulate at the engulfment pole to mediate membrane fission

We asked how many copies of FisB are recruited to the engulfment pole at the time of membrane fission and how this number is affected by the expression level. For this quantification, we used DNA-origami based fluorescence standards we recently developed^44^. These standards consist of DNA rods (∼410 nm long and 7 nm wide) labeled with AF647 at both ends and a controlled number of mEGFP molecules along the rod (Figure 2A).

**Figure 2.**
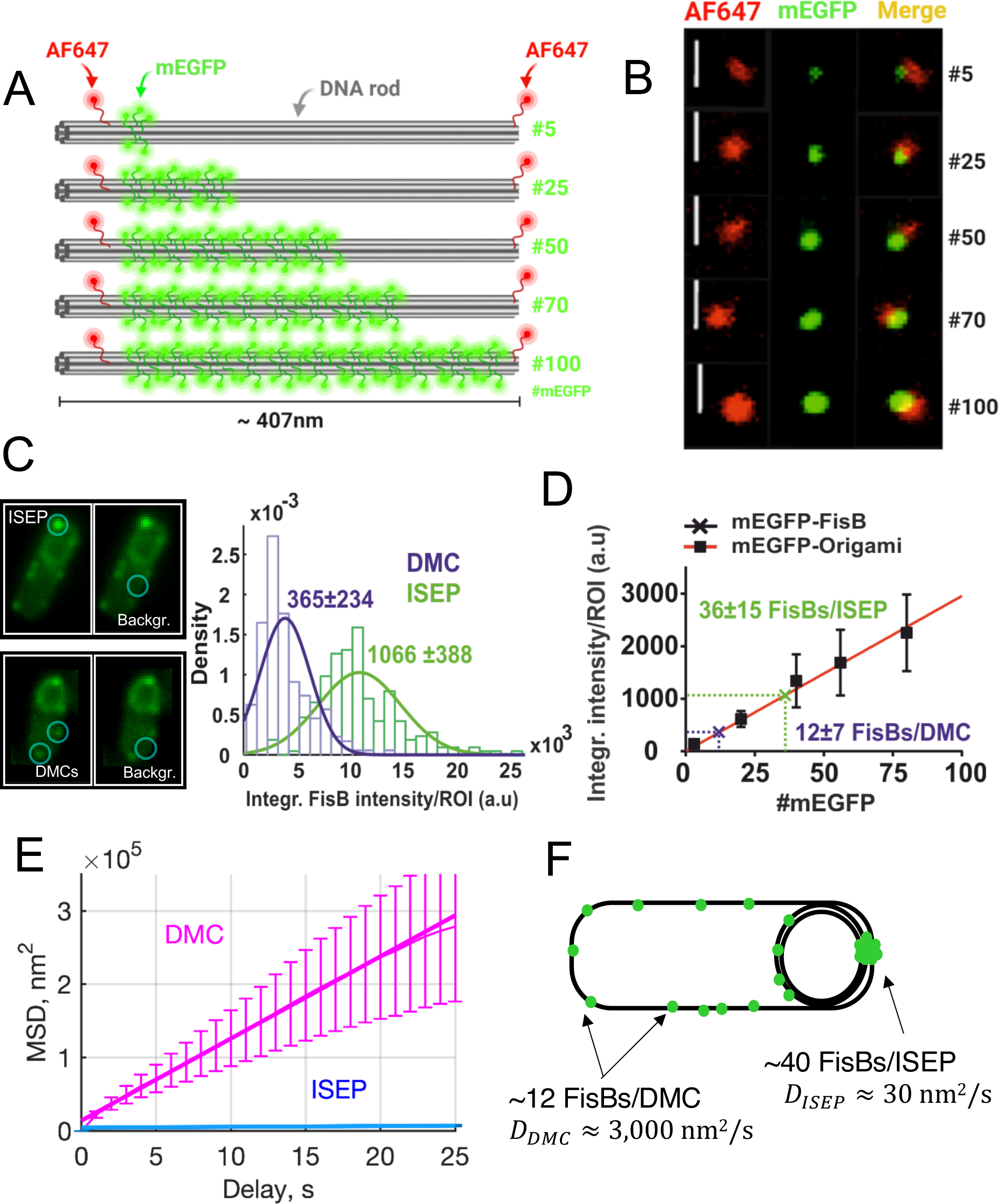
Estimation of mEGFP-FisB copies at the engulfment pole at t=3 h using DNA-origami calibration standards and mobility of FisB clusters. **A.** Simplified schematic of the DNA-origami-based mEGFP standards used in this study. Using DNA origami, DNA rods bearing AF647 at both ends and the indicated numbers of mEGFP molecules along the rod were designed. In the actual rods, the labeling efficiency was found to be ∼80%, so the actual copies of mEGFP per rod were 4, 20, 40, 56, and 80. **B.** Representative wide field images of the DNA-origami-based mEGFP standards used in this study. Bars are 1 μm. **C**. Distributions of total fluorescence intensities (sum of pixel values) for the intense spot at the engulfment pole (ISEP) and the dim, mobile clusters (DMC). Background was defined individually for every cell where an ISEP or DMC intensity measurement was performed. Examples are shown on the left. **D.** Total fluorescence intensity (sum of pixel values) for DNA-origami rods as a function of mEGFP copy numbers. The best fit line passing through the origin has slope 29.56 au/mEGFP (*R*^2^ = 0.97). The total intensity of the ISEP and DMCs correspond to ∼40 and ∼12 copies of mEGFP respectively. **E.** Mean-squared displacement (MSD) as a function of delay time for DMCs (magenta) and ISEPs (blue). Cells expressing mGFP-FisB (strain BAM003) were imaged using time-lapse microscopy. Forty-five cells from 10 different movies at t=2.5 hr and 30 cells from 10 different movies at t=3 hr after nutrient downshift were analyzed. (See S1 Appendix Movie 1 for a representative single bacterium at t=2.5 hrs showing several mobile DMCs and Movie 2 for a representative single bacterium at t=3 hrs showing an immobile ISEP.) Fits to the initial 25 s (∼10 % of delays) yielded *D_DMC_* = 2.80 ± 0.05 × 10^3^ nm^2^/s (± 95% confidence interval, R² = 0.999, 24 tracks) and *D_ISEP_* = 2.80 ± 0.51 × 10 nm^2/s (± 95% confidence interval, R² = 0.850, 25 tracks). **F.** Summary of FisB copy number and cluster mobility estimation.

DNA-origami standards carrying different mEGFP copies were imaged using widefield fluorescence microscopy (Figure 2B). For each type of rod, the average total fluorescence intensity of single-rods was computed and plotted against the number of mEGFP molecules per rod, generating the calibration curve in Figure 2D. We generated *B. subtilis* cells expressing mEGFP-FisB at native levels (BAL001) in a *ΔfisB* background so that images of these cells obtained under identical imaging conditions as for the calibration curve in Figure 2D could be used to compute mEGFP-FisB copy numbers. We imaged mEGFP-FisB cells at t=3 h after sporulation was induced. From these same images, we estimated the total fluorescence of dim, mobile clusters (DMC) and ISEP in *B. subtilis* cells as a sum of background-corrected pixel values (Figure 2C). Using the average values of these total intensities, we estimate ∼40 copies at the ISEP, and ∼12 per DMC from the calibration in Figure 2D. From the total intensity of cells (S1 Appendix Fig. 2E), we also estimate there are ∼1000 FisB molecules per cell. Two independent estimates, based on *B. subtilis* calibration strains^45^ and quantitative immunoblotting, resulted in slightly larger and smaller estimates of these copy numbers, respectively (S1 Appendix and S1 Appendix Figs. 2, 3).

**Figure 3.**
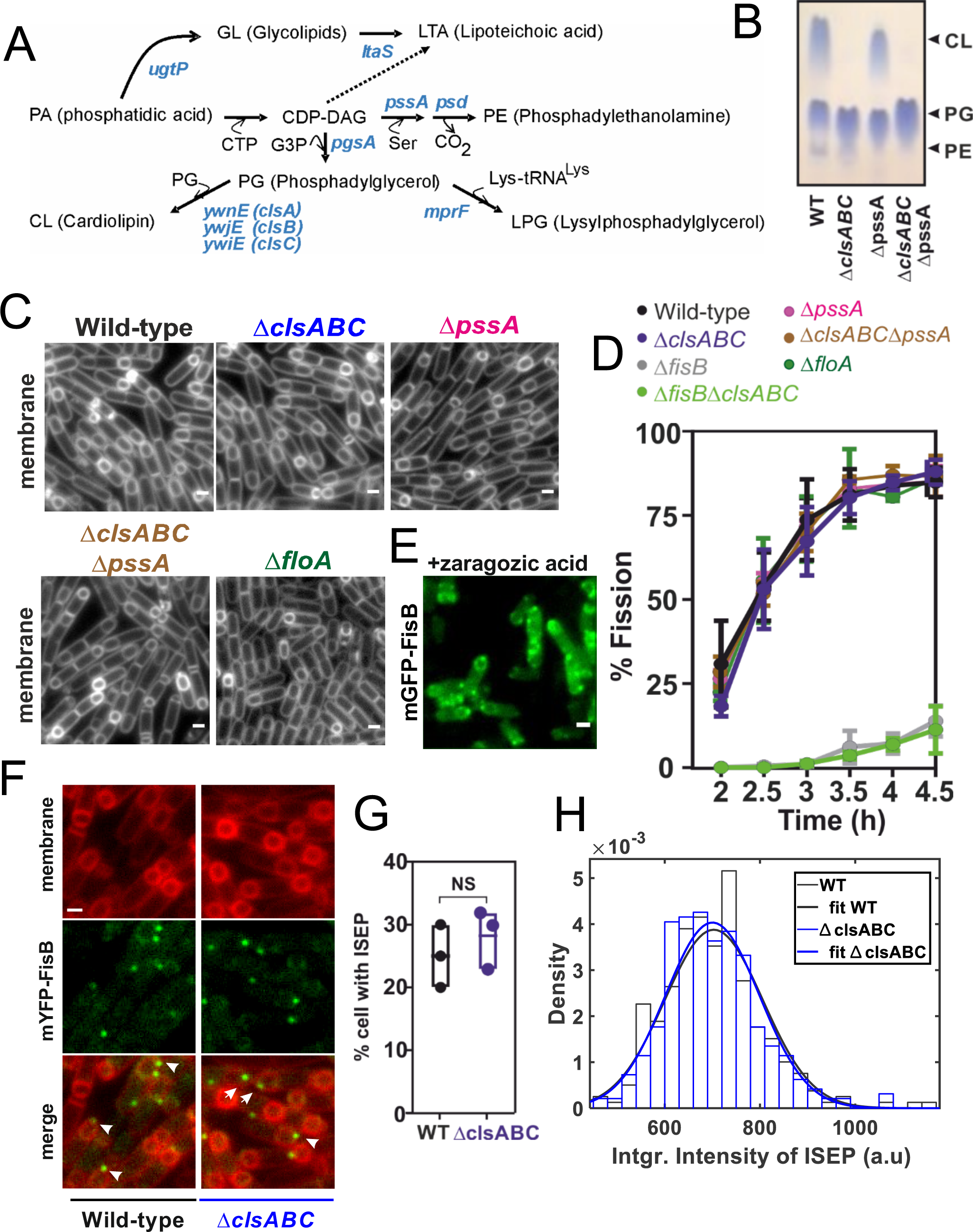
Membrane fission is insensitive to membrane lipid composition. **A.** Pathways for membrane lipid synthesis in *B. subtilis*. Lipid synthetases responsible for each step are highlighted in blue. **B.** Thin-layer chromatography (TLC) of the total lipid extracts of wild-type and indicated lipid synthesis-deficient cells. Cells were collected 3 hrs after induction of sporulation by nutrient downshift. Phospholipid spots (PLs) were visualized by staining with Molybdenum Blue spray reagent. Purified CL, PG, and PE were used as standards to identify the PLs of *B. subtilis*. Arrows indicate locations to which individual standards migrate. **C.** Membranes from cells of the indicated genetic backgrounds were visualized with TMA-DPH at t=3h. The images are from cells mounted on agarose pads containing sporulation medium. Bar,1 μm. **D.** Percentage of cells from indicated strains that have undergone membrane fission as a function of time after initiation of sporulation. For every strain, 150-220 cells from 3 independent experiments were analyzed at the indicated times during sporulation. **E.** mGFP-FisB (strain BAM003) treated with the squalene-synthase inhibitor zaragozic acid, imaged at t=3 h. **F.** Cells expressing mYFP-FisB (low expression levels) in either wild type (BAL002) or in a CL deficient strain (BAL037) at t=3h. Membranes were visualized with the fluorescent dye TMA-DPH. Examples of sporulating cells with a discrete mYFP-FisB focus at the cell pole (ISEP) are highlighted (white arrows). Foci were semi-automatically selected with SpeckletrackerJ^97^. **G.** The percentage of cells with an intense spot at engulfment pole for wild-type (BAL002) or cardiolipin-deficient (BAL037) mYFP-FisB expressing cells at t=3h (low expression). For each strain, 150-220 cells from 3 independent experiments were analyzed. **H.** Distributions of total fluorescence intensities (sum of pixel values) at ISEP for wild-type (BAL002) or cardiolipin-deficient (BAL037) mYFP-FisB cells at 3hr into sporulation. For every strain, 150 ISEPs were analyzed. Scale bars are 1 μm.

We tracked the DMC to estimate how rapidly they move. From the tracks, we calculated the mean squared displacement (MSD) as a function of time (Figure 2E). The short-time diffusion coefficient estimated from the MSD is *D*_*DMC*_ ≈ 2.8 × 10^3^ nm^2^/s (95% confidence interval CI=2.76 − 2.85 × 10^3^ nm^2^/s). This value is comparable to the diffusivity of FloA and FloT clusters of ∼100 nm with *D* ≈ 6.9 × 10^3^ and 4.1 × 10^3^ μm^2^/s, respectively^46^. By comparison, ISEP have *D*_*ISEP*_ ≈ 28 nm^2^/s (CI=22.9 − 33.1 nm^2^/s), two orders of magnitude smaller.

We performed similar estimations of FisB copy numbers for the low expression strain (BAL004) (S1 Appendix Fig. 4). We found ∼160±66, 122±51, or 83±6 (±SD) copies per cell using *B. subtilis* standards, DNA-origami, or the quantitative WB methods, respectively. For the ISEP, we found 8±2, 6±2, or 5±3 (±SD) copies of mGFP-FisB using the three approaches, respectively (S1 Appendix Table 1). About 6 % of the total mGFP-FisB signal accumulated in ISEP, close to the ∼4% in the native-expression strain (S1 Appendix Fig. 4E). The DMC were too dim to quantify reliably. Assuming DMCs to be ∼3-fold dimmer than ISEP like in the native-expression strain, each DMC would contain 2-3 mGFP-FisB, just below our detection limit. Interestingly, lowering the total expression of FisB per cell ∼8-fold resulted in a ∼6-fold reduction in the average number of FisB molecules found at the membrane fission site. Thus, only ∼6 copies of FisB are sufficient to mediate membrane fission, but only after some delay (Figure 1E,F).

**Figure 4.**
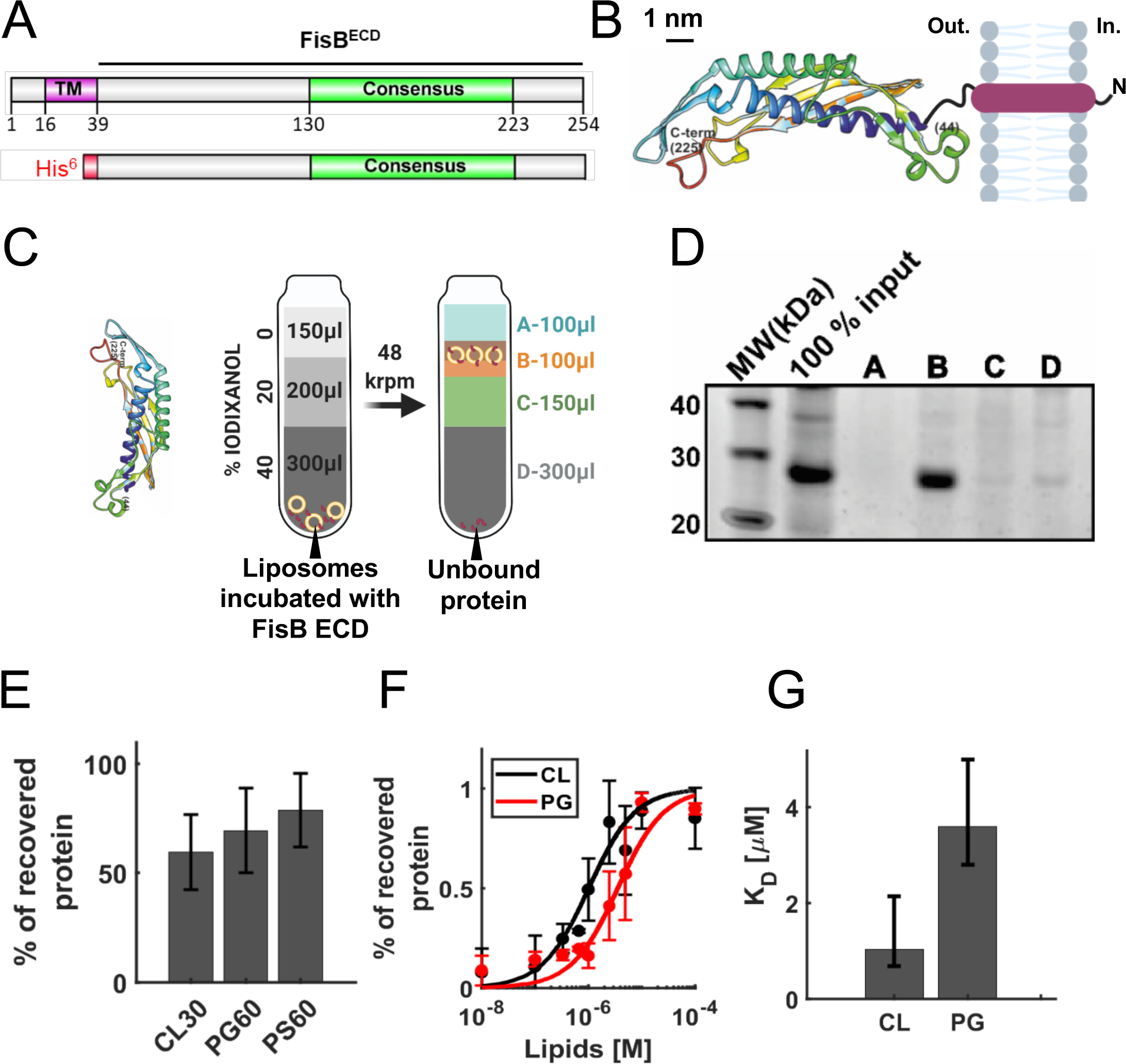
Binding of FisB ECD to acidic lipids. **A**. Domain structure of FisB and its His_6_-tagged extracytoplasmic domain (ECD) used in floatation experiments. **B.** Predicted model of FisB^44-225^ comprising most of the ECD^57^, schematically attached to the membrane. **C**. Schematic of the floatation assay. Liposomes (40 nmol total lipid) and FisB ECD (200 pmol) were incubated for 1 hour (total volume of 100 μl) at room temperature and layered at the bottom of an iodixanol density gradient. Upon ultracentrifugation, liposomes float to the top interface, whereas unbound protein remains at the bottom. Four fractions were collected as indicated and analyzed by SDS-PAGE. **D**. SYPRO orange stained gel of FisB ECD incubated with liposomes containing 45 mole % CL. The percentage of recovered protein is determined by comparing the intensity of the band in fraction B to the input band intensity. **E**. Indistinguishable amounts of FisB ECD are recovered when FisB ECD is incubated with liposomes containing different acidic lipid species as long as the charge density is similar. CL30, PG60, PS60 indicate liposomes containing 30 mole % CL, 60 mole % PG and 60 mole % PS, respectively. CL carries 2 negative charges, whereas PG and PS carry one each. The rest of the liposome composition is PC. **F**. Fraction of liposome-bound iFluor555-labeled FisB ECD (iFluor555-FisB ECD, 100 nM) recovered after floatation as a function of lipid concentration. Titration curves were fit to *f_b_* = *K*[*L*]/(1 + *K*[*L*]), where *f_b_* is the bound fraction of protein, [*L*] is the total lipid concentration (assumed to be ≫ [protein bound]), and *K* = 1/*K_d_* the apparent association constant, and *K_d_* is the apparent dissociation constant. **G**. Best fit values for *K*_*d*_ were 1.0 μM for CL (95% confidence interval, CI=0.7-2.1 μM) and 3.6 μM for PG (CI=2.8-5.0 μM), respectively. iFluor555-FisB ECD (100 nM) was incubated with10^-8^ to 10^-4^ M lipids for 1 h at room temperature before flotation. Liposomes contained 45 mole % of CL or PG and 55% PC.

In summary, ∼40 FisB molecules accumulate at the fission site to mediate membrane fission. Only 3-4 DMCs need to reach the fission site to provide the necessary numbers. When FisB expression is lowered ∼8-fold, ∼6 FisB molecules accumulate at the engulfment pole to mediate membrane fission, but fission takes longer.

### FisB localization and membrane fission are independent of cardiolipin, phosphatidylethanolamine and flotillins

To investigate how FisB is recruited to the membrane fission site, we began by testing a potential role for the cell wall remodeling machinery, the protonmotive force, and the membrane potential, and found none influenced FisB dynamics (S1 Appendix Results and S1 Appendix Fig. 6).

**Figure 6.**
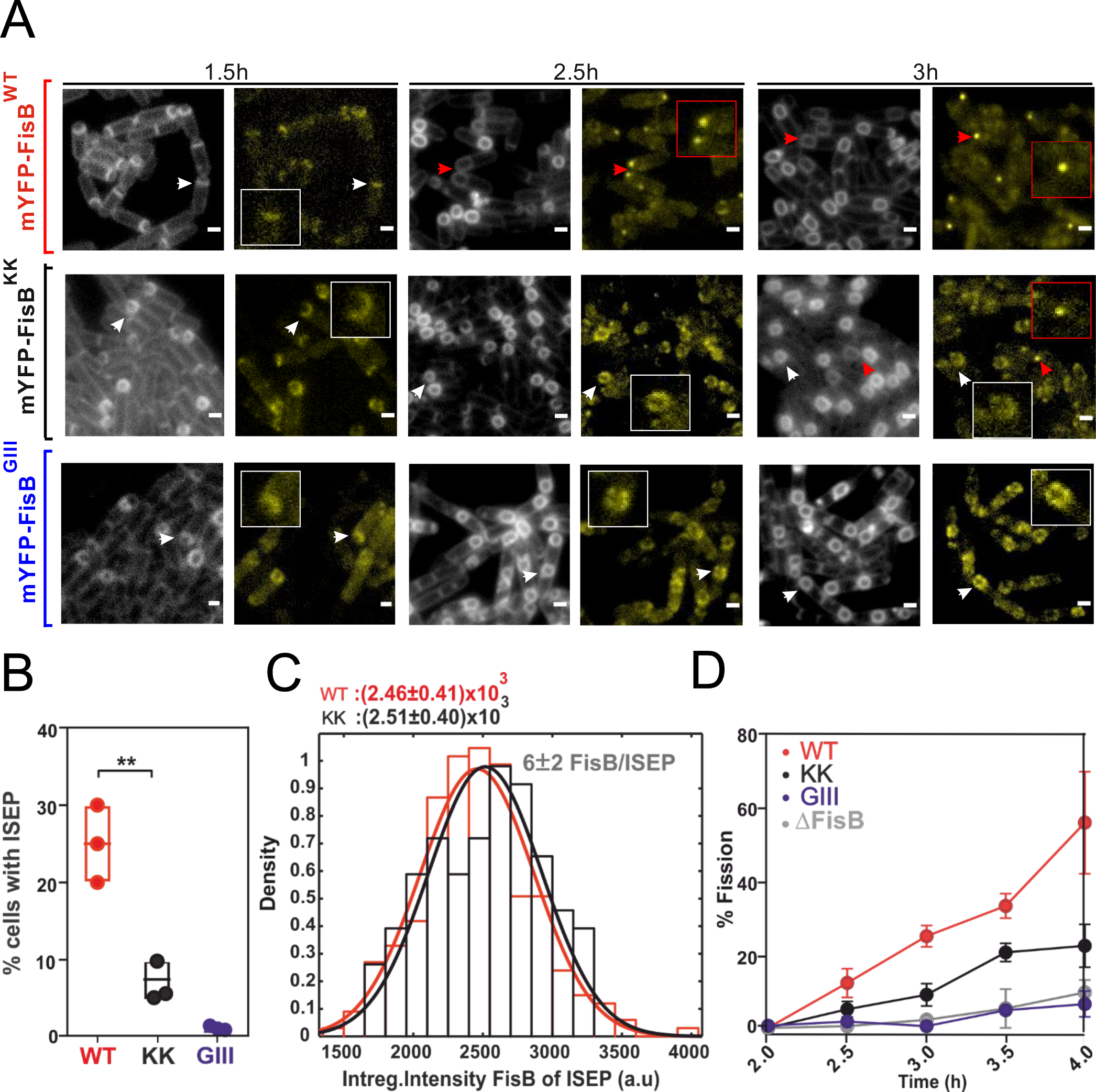
FisB clustering and binding to acidic lipids are both required for ISEP formation and membrane fission. **A.** Snapshots of sporulating *ΔfisB* cells expressing mYPF-FisB^WT^ (BAL002), mYPF-FisB^KK^ (BAL006), or mYPF-FisB^GIII^ (BAL007), at low levels. For each time point after downshifting to the sporulation medium, cell membranes were labeled with TMA-DPH and images were taken both in the membrane (left) and the YFP (right) channels. By t=2.5 h, some foci at the engulfment pole (ISEP) are visible for WT cells that have undergone membrane fission (red boxes), but not for the KK or GIII mutants (white boxes). A small fraction of KK mutants (7.3%) accumulated FisB at the engulfment pole and underwent membrane fission at t=3h. Scale bars represent 1 μm. **B.** Percentage of cells with an intense spot at the engulfment membrane (ISEP) at t=3 h into sporulation, for WT FisB, FisB^KK^, or FisB^GIII^. For every strain, 200-300 cells from three independent experiments were analyzed at the indicated times during sporulation. **C**. Distribution of background-corrected integrated intensities (sum of pixel values) of ISEP fluorescence for *ΔfisB* cells expressing mYFP-FisB^WT^ or mYPF-FisB^KK^. The distributions are indistinguishable. Since low-expression cells accumulate, on average, 6±2 FisB^WT^ molecules at the ISEP (Fig. S4D), so do FisB^KK^ cells. 175 and 68 ISEPs were analyzed for WT and KK mutant strains. **D**. Percentage of cells that have undergone membrane fission at the indicated time points. (For every strain, 200-300 cells from 3 independent experiments were analyzed at the indicated times during sporulation.)

**Figure 5.**
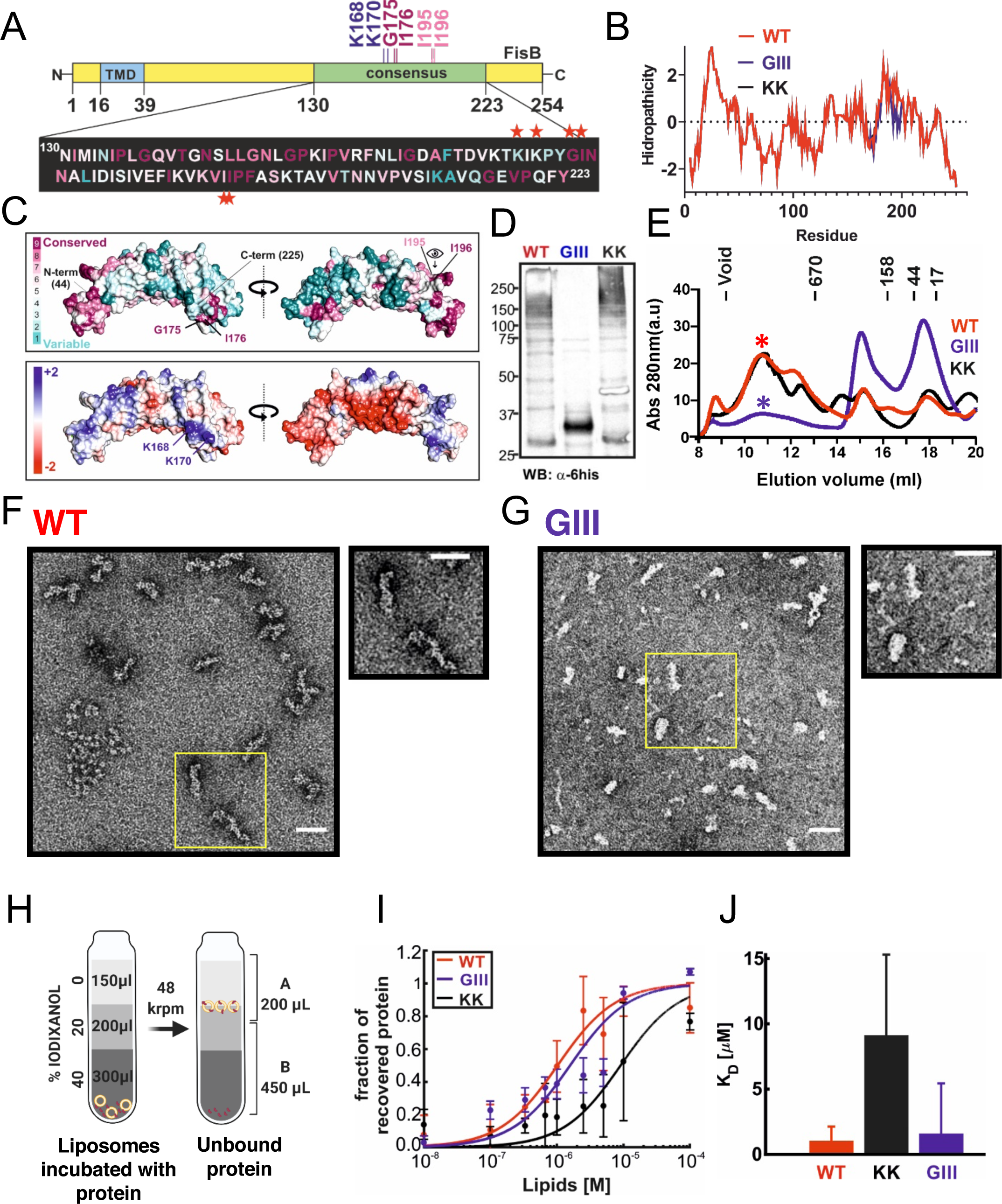
FisB mutants selectively impaired in oligomerization and membrane binding. **A**. Mutated residues shown on the FisB domain structure. **B**. Kyle-Doolittle hydrophobicity profile of the FisB sequence for wild-type (WT), FisB K168D,K170E (FisB^KK^), and FisB G175A,I176S, I195T, I196S (FisB^GIII^) mutants. **C.** Mutations shown on the predicted model^57^ of FisB^44-225^. Residue conservation (top) and electrostatic potential (bottom) are mapped onto the structure. **D**. Western blot of cell lysates from *E. coli* cells expressing FisB-ECD^WT^, FisB-ECD^GIII^, or FisB-ECD^KK^, probed with an anti-histidine antibody. High molecular weight bands in the WT and KK lanes are largely absent in the GIII lane, indicating FisB^GIII^ is less prone to forming oligomers. **E**. Size-exclusion chromatography of FisB WT and the GIII and KK mutants. Intensities of high and low molecular weight peaks are reversed for FisB WT and the GIII mutant, whereas the KK mutant has a profile similar to WT. **F**. A fraction corresponding to the high-molecular peak in E (indicated by *) for FisB WT was collected and imaged using negative-stain electron microscopy (EM), which revealed flexible, elongated structures ∼50 nm × 10 nm. **G**. A similar analysis for FisB^GIII^ revealed more heterogeneous and less stable structures. Scale bars in F, G are 50 nm. **H**. Schematic of the floatation experiments to determine the apparent affinity of FisB mutants for liposomes containing acidic lipids. Experiments and analyses were carried out as in Figure 4, except only two fractions were collected. iFluor555-FisB ECD (100 nM) was incubated with10^-8^ to 10^-4^ M lipids for 1 h at room temperature before floatation. Liposomes contained 45 mole % of CL and 55% PC. **I**. Fraction of protein bound to liposomes as a function of total lipid concentration. Data was fitted to a model as in Figure 4F. The data and fit for FisB WT is copied from Figure 4F for comparison. **J.** Best fit values for *K*_*d*_ were 1.0 μM for WT (95% confidence interval, CI=0.7-2.1 μM), 9.1 μM for KK (CI=6.5-15.3 μM), and 1.6 for GIII (CI=0.9-5.1 μM), respectively.

We then tested whether lipid microdomains play a role in recruitment to the site of fission. Previously, we reported that the recombinant, purified extracytoplasmic domain (ECD, see Figure 4A) of FisB interacts with artificial lipid bilayers containing CL^23^. To test if FisB-CL interactions could be important for the subcellular localization of FisB and membrane fission, we generated a strain (BAM234) that carries deletions of the three known CL synthase genes *ywnE (clsA)*, *ywjE (clsB)* and *ywiE (clsC)*^47^ (Figure 3A). The CL synthase-deficient strain did not contain detectable levels of CL at t=3 hours after sporulation was initiated (Figure 3B). CL-deficient cells grew normally but had a reduction in sporulation efficiency as assayed by heat-resistant (20 min at 80°C) colony forming units (S1 Appendix Table 2 and S1 Appendix Figure 5)^33^. A reduction in sporulation efficiency measured in this manner can be due to a defect at one or several steps during sporulation or germination. Importantly, the membrane fission time course of *ΔclsABC* cells was indistinguishable from those of wild-type cells (Figure 3C,D), indicating the defect in sporulation occurs at a stage after membrane fission. In addition, mYFP-FisB localization and dynamics were similar in *ΔclsABC* (BAL037) and wild-type (BAL002) cells (Figure 3F-H). The fraction of cells that had an ISEP, and the intensity of the ISEP, reflecting the number of FisB molecules recruited to the membrane fission site, were indistinguishable for wild-type and *ΔclsABC* cells (Figure 3G,H). We conclude that CL is not required for the subcellular localization of FisB or membrane fission.

Next, we tested a potential role for phosphatidylethanolamine (PE), another lipid implicated in membrane fusion and fission^48,49^ and that forms microdomains^50^. We deleted the *pssA* gene which encodes phosphatidylserine synthase that mediates the first step in PE synthesis (Figure 3A) to generate cells lacking PE (strain BAL031, Figure 3B). Kinetics of membrane fission during sporulation were identical in *ΔpssA* and wild-type cells (Figure 3D), indicating PE does not play a significant role in membrane fission.

PE and CL domains in *B. subtilis* membranes tend to occur in the same sub-cellular regions^50^, raising the possibility that CL and PE may compensate for each other. To test whether removing both CL and PE affects fission, we generated a quadruple mutant (BAL030) lacking both CL and PE (Figure 3B), leaving PG as the major phospholipid component of the membrane. Surprisingly, the quadruple mutant underwent fission with indistinguishable kinetics compared to wild-type (Figure 3C,D). Thus, two lipids with negative spontaneous curvature and implicated in membrane fusion and fission reactions in diverse contexts have no significant role in membrane fission mediated by FisB during sporulation.

In addition to CL and PE microdomains, bacteria also organize many signal transduction cascades and protein-protein interactions into functional membrane microdomains (FMMs), loose analogs of lipid rafts found in eukaryotic cells^40^. The FMMs of *B. subtilis* are enriched in polyisoprenoid lipids and contain flotillin-like proteins, FloT and FloA, that form mobile foci in the plasma membrane^51,52^. FloT-deficient cells have a sporulation defect, but which sporulation stage is impaired is not known^46^. We observed that *ΔfloA* (BAL035), but not *ΔfloT* (BAL036), cells are impaired in sporulation as assayed by heat-resistant colony forming units (S1 Appendix Table 2, and S1 Appendix Figure 5). However, when we monitored engulfment and membrane fission, we found both proceeded normally in *ΔfloA* cells (Figure 3D). Thus, the sporulation defect in *ΔfloA* cells lies downstream of engulfment and membrane fission. This was confirmed by blocking formation of FMMs during sporulation by addition of 50 μM zaragozic acid^53^ to the sporulation medium which had no effect on the localization of mGPF-FisB (Figure 3E).

Together, these results imply that FisB-mediated membrane fission that marks the end of engulfment during sporulation is insensitive to the negative-curvature lipids CL, PE, and to FloA/T-dependent lipid domains.

### FisB binds to acidic lipids

PG can substitute for CL as a binding partner for many proteins^54,55^. To see if this might also be the case for the FisB ECD, we quantified the affinity of this domain for both lipids.

Most, but not all, algorithms (S1 Appendix Fig. 7) predict FisB to possess a single transmembrane domain (TMD) with a small N-terminal cytoplasmic domain and a larger (23-kDa) ECD, as depicted in Figure 4A. We first confirmed this predicted topology using a cysteine accessibility assay^56^ (S1 Appendix Fig. 8, Materials and Methods, and S1 Appendix Results). Our attempts to determine the structure of recombinant, purified FisB ECD were unsuccessful, but a computational model of FisB for residues 44 to 225, covering most of the ECD is available^57^ and is shown in Figure 4B. The model predicts a curved ECD structure, with ∼3 nm and ∼5 nm for the inner and outer radii of curvatures. The overall topology of FisB, with the predicted ECD structure is depicted in Figure 4B.

**Figure 7.**
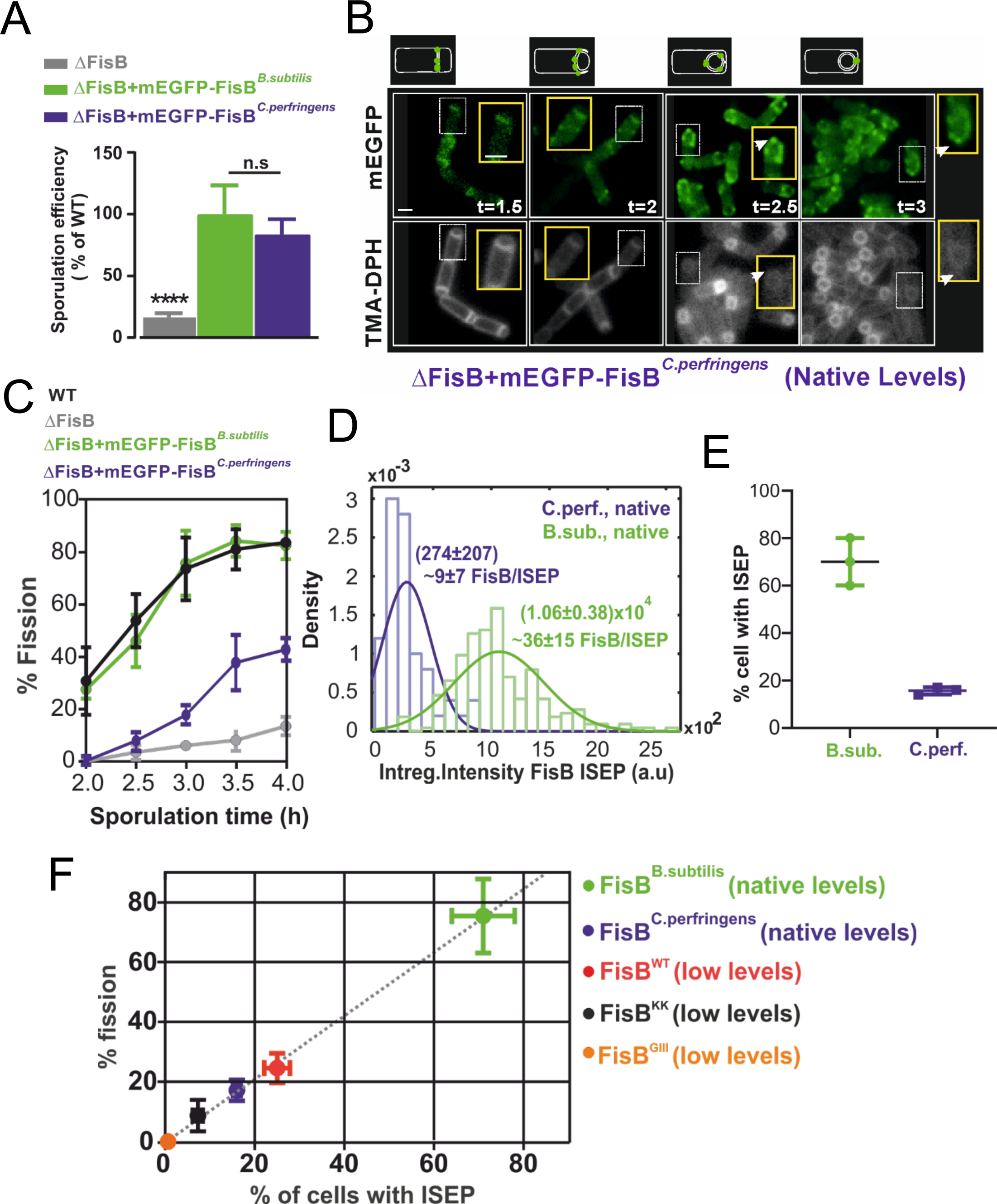
*C. perfringens* FisB can substitute for *B. subtilis* FisB despite poor sequence identity. **A.** Heat-resistant colony forming units for *ΔfisB* cells expressing *B. subtilis* (BAL001) or *C.perfringens* FisB (BAL005) at native levels, presented as a percentage of the WT sporulation efficiency. Results are shown as means ± SD for three replicates per condition. **B.** Snapshot of *ΔfisB* cells expressing mEGFP-FisB*^Cperfringens^*. Aliquots were removed at the indicated times, membranes labeled with TMA-DPH, and both the TMA-DPH and the EGFP channels imaged after mounting into agar pads. White boxed areas are shown on an expanded scale in yellow boxes. Arrows indicate cells with ISEP that have undergone membrane fission. Bar, 1 μm. **C**. Percentage of cells that have undergone membrane fission as a function of sporulation time for wild-type cells, *ΔfisB* cells, *ΔfisB* cells expressing *B. subtilis* mEGFP-FisB at native levels, or *ΔfisB* cells expressing mEGFP-FisB*^Cperfringens^*. The plots for the first three conditions are reproduced from Figure 1F for comparison. **D.** Distribution of background-corrected total fluorescence intensity of ISEP for *ΔfisB* cells expressing mEGFP-FisB*^Cperfringens^* or mEGFP-FisB*^Bsubtilis^* at native levels. From the calibration in Figure 2D, we estimate 9±7 FisB*^Cperfringens^* per ISEP. The distribution for mEGFP-FisB*^Bsubtilis^* is reproduced from Figure 2C for comparison. (150 and 93 ISEPs were analyzed for mEGFP-FisB*^Bsubtilis^* and mEGFP-FisB*^Cperfringens^*, respectively.) **E**. Percentage of cells with ISEP, for *ΔfisB* cells expressing mEGFP-FisB*^Cperfringens^* or mEGFP-FisB*^Bsubtilis^*. (For each strain, 200-300 cells from 3 independent experiments were analyzed.) **F**. Percentage of cells that have undergone membrane fission at t=3 h vs. the percentage of cells with ISEP at the same time point, for the conditions indicated. There is a nearly perfect correlation between these two quantities (the dashed line is a best-fit, *y* = 1.03*x*, *R*^2^ = 0.96).

**Figure 8.**
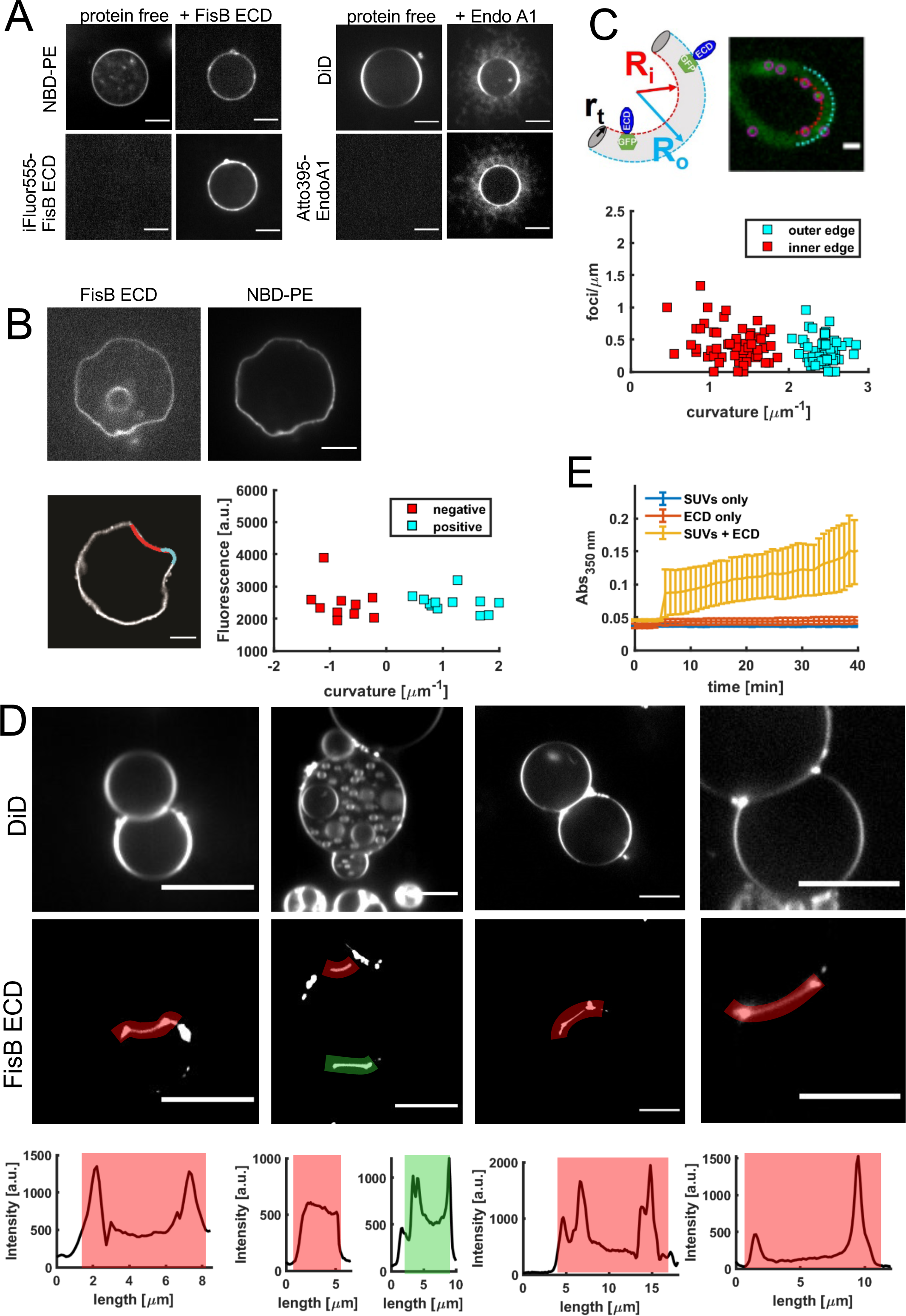
FisB does not sense or induce membrane curvature. **A.** FisB ECD does not induce deformation of GUV membranes. Left: GUVs incubated with 2 μM iFluor555-labeled FisB ECD did not show any tubulation or invagination of the GUV membrane. GUVs were composed of (in mole %: 25 *E. coli* PE, 5 *E. coli* CL, 50 *E. coli* PG, 19 eggPC, 1 NBD-PE). Right: incubation of 2 μM endophilin A1 (EndoA1, labeled with Atto395) with GUVs (45% DOPS, 24.5% DOPC, 30% DOPE and 0.5% DiD) resulted in extensive tubulation of membranes, as reported previously^22^. The two proteins have similar affinities for GUV membranes under these conditions (Figure 4F and ref. 106). **B**. FisB ECD cannot deform deflated GUVs and its membrane localization is independent of curvature. To avoid potential issues with high membrane tension preventing membrane deformation, GUVs were deflated using osmotic stress, which resulted in deformed GUVs with both negatively and positively curved regions. FisB ECD bound to these GUVs was unable to induce any high-curvature deformations. The intensity of iFluor555-FisB ECD along a membrane contour (proportional to coverage) was plotted against membrane curvature in the corresponding region. There was no correlation between membrane curvature and FisB ECD coverage. **C**. FisB localization does not depend on curvature in filamentous *B. subtilis* cells. GFP-FisB was expressed under an inducible promoter during vegetative growth and cell division was blocked by inducing expression of MciZ^68^. Cells grew into long flexible filaments that were bent to varying degrees. The linear density of GFP-FisB spots (spots/μm) was independent of filament curvature. **D**. FisB ECD bridges GUV membranes. iFLuor555-FisB ECD (100 nM) was incubated with GUVs (same composition as in A and B). Many GUVs were found adhering to one another. iFluor555-FisB ECD signals were enhanced in the adhesion patches, in particular at the rims. Intensity profiles along the highlighted contours are shown below the examples. **E**. FisB ECD aggregates small liposomes. Liposomes (in mole %: 25 *E. coli* PE, 5 *E. coli* CL, 50 *E. coli* PG, 19 eggPC, 50 μM total lipid) were incubated in the absence and presence of FisB ECD (unlabeled) and their aggregation monitored by absorbance at 350 nm. FisB was added at 5 min (1 μM final), which caused the absorbance to increase, indicating increased liposome aggregation.

We probed interactions of FisB ECD with PG using a liposome co-flotation assay, illustrated in Figure 4C. Purified recombinant, soluble FisB ECD (Fig. 4A, bottom) was incubated with liposomes and subsequently layered at the bottom of a discontinuous density gradient. Upon equilibrium ultracentrifugation, the lighter liposomes float up to the interface between the two lowest density layers together with bound protein, while unbound protein remains at the bottom of the gradient. We collected fractions and determined the percentage of protein co-floated with liposomes using SDS-PAGE and densitometry, as shown in Figure 4D. We first determined that binding of FisB ECD to liposomes containing CL was not dependent on pH or the divalent ion Ca^2+^ (S1 Appendix Fig. 9F,G). By contrast, the fraction of liposome-bound protein decreased rapidly as the ionic strength increased (S1 Appendix Fig. 9H). These results indicated binding was mainly electrostatic in nature.

**Figure 9.**
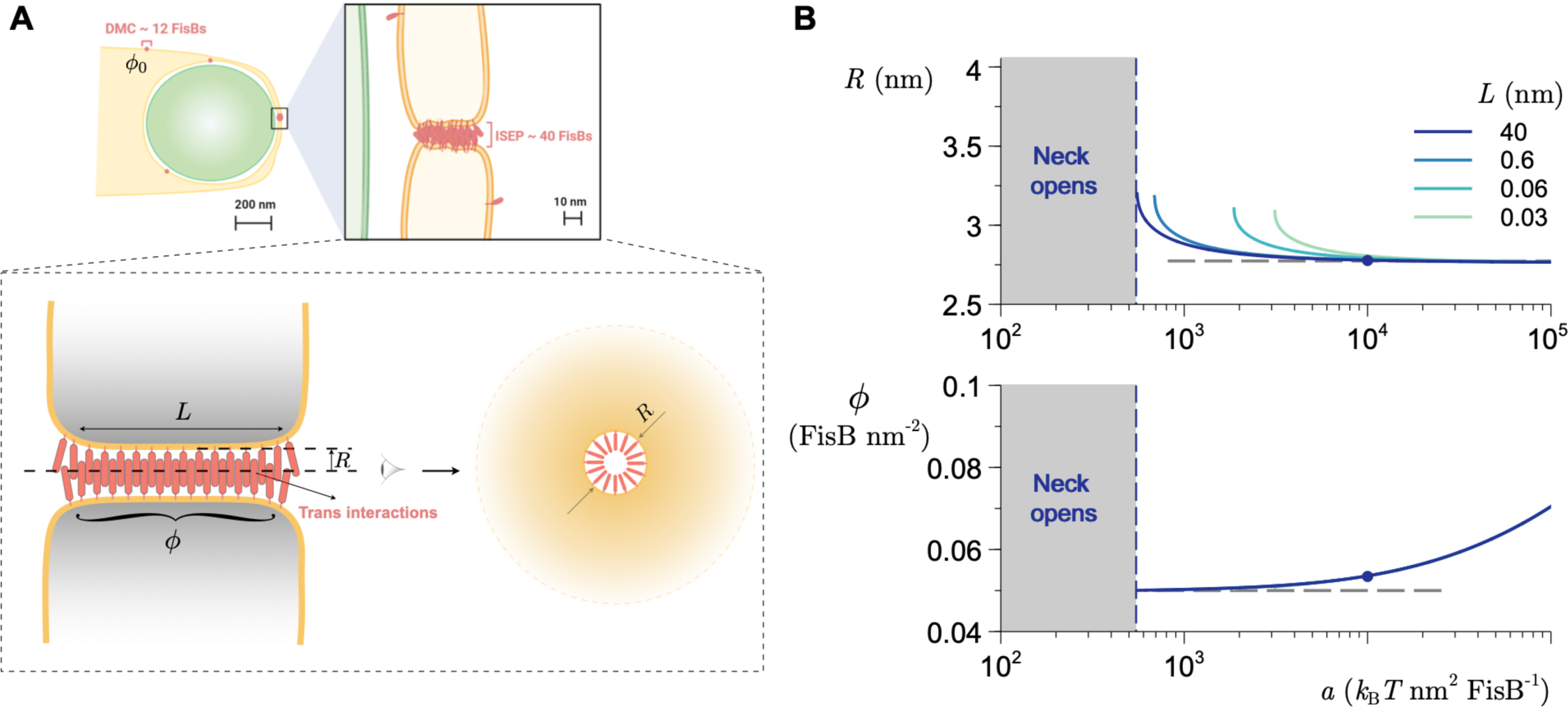
Modeling supports recruitment of FisB to membrane neck via oligomerization without curvature sensing. **A.** Left: Schematic of the late stages of engulfment, when a small membrane neck connects the engulfment membrane to the rest of the mother cell membrane. Right: Schematic of FisB accumulation at the fission site. FisB freely moves around the engulfment membrane and other regions of the mother cell membrane, forming clusters of up to ∼12 molecules. Cluster motions are independent of lipid microdomains, flotillins, the cell-wall synthesis machinery, and voltage or pH gradients. About 40 copies of FisB accumulate at the membrane neck in an immobile cluster. Bottom: Modeled axisymmetric membrane neck of radius *R* and length *L* connecting two membrane sheets. The uniform areal concentration of FisB in the neck is *ɸ*. **B.** Top: Equilibrium radius of the neck as a function of FisB trans homo-oligomerization strength, *a*, for several values of neck length, *L*. Below a minimum interaction strength, FisB cannot stabilize the neck and the neck opens. The horizontal line is the radius corresponding to the minimum of the potential describing the trans interaction, *R* = 2^1/6^σ (Eq. 3). Bottom: Equilibrium FisB concentration in the neck as a function of *a*. The horizontal line is 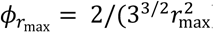), the concentration of FisB at the onset of in-plane crowding. Model parameters (see Eqs. 5 and 6): *κ* = 20 *k*_B_*T* (ref. 107), *ɸ*_0_ = 100 FisB *μ*m^-2^, *γ* = 10^-4^ N m^-2^ (ref. ^108^), σ_cis_ ≃ 2.47nm, *ɸ*_max_= 5 × 10^4^ FisB *μ*m^-2^, *L* = 40 nm, and for the dot *a* ≃ 10^4^ *k*_B_*T* nm^2^ FisB^-1^. For details, see S2 Appendix.

At neutral pH, CL carries two negative charges, whereas PG and phosphatidylserine (PS), a lipid not normally found in *B. subtilis*^58^, carry only a single negative charge. If binding is mediated mainly by electrostatic interactions, then liposomes carrying PG or PS at two times the mole fraction of CL should bind nearly the same amount of FisB ECD, since the surface charge density would be the same. Indeed, similar amounts of FisB ECD were bound to liposomes carrying 30% CL, 60% PG, or 60% PS (Figure 4E). FisB ECD did not bind neutral phosphatidylcholine PC liposomes^23^.

To quantify the affinity of recombinant soluble FisB ECD for CL vs PG, we then titrated liposomes containing 45 mole % CL or PG and measured binding of 100 nM FisB ECD (Figure 4F). In these experiments, we used iFluor555 labeled FisB ECD (iFluor555-FisB ECD) and detected liposome-bound protein using fluorescence rather than densitometry of SYPRO-stained gels, which extended sensitivity to much lower protein concentrations. The titration data were fit to a model to estimate the apparent dissociation constant, *K*_*d*_ (see Materials and Methods), which was 1.0 μM for CL (95% confidence interval CI=0.7-2.1 μM) and 3.6 μM for PG, respectively (CI=2.8-5.0, Figure 4F,G).

Together, these results suggest that while FisB has higher affinity for CL than for PG, the higher affinity results mainly from the higher charge carried by CL. FisB does not bind CL with much specificity; at the same surface charge density, FisB ECD binds PG, or even PS which is not a *B. subtilis* lipid, with similar affinity. Thus, *in vivo* FisB is likely to bind CL as well as PG which is much more abundant.

### Purified FisB ECD forms soluble oligomers

FisB forms clusters of various sizes in cells as described above (Figure 1, 2) and does not appear to have other protein interaction partners^23^. Thus, homo-oligomerization of FisB may be important for its function. We explored oligomerization of recombinant, soluble FisB ECD (Figure 5). When FisB ECD bearing a hexa-histidine tag is expressed in *E*. *coli* and purified to homogeneity by affinity chromatography, samples analyzed by SDS-PAGE show multiple bands corresponding to different oligomeric states (Figure 5D and S1 Appendix Figure 9B). Size-exclusion chromatography (SEC) analysis resolved the purified protein into predominant high molecular weight oligomeric structures eluting over a wide range of sizes, and low molecular weight peaks comprising minor components (Figure 5E and S1 Appendix Figure 9C, top). The minor peak at ∼23 kDa (18 ml elution volume) corresponds to monomeric FisB ECD, whereas the peak at ∼400 kDa (15 ml) is FisB ECD that co-elutes with another protein, likely the 60 kDa chaperone GroEL, a common contaminant in recombinant proteins purified from *E. coli* (S1 Appendix Figure 9D). To rule out potential artefacts caused by the hexa-histidine affinity tag, we also purified FisB ECD using a GST-tag, which yielded similar results.

The SEC of high molecular weight peaks collected from the initial chromatogram did not show a redistribution when re-analyzed (S1 Appendix Figure 9C, bottom), suggesting that once formed, the oligomeric structures are stable for an hour or longer. We analyzed the high molecular-weight SEC fractions (peaks 1 and 2) using electron microscopy (EM) after negative staining. This analysis revealed rod-like structures quite homogeneous in size, ∼50 nm long and ∼10 nm wide (Figure 5F and S1 Appendix Figure 9E). These structures displayed conformational flexibility, which precluded structural analysis using cryoEM (and likely hampered our attempts to crystallize FisB ECD). We estimate every rod-like oligomer can accommodate ∼40 copies of the predicted structure of FisB^44-225^ shown in Figure 4B, similar to the number of FisB molecules recruited to the membrane fission site in cells (Figure 2).

### A FisB mutant that is selectively impaired in homo-oligomerization

To determine whether self-oligomerization and lipid-binding interactions are important for FisB’s function, we generated a series of mutants, characterized oligomerization and lipid-binding of the mutant proteins *in vitro*, and analyzed FisB localization dynamics and membrane fission during sporulation *in vivo*.

We suspected self-oligomerization of FisB was at least partially due to hydrophobic interactions. Accordingly, we first mutated conserved residues G175, I176, I195 and I196 in a highly hydrophobic region of FisB ECD (Figure 5A,B), producing a quadruple mutant, G175A,I176S, I195T, I196S (FisB^GIII^). These residues are on the surface of the predicted structure of FisB ECD (Figure 5C), so are not expected to interfere with folding. Purified FisB^GIII^ ECD displayed reduced oligomerization when analyzed using SDS PAGE or size exclusion chromatography (Figure 5D,E). Though much reduced in amplitude, a broad, high molecular weight peak was still present in size exclusion chromatograms (Figure 5E). Negative-stain EM analysis of this fraction revealed oligomerization with less defined size and structure compared to wild type FisB ECD (Figure 5G).

To test whether lipid binding of the GIII mutant was affected, we used the co-flotation assay described above, except only two fractions were collected (Figure 5H,I). This analysis revealed that, despite being impaired in self-oligomerization, FisB^GIII^ ECD has lipid binding properties similar to wild-type with a dissociation constant 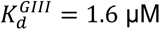 (95% confidence interval CI=0.9-5.1 μM), indistinguishable from that of wild type FisB ECD^WT^ (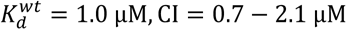, Figure 5J).

### FisB^K168D,K170E^ (FisB^KK^) is selectively impaired in binding acidic lipids

To engineer lipid-binding mutants, we took advantage of our observation that FisB binding to anionic lipids is principally mediated through electrostatic interactions (S1 Appendix Figure 9H). We generated a series of mutants in which we either neutralized or inverted up to four charges (S1 Appendix Fig. 11 and S1 Appendix Table 2). The ECD of a set of charge neutralization mutants were expressed in *E. coli*, purified and tested for lipid binding using the liposome co-floatation assay. The largest reductions in lipid binding were observed when lysines in a region comprising residues 168-172 were neutralized (S1 Appendix Fig. 11A). This region corresponds to a highly positively charged pocket in the predicted model of FisB 44-225 (Figure 5C).

A partially overlapping set of FisB mutants were expressed in a *ΔfisB* background and tested for sporulation efficiency by monitoring formation of heat-resistant colonies (S1 Appendix Fig. 11B-E). Again, the strongest reductions in sporulation efficiency were found when lysines 168, 170 or 172 were mutated (S1 Appendix Fig. 11D). We decided to characterize the K168D, K170E mutation in more detail, as it produced the strongest reduction in sporulation efficiency.

We purified the ECD of FisB^K168D,K170E^ (FisB^KK^) from *E. coli* and tested its binding to liposomes containing 45 mole % CL using the co-floatation assay (Figure 5H-J). The dissociation constant for FisB^KK^-acidic lipid binding was 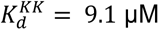 (CI=6.5-15.3 μM), nearly 10-fold lower than that for wild-type FisB ECD (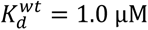, CI = 0.7 −2.1 µM, Figure 5I,J). Importantly, formation of oligomers was not affected (Figure 5D,E). Thus, FisB^KK^ is specifically impaired in binding to acidic lipids.

### FisB-lipid interactions and homo-oligomerization are important for targeting FisB to the fission site

Using the FisB mutants selectively impaired in binding to lipids or homo-oligomerization, we investigated whether these activities are important for FisB’s function *in vivo*. To analyze FisB clustering and targeting to the fission site, we fused wild-type FisB or the two mutants to an N-terminal monomeric YFP (mYFP) and expressed the fusions at lower levels, which facilitated observation of ISEPs (Figure 6A). We induced these strains to sporulate and monitored FisB dynamics and membrane fission. Both the lipid-binding (FisB^KK^) and the oligomerization mutant (FisB^GIII^) were targeted to the cell membrane, unlike many other mutants we tested (S1 Appendix Figure 11E and S1 Appendix Table 2). At t=1.5 h after the nutrient downshift, mYFP-FisB signals were visible in all strains without any distinguishing features. At t=2.5 h, a subset of cells expressing the wild-type FisB fusion had undergone membrane fission and these cells had an ISEP. By contrast, membrane fission was not evident in either of the mutants. By t=3 h, 25% of WT FisB cells had undergone fission, nearly always with an accompanying ISEP. In the lipid binding FisB^KK^ mutant, only 8% of the sporulating cells had accomplished membrane fission (Figure 6B), but more than 90% of those that did had an ISEP (53/58 cells). Membrane fission events and the accompanying bright mYFP-FisB spots were very rare (0.6%) in the oligomerization-deficient FisB^GIII^ mutant.

The distribution of fluorescence intensities of the foci from low-expression WT and KK cells were indistinguishable (Figure 6C). Using the DNA-origami fluorescence intensity calibration (Figure 2), we estimate 6±2 copies of low-expression FisB WT or the KK mutant to have accumulated at the fission site. For the GIII mutant, there were not enough cells with an intense spot to perform a similar analysis.

From TMA-DPH labeling, we determined the fraction of cells that successfully completed fission as a function of time (Figure 6D). Oligomerization-deficient FisB^GIII^ was not able to induce fission, whereas the lipid-binding mutant FisB^KK^ had a partial, but severe defect (∼50% reduction compared to wild-type). Importantly, both mutants were expressed at levels similar to the wild-type (S1 Appendix Fig. 10), so the defects to form an ISEP and undergo membrane fission are not due to lower expression levels.

Together, these results suggest FisB-lipid and FisB-FisB interactions are both important for targeting FisB to the fission site.

### *C. perfringens* FisB can substitute for *B. subtilis* FisB

So far, our results suggest FisB-FisB and FisB-acidic lipid interactions are the main drivers for targeting FisB to the membrane fission site. If no other partners are involved, FisB should be largely an independent fission module, i.e. FisB homologs from different sporulating bacteria should be able to substitute for one another at least partially, even if sequence homology is low outside the consensus region. To test this idea, we expressed *Clostridium perfringens* FisB (FisB*^Cperf^*) in *B. subtilis* cells lacking FisB (BAL005). The sequence identity is only 23% between FisB sequences from these two species. In the heat-kill assay, FisB*^Cperf^* fully rescued *B. subtilis ΔfisB* defects (Figure 7A). *C. perfringens* FisB fused to mEGFP (mEGFP-FisB*^Cperf^*) had similar dynamics as FisB*^Bsubti^*, forming DMCs at early times that gave way to an ISEP where membrane fission occurs (Figure 7B). Population kinetics of membrane fission were slower with FisB*^Cperf^* (Figure 7C), but nearly every cell that underwent fission had an ISEP as for the wild type protein (220/239, or 92%). The intensity distribution of mEGFP-FisB*^Cperf^* ISEP was shifted to smaller values compared to mEGFP-FisB*^Bsubti^* ISEP (Figure 7D). Since the average ISEP intensity for FisB*^Bsubti^* corresponds to ∼40 copies (Figure 2), we deduce ∼9 copies of FisB*^Cperf^* accumulate at ISEP at the time of membrane fission. At t=3 h into sporulation, the percentage of cells with an ISEP was lower for cells expressing mEGFP-FisB*^Cperf^* (Figure 7E).

In all conditions tested so far, nearly all cells that had undergone membrane fission also had an intense FisB spot at the engulfment pole (Figs. 2,3,6, and 7). When we plotted the percentage of cells having an ISEP against the percentage of cells that have undergone fission at t=3 h, we found a nearly perfect correlation (Figure 7F). FisB*^Cperf^* fit this pattern well, despite having a low sequence identity to FisB*^Bsubti^*, suggesting a common localization and membrane fission mechanism, likely based on a few conserved biophysical properties.

### FisB does not have a preference for highly curved membrane regions, but can bridge membranes

A number of proteins localize to sub-cellular sites due to their preference for curved membrane regions^59-62^. During late stages of engulfment, the most highly curved region in the cell is the membrane neck connecting the engulfment membrane with the rest of the mother cell membrane and this is where FisB accumulates. We therefore asked whether curvature-sensing could be a mechanism driving FisB’s localization. To test this possibility, we undertook three independent series of experiments.

First, we used the principle that any protein which preferentially binds curved membranes at low membrane coverage can also induce membrane curvature when present at sufficiently high coverage^60,63^. Thus, we tested whether the soluble ECD of FisB could generate curved regions in highly malleable membranes of giant unilamellar vesicles (GUVs) at high coverage. We incubated 2 μM purified soluble FisB ECD labeled with iFluor555 with GUVs and monitored protein coverage and membrane deformations using spinning-disc confocal microscopy. Even when the GUV membranes were covered uniformly with iFluor555-FisB ECD we could not observe any GUV membrane deformations (Figure 8A). As a positive control, we used purified Endophilin A1 (EndoA1, labeled with Atto395), an N-BAR domain containing endocytic protein^64-66^. We incubated 2 μM EndoA1 with GUVs composed of 45% DOPS, 24.5% DOPC, 30% DOPE and 0.5% DiD, which resulted in extensive tubulation of GUV membranes (Figure 8A), as reporte previously^67^. Importantly, the difference in the membrane sculpting ability of the two proteins is not due a weaker affinity of FisB ECD for membranes (*K_d_* ≈ 1 µM for membranes with 45 mole % CL, Figure 4F) compared to endophilin (*K_d_* = 1.15 µM for membranes containing 45% DOPS, 30% DOPE, 24.5% DOPC, 0.5% TR-DHPE^65^).

Second, we slowly deflated GUVs to facilitate any potential membrane curvature generation by FisB ECD (which works against membrane tension) and/or to provide curved regions to test if FisB ECD accumulated there. Deflated GUVs displayed curved regions because their larger surface-to-volume ratios no longer allowed spherical shapes. Even under these favorable conditions, FisB ECD was not able to generate highly curved regions on these deflated GUVs (Figure 8B). In addition, if FisB ECD had a preference for negatively (positively) curved regions, it should accumulate at such regions while being depleted from positively (negatively) curved areas. Quantification of FisB ECD coverage at negatively or positively curved membrane regions showed no curvature preference (Figure 8B).

Third, we tested if FisB’s localization in live *B. subtilis* cells depended on membrane curvature. To avoid potentially confounding effects of other cues that may be present during sporulation, we expressed GFP-FisB under an inducible promoter during vegetative growth. In addition, we blocked cell division by inducing expression of MciZ^68^. MciZ normally blocks binary cell division during sporulation, but when expressed during vegetative growth, cells grow into long flexible filaments that are bent to varying degrees, providing regions with different membrane curvatures. We imaged GFP-FisB spots along curved edges of these filaments and plotted the linear density of GFP-FisB spots (spots/μm) as a function of filament curvature (Figure 8C). There was no clear correlation between GFP-FisB spot density and filament curvature. Although this method generates a limited amount of curvature, a similar approach was previously used to show that DivIVA preferentially localizes to negatively curved regions^69^.

In the GUV experiments, we noticed that FisB ECD caused GUVs to adhere to one another when they came into contact, accumulating at the adhesion patch between the membranes and at the rims (Figure 8D). Absorbance measurements using small unilamellar vesicles (SUVs) confirmed that FisB ECD can bridge membranes and aggregate liposomes (Figure 8E).

Overall, these experiments suggest that FisB does not have any intrinsic membrane curvature sensing/sculpting ability, but it can bridge membranes.

### Modeling suggests self-oligomerization and membrane bridging are sufficient to localize FisB to the membrane neck

To test the hypothesis that the homo-oligomerization, lipid-binding and the unique membrane topology could be sufficient to recruit FisB to the membrane neck, we considered a minimal model based on free energy minimization. As depicted in Figure 9, we consider the free energy *F* of an axisymmetric membrane neck of radius *R* and length *L* connecting two membrane sheets, corresponding to the local geometry where the engulfment membrane meets the rest of the mother cell membrane. We assume the surface density *ɸ* of FisB proteins in the neck is uniform and reaches equilibrium with a surface density *ɸ*_0_ of FisB in the surrounding membranes, and ask whether the neck geometry alone is enough to account for the observed FisB accumulation. The energy functional consists of a term accounting for membrane bending and tension, *F*_m_, and another term accounting for FisB protein-protein interactions, *F*_p_. We employ the classical Helfrich-Canham theory^70-75^ for the energy of the membrane

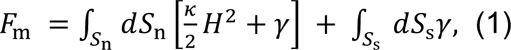

where *S*_1_ and *S*_s_ are the surfaces of the membrane neck and sheets, *H* is twice the mean curvature, *κ* is the bending modulus, and *γ* is the surface tension. The two membrane sheets are assumed to be planar, thus their only contribution to the energy comes from membrane tension.

For FisB, we include translational entropy, the energy of homo-oligomerization in trans between opposing membranes, and an energy that limits crowding. As shown above, FisB proteins do not exhibit curvature sensing, so we do not include a term coupling FisB density to membrane curvature in Eq. (1). This results in the following expression for the protein free energy

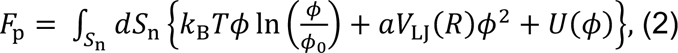

where the first term accounts for translational entropy, the second term is an energy per unit area accounting for trans interactions of FisB, and which for simplicity is assumed to be proportional to the standard Lennard–Jones (LJ) potential accounting for a longer-range attraction and shorter-range repulsion,

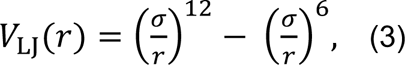

and finally *U*(*ɸ*) is an energy penalty for crowding that increases rapidly above a certain FisB concentration. To obtain *U*(*ɸ*) we assume a purely repulsive, truncated and shifted LJ potential between cis-neighboring FisB molecules, which we take to occupy a triangular lattice. Therefore, *U*(*ɸ*) = *ε*[*V*_LJ_(*r*(ɸ)) − *V_LJ_*(*r*_max_)] when *r* ≤ *r*_max_ and 0 when *r* > *r*_max_, where we have chosen *r*_max_ = 2^1/6^σ_cis_, namely the minimum of the LJ potential with length scale σ_cis_. The result is

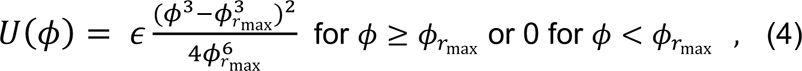

Where 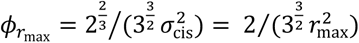 is the FisB concentration corresponding to a nearest neighbor distance *r*_max_.

We minimize *F* = *F*_0_ + *F*_4_ with respect to *ɸ* to obtain an equation for the equilibrium density of FisB proteins in the neck

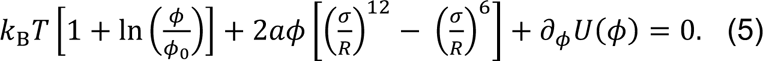

Then, minimizing *F* with respect to *R* yields an equation that determines the equilibrium radius of the neck

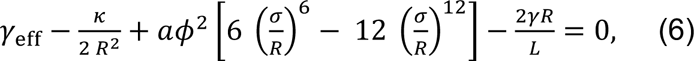

where 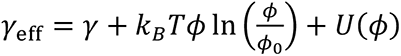.

Figure 9B shows *R* and *ɸ* as functions of the FisB trans homo-oligomerization strength *a*, for different values of surface tension *γ* and neck length *L*. For realistic parameters (the dot), we find that FisB trans interactions are strong enough to stabilize the neck at *R* ∼ 3 nm, with a close-packed concentration of FisB in the neck *ɸ* ≈ *ɸ*_max_. For these same parameters, there is a critical lower limit of *a* below which the FisB interactions are too weak to stabilize the neck, so the neck opens, i.e. *R* → ∞ in our simple model. Additionally, Figure 9B shows that the shorter the length of the neck *L*, the stronger the trans interactions needed to stabilize the neck at a finite radius. This makes intuitive sense: the longer the neck, the more FisB can be present to hold the neck together in opposition to membrane tension. (Note that expanding the radius of the neck actually decreases the total membrane area, which is the sum of the membrane in the neck and in the parallel sheets, so that surface tension tends to make the neck expand – see S2 Appendix).

While the above results suggest that an accumulation of FisB at the neck can be energetically stable, one question is how long it might take to reach that state? We expect nucleation of a critical cluster of FisB to be rate limiting, since the time required for diffusion and capture to reach ∼40 FisB in the neck is quite short ∼3.9 s (see S2 Appendix). To obtain a simple estimate of the nucleation time for both low-expression and native-expression strains, we assume that FisB proteins diffuse independently on the entire membrane and that nucleation of a stable cluster in the neck occurs when *n* proteins happen to be in the neck at the same time. To this end, we need to estimate the fraction of time there are *n* or more FisB in the neck, as well as the correlation time, i.e. the time between uncorrelated samples. Since we assume FisB proteins are independent, the number of proteins in the neck will be Poisson distributed, so we only need to know the average in the neck to obtain the full distribution. The average number of FisB in the neck is its area, 2π*RL*, times the background concentration, *ɸ*_0_. Furthermore, the correlation time is simply the time for a FisB to diffuse the length of the neck *L*^2^/*D*. Using *ɸ*_0_ ≃ 20 FisB *μ*m^-2^ (see “About 40 FisB molecules accumulate at the engulfment pole to mediate membrane fission” above) for the low-expression strain yields <FisB>≃ 0.03 in the neck. Assuming that the one-hour delay in membrane fission during sporulation of low-expression strain is due to the time for nucleation, we can infer that the number of FisB proteins required for nucleation is *n* ≈ 3 (see S2 Appendix). If the native-expression strain also needs *n* ≈ 3 to nucleate, we can estimate its corresponding nucleation time using *ɸ*_0_ ≃ 100 FisB *μ*m^-2^, which yields <FisB>≃ 0.15 and a nucleation time of ∼30 s. We conclude that for native expression levels of FisB, nucleation of a stable cluster of FisB at the neck is not likely to be rate limiting for the process of membrane fission.

## DISCUSSION

Previously, we showed that FisB is required for the membrane fission event that marks the completion of engulfment of the forespore by the mother cell^23^. Here, we found that a cluster of FisB molecules is nearly always present at the membrane fission site as evidenced by an intense fluorescent spot at the engulfment pole (ISEP) using fluorescently tagged FisB. The number of FisB molecules accumulated at the ISEP correlates well with the fraction of cells having undergone membrane fission at a given time point after induction of sporulation (Figs. 1,7). In addition, the number of wild-type FisB molecules per ISEP correlates with the total number of FisB molecules per cell (S1 Appendix Figure 4). Thus, the kinetics of membrane fission is determined by the accumulation of FisB molecules at the fission site. Lowering FisB expression could slow membrane fission by slowing the accumulation of FisB at the pole, or by reducing the number of FisB molecules driving fission after they are localized at the fission site. Our modeling results are consistent with slower ISEP nucleation in the low-expression strain, however, currently, we cannot experimentally distinguish between the two possibilities, and both may be operating simultaneously.

How is FisB recruited to the fission site? Our results suggest FisB does not rely on existing landmarks, lipid microdomains, cell-wall remodeling machinery, pH or voltage gradients across the cell membrane, or membrane curvature cues for its dynamic localization. In addition, we could not detect proteins interacting with FisB other than itself using an anti-GFP resin pulling on YFP-FisB^23^. By contrast, we found self-oligomerization and binding to acidic lipids to be critical for FisB’s function, and purified FisB ECD can bridge artificial membranes. Together, these results suggest FisB-FisB and FisB-lipid interactions are key drivers for FisB clustering and accumulation at the membrane fission site.

Can FisB oligomerization and lipid binding be sufficient to accumulate an immobile cluster of FisB molecules at the engulfment pole? Modeling suggests this is indeed the case. First, the narrow neck enables FisB’s on opposing membranes to come close enough to interact in trans. We infer this to be the preferred orientation for FisB-FisB interaction, since otherwise large clusters would be expected to form elsewhere as well. Second, the unique geometry of the neck connecting the engulfment membrane to the rest of the mother cell membrane plays an important role, as this is the only region in the cell where a cluster of FisB molecules can be “trapped”, i.e., once a cluster is formed inside the neck, it cannot diffuse away without breaking apart. This idea is supported by the fact that we do not observe any FisB accumulation at the leading edge of the engulfment membrane until a thin neck has formed at the end of engulfment.

The first FisB oligomers that appear during sporulation are dim, mobile clusters (DMCs), each containing about a dozen FisB molecules. (One possibility is that the DMCs may correspond to local membrane folds stabilized by FisBs interacting in trans.) Diffusion of DMCs appears to be Brownian on the 10-20 s time scale (Figure 2), though a rigorous analysis would require taking into account the geometry of the system. A DMC can diffuse a typical distance of ∼1 μm in ∼ 5 min (*D*_*DMC*_ ≈ 3 × 10^-3^µm^2^/s, Figure 2E). By comparison, engulfment in individual cells takes ∼60 min on average^76^. Though the engulfment time is much longer than the DMC diffusion time, the neck region, with an inner diameter of several nanometers, only forms at the very end of the engulfment process. Thus, ∼40 FisB molecules could be recruited to the neck through diffusion-limited capture of a few DMCs. However, we could not image such capture events directly, and cannot rule out that FisB can also diffuse as monomers and could be recruited to the neck in that form. Indeed, a simple model of the rate of nucleation of a cluster of FisBs at the neck suggests that as few as three FisBs interacting in trans could be sufficient to form a stable cluster there, with a nucleation time significantly shorter than the engulfment time at native expression levels.

How many FisB molecules are needed for efficient membrane fission? In cells completely lacking FisB, ∼5% of the cells undergo membrane fission by t=3 h, compared to ∼80 % or ∼30% for cells expressing FisB at native or ∼8-fold reduced levels, respectively (Figure 1F). The former achieve fission with ∼40 copies, while the latter with only ∼6. Thus, FisB is not absolutely required for membrane fission, but it makes it much more efficient, i.e. FisB catalyzes membrane fission. The variable stoichiometry suggests that FisB does not oligomerize into a specific quaternary structure with a definite stoichiometry. This variability appears to be a common property among proteins catalyzing membrane fusion and fission, such as SNAREs^77-79^ or dynamin^14^. The smallest clusters associated with membrane fission had ∼6 FisB copies on average. This number is likely sufficient to form at least one ring inside the membrane neck that eventually undergoes fission. Given that fission can occur in the absence of FisB, it is likely that the FisB cluster cooperates with other cellular processes to produce stress on this membrane neck.

We found FisB dynamics and membrane fission are not affected by removal of CL, PE, or both. CL and PE are widely implicated in membrane fission and fusion reactions due to their tendency to form non-bilayer structures^48,80-83^. The fact that CL or PE do not affect membrane fission during sporulation is remarkable, because such lipids usually affect the kinetics and/or the extent of fusion/fission reactions even if they are not absolutely required^81^. We tested the role of CL in a strain that lacked all three known CL synthases, with no detectable CL levels. A previous study reported that in *ΔclsABC B. subtilis* cells, CL levels increase from undetectable during vegetative growth to readily detectable during sporulation^31^, suggesting a yet unidentified sporulation-specific CL synthase may exist. Our results differ from those of Kawai et al. in that we were unable to detect any CL in *ΔclsABC B. subtilis* cells during vegetative growth or sporulation. We suggest the differences may be due to the different strain backgrounds used^84^, PY79^85^ here vs. Bs168^86^ in Kawai et al. or differences in detection sensitivities.

Overall, our results suggest FisB localizes to the membrane fission site using only lipid-binding, homo-oligomerization, and the unique geometry encountered at the end of engulfment. We propose that accumulation of a high enough density of FisB leads to membrane fission, possibly by generating increased stress in the FisB network-membrane composite, or in cooperation with another cellular process. A FisB homologue with low sequence identity partially rescued fission defects in *ΔfisB B. subtilis* cells, consistent with the idea that FisB acts as an independent module relying mainly on homo-oligomerization, lipid-binding, and sporulation geometry.

## MATERIALS AND METHODS

### Materials

*E. coli* cardiolipin (CL), *E. coli* L-α-phosphatidylglycerol (PG), egg L-α-phosphatidylcholine (eggPC), *E.coli* L-α-phosphatidylethanolamine (PE), 1,2-dioleoyl-sn-glycero-3-phosphoethanolamine-N-(7-nitro-2-1,3-benzoxadiazol-4-yl) (NBD-PE), 1,2-dioleoyl-sn-glycero-3-phosphoethanolamine (DOPE), 1,2-dioleoyl-sn-glycero-3-phosphocholine (DOPC), 1,2-dioleoyl-sn-glycero-3-phospho-L-serine (DOPS) were purchased from Avanti Polar Lipids. 1-(4-Trimethylammoniumphenyl)-6-Phenyl-1,3,5-Hexatriene *p*-Toluenesulfonate (TMA-DPH) and *N*-(3-Triethylammoniumpropyl)-4-(6-(4-(Diethylamino) Phenyl) Hexatrienyl) Pyridinium Dibromide (FM4-64), and 1,1’-Dioctadecyl-3,3,3’,3’-Tetramethylindodicarbocyanine (DiD) were from Thermo Fisher Scientific. Molybdenum Blue spray reagent was from Sigma-Aldrich. Carbonyl cyanide m-chlorophenyl hydrazone (CCCP) was purchased from Abcam and valinomycin was purchased from VWR. 3-(N-maleimidylpropionyl)biocytin (MBP) was obtained from Invitrogen and the HRP-conjugated antibody from eBioscience. Zaragozic acid was purchased from Sigma-Aldrich. 4-acetamido-4′-maleimidylstilbene-2,2′-disulfonic acid (AMS) and zaragozic acid were from obtained from Cayman Chemical Company.

### General *B. subtilis* methods

*B*. *subtilis* strains were derived from the prototrophic strain PY79^85^. Sporulation was induced in liquid medium at 37°C by nutrient exhaustion in supplemented DS medium (DSM)^87^ or by resuspension according to the method of Sterlini & Mandelstam^88^. Sporulation efficiency was determined in 24–30 h cultures as the total number of heat-resistant (80°C for 20 min) colony forming units (CFUs) compared to wild-type heat-resistant CFUs. Lipid synthesis mutants were from the *Bacillus* knock-out (BKE) collection^89^ and all were back-crossed twice into *B*. *subtilis* PY79 before assaying and prior to antibiotic cassette removal. Antibiotic cassette removal was performed using the temperature-sensitive plasmid pDR244 that constitutively expresses Cre recombinase^89^. Cassette removal was further confirmed by PCR with primers flanking the deletion. *B*. *subtilis* strains were constructed using plasmidic or genomic DNA and a 1-step competence method. Site directed mutagenesis was performed using Agilent’s Quick-change Lightning kit following manufacturer’s instructions and mutations were confirmed by sequencing. The strains and plasmids used in this study are listed in S1 Appendix Tables 2 and 3, respectively.

### Live-cell fluorescence microscopy of *B. subtilis*

Cells were mounted on a 2% agarose pad containing resuspension medium using a gene frame (Bio-Rad). Cells were concentrated by centrifugation (3300g for 30 s) prior to mounting and visualization. This step had no impact on the localization of the fusion proteins. Fluorescence microscopy was performed using a Leica DMi8 wide-field inverted microscope equipped with an HC PL APO 100×DIC objective (NA=1.40) and an iXon Ultra 888 EMCCD Camera from Andor Technology. Membranes were stained with TMA-DPH at a final concentration of 100 μM. Excitation light intensity was set to 50% and exposure times were 300 ms for TMA-DPH (*λ*_ex_=395/25 nm; *λ*_em_=460/50 nm); 500 ms for m(E)GFP (*λ*_ex_=470/40; *λ*_em_=500-550) and 1 s for mYFP (*λ*_ex_=510/25; *λ*_em_>530) respectively. Images were acquired with Leica Application Suite X (LAS X) and analysis and processing were performed using the ImageJ software^90^.

### Determination of FisB’s topology

We used the substituted cysteine accessibility method (SCAM^91^) to determine the topology of FisB. We first generated stains expressing FisB versions with a single cysteine substitution at position G6, L137, or A245, in a *ΔfisB* background. FisB does not have any endogenous cysteines. These point mutations decreased the sporulation efficiency slightly (S1 Appendix Table 2), we assume without affecting the topology. We selectively biotinylated extra- or intracellular cysteines of *B. subtilis* protoplasts, produced by addition of 0.5 mg/ml lysozyme and incubating cells at 37°C for 1h with gentle rocking. Protoplasts were then incubated with the membrane-impermeant reagent 3-(N-maleimidylpropionyl)biocytin (MBP). To selectively label extracellular cysteines, protoplasts of sporulating cells at 2.5 h into sporulation were incubated with 100 μM MPB. The reaction was quenched with 50 mM DTT before cells were lysed with hypotonic shock. To label intracellular cysteines selectively, extracellular cysteines of protoplasts were first blocked AMS before cells were lysed and incubated with 100 μM MPB. The reaction was quenched by addition of 100 μM MPB. FisB was pulled down from the cell lysates as described in^91^ using an anti-Myc antibody (mAb #2276) and biotinylated proteins were detected by Western Blot using a HRP-conjugated-Avidin antibody. Further details are provided in the S1 Appendix.

### Expression, purification, and labeling of recombinant FisB protein

Recombinant soluble FisB ECD was purified as described in^23^ but with slight modifications. Briefly, His_6_-FisB ECD was expressed in *E. coli* BL21 (DE3) from New England Biolabs and purified using HisPur™ Ni-NTA Resin from Thermo Fisher Scientific. Protein expression was induced with 1 mM IPTG at OD_600_ = 0.6 overnight at 16°C. Cells were harvested by centrifugation and the pellet was resuspended in Lysis Buffer (20 mM HEPES, 500 mM NaCl, 0.5 mM TCEP, 20 mM Imidazole, 2% glycerol, 20 mM MgCl2) and flash-frozen in liquid nitrogen. Pellets were thawed on ice and cells were lysed by 5 passes through a high-pressure homogenizer (Avestin EmulsiFlex-C3). The lysate was spun down at 100,000×g and the soluble fraction was incubated with HisPur™ Ni-NTA Resin for 2.5 h at 4°C while rotating. The bound protein was washed with Lysis Buffer, Lysis Buffer containing 50 mM and finally 100 mM Imidazole. The protein was eluted in Elution Buffer (20 mM HEPES, 500 mM NaCl, 0.5 mM TCEP, 200 mM Imidazole, 2% glycerol, 20 mM MgCl2). The protein was concentrated using a Vivaspin centrifugal concentrator with a 10 kDa molecular weight cutoff and the concentration determined by Bradford protein assay. The protein was stored at -80°C.

In experiments with labeled FisB ECD, we used a cysteine mutation, G123C (FisB ECD does not have any endogenous cysteines). After expression and purification as above, iFluor555-maleimide (AAT Bioquest) was reacted with FisB ECD^G123C^ following the manufacturer’s instructions. G123 is in a loop that if removed does not interfere with FisB’s function (S1 Appendix Figure 10).

### Analytical size-exclusion chromatography (SEC) and negative-stain electron microscopy (EM)

For SEC analysis His_6_-FisB ECD was loaded onto a Superose 6 Increase 10/300 GL column (GE) previously equilibrated with 20 mM HEPES, pH 7.5, 500 mM NaCl, 0.5 mM TCEP, 2% glycerol, 20 mM MgCl2, running at a flow rate of 0.5 ml/min at 4°C. The column was calibrated with Bio-Rad’s Gel Filtration Standards. For negative stain EM analysis, 4 μL of the indicated elution fractions were applied to 200-mesh copper grids coated with ∼10 nm amorphous carbon film, negatively stained with 2% (wt/vol) uranyl acetate, and air-dried. Images were collected on a FEI Tecnai T12 microscope, with a LaB6 filament operating at 120 kV, and equipped with a Gatan CCD camera.

### Inhibition of cell wall synthesis and analyses of FisB motions

Overnight cultures of GFP-Mbl (BDR2061) or IPTG-induced mGFP-FisB (BMB014) were diluted in CH medium to OD_600_ = 0.05. Expression of GFP-FisB was induced with 1 mM IPTG for 2h at 37°C. Expression of GFP-Mbl was induced with 10 mM xylose for 30 min when BDR2061 reached OD_600_ = 0.5. For imaging untreated cells, 1 ml of cells was washed twice with 1 ml PBS and finally resuspended in 10 μl PBS. 2 μl of cell suspension was spread on a 2% PBS agar pad for imaging. To inhibit cell-wall synthesis 50 μg/ml fosfomycin was added to the cultures 45 min before imaging. 1 ml of cells was washed twice with PBS containing 50 μg/ml fosfomycin and mounted on a PBS agar pad also containing fosfomycin. Cells were imaged using a Olympus IX81 microscope with a home-built polarized TIRF setup^92,93^. Exposure times were 50 ms for BDR2061 and 100 ms for BMB014. Movies were acquired at 1 frame/s. Movies collected for BMB014 were corrected for bleaching using the Bleaching Correction function (exponential method) in ImageJ. Kymographs were created with imageJ along the indicated axes. GFP fusion proteins were tracked using the ImageJ plugin TrackMate^94^. A Laplacian of Gaussian (LoG) filter was used to detect particles with an estimated blob diameter 400 μm. Particles were tracked using the Simple LAP tracker with a 0.25 μm maximum linking distance and no frame gaps. MATLAB (Mathworks, Natick, MA) was used for further processing of the tracks. Mean squared displacement (MSD) was calculated using the MATLAB class @msdanalyzer^95^.

The asymmetry of individual tracks (S1 Appendix Fig. 5F) was calculated as described in^96^ using:

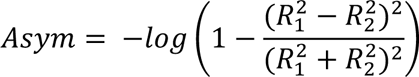

where *R*_1_ and *R*_2_ are the principal components of the radius of gyration, equal to the square roots of the eigenvalues of the radius of gyration tensor ***R*_*g*_**:

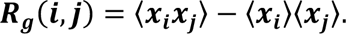

### Tracking fluorescently labeled FisB spots and estimation of diffusion coefficients

For estimating the mobility of DMC and ISEP, time-lapse movies were recorded with a frame rate of 1 s using wide-field microscopy (50% LED intensity, 300 ms exposure time, gain 300). Spot positions were tracked using SpeckleTrackerJ^97^, a plugin for the image analysis software ImageJ^90^. Mean-squared displacements (MSDs) were calculated using the MATLAB class @msdanalyzer^95^.

### Dissipation of membrane potential

Cells were concentrated by centrifugation (3300xg for 30 s) and 100 μM CCCP or 30 μM valinomycin was added just prior to mounting cells onto a 2% PBS agar pad also containing 100 μM CCCP or 30 μM valinomycin.

### Lipid extraction and thin-layer chromatography (TLC)

Lipids were extracted from *B. subtilis* cells at 3 h into sporulation according to the method of Lacombe and Lubochinsky^98^. Lipid extracts were analyzed by TLC on silica gel plates in mixtures of chloroform:hexane:methanol: acetic acid (50:30:10:5). Phospholipids were detected with Molybdenum Blue Reagent (Sigma-Aldrich).

### Liposome preparation

Small unilamellar vesicles (SUVs) were prepared by mixing 1 μmol of total lipids at desired ratios. A thin lipid film was created using a rotary evaporator (Buchi). Any remaining organic solvent was removed by placing the lipid film under high vacuum for 2h. The lipid film was hydrated with 1 ml of RB-EDTA buffer [25 mM HEPES at pH 7.4, 140 mM KCl, 1 mM EDTA, 0.2 mM tris(2-carboxyethyl) phosphine] by shaking using an Eppendorf Thermomix for >30 min. The lipid suspension was then frozen and thawed 7 times using liquid nitrogen and a 37°C water bath and subsequently extruded 21 times through a 100 nm pore size polycarbonate filter using a mini-extruder (Avanti). All SUVs contained 1% NBD-PE to determine the final lipid concentration.

Giant unilamellar vesicles (GUVs) were prepared by electroformation^99^. Briefly, lipids dissolved in chloroform were mixed in a glass tube at desired ratios and spotted on two indium tin oxide (ITO) coated glass slides. Organic solvent was removed by placing the lipid films in a vacuum desiccator for at least 2 h. A short strip of copper conductive tape was attached to each ITO slide which were then separated by a 3 mm thick Polytetrafluoroethylene (PTFE) spacer and held together with binder clips. The chamber was filled with 500 μl Swelling Buffer (1 mM HEPES, 0.25 M sucrose, 1 mM DTT) and sealed with Critoseal (VWR International, Radnor, PA). GUVs were formed by applying a 1.8 V sinusoidal voltage at 10 Hz for at least 2 h at room temperature.

For experiments involving FisB ECD the GUVs were composed of (all in mole percentages): 25 *E. coli* PE, 5 *E. coli* CL, 50 *E. coli* PG, 19 eggPC and 1 DiD or 1 NBD-PE. For experiments in which EndoA1 was used, GUV composition was (all in mole %): 45 DOPS, 24.5 DOPC, 30 DOPE and 0.5 DiD.

### Liposome-protein co-floatation

For initial experiments, 40 nmol total lipid was incubated with 200 pmol FisB ECD for 1h at room temperature in a total volume of 100 μl. 200 μl of 60% Optiprep (iodixanol, Sigma-Aldrich) was added to the sample creating a 40% Optiprep solution. The sample was then layered at the bottom of a 5 mm x 41 mm Beckman ultracentrifuge tube (#344090) and overlaid with 200 μl of 20% Optiprep and finally 150 μl of buffer (Figure 4 C). Liposome-bound proteins co-float to a light density, while unbound proteins pellet upon ultracentrifugation for 1.5 h at 48 krpm. Fractions were collected as shown in Figure 4C and the amount of recovered protein was determined by SDS-PAGE (Nu-PAPGE 12% Bis-tris gel, Thermo Fisher Scientific) stained with SYPRO^TM^ Orange (Invitrogen).

### Liposome aggregation using absorbance

SUVs were prepared by extrusion as described above but using a 50 nm polycarbonate filter. SUVs were composed of 50 mole % *E. coli* PG, 25 mole % *E. coli* PE, 20 mole % eggPC, 5 mole % *E. coli* CL. The absorbance at 350 nm of 50 μM total lipid was measured for 5 min, before addition of 1 μM FisB ECD. Absorbance increases with increasing liposome aggregation due to increased scattering^100^.

### Filamentous *B. subtilis* cells to test for curvature-sensitive localization of FisB

An overnight culture of BMB014 was diluted into fresh CH medium^101^ to OD_600_=0.05. 1 mM IPTG and 20 mM xylose were added to induce the expression of GFP-FisB and MciZ, respectively. The latter inhibits cytokinesis^68^. The culture was grown at 37°C for 30 min before 3-5 μl of cells were transferred onto a 3% agar pad also containing 1 mM IPTG and 20 mM xylose. Cells were grown on the agar pad for 2h at 37°C prior to imaging. GFP-FisB foci were detected using the ImageJ plugin TrackMate as described above. Radii of the inner and outer edges were determined by manually fitting a circle to the cells using ImageJ.

### Determination of binding constants

For determination of binding constants, the floatation protocol was slightly modified. Varying amounts of lipids were incubated with 100 nM iFluor555-FisB ECD for 1 h at room temperature in a total volume of 100 μl. Density gradients were created as before using Optiprep (iodixanol), however only 2 fractions were collected (Figure 5H). The protein concentration in fraction A was too small to be quantified by SDS-PAGE. Therefore, the sample was concentrated by trichloroacetic acid (TCA) precipitation. Briefly, 50 μl of TCA was added to fraction A and incubated for 30 min at 4°C. The sample was spun at 14 krpm in an Eppendorf microfuge for 5 min. The pellet was washed twice with ice-cold acetone and subsequently dried for 10 min in a 95°C heating block. 10 μl of 2X SDS sample buffer was added to the dried pellet and the sample was boiled for 10 min at 95°C and loaded on a 12% bis-tris gel. The amount of recovered protein was determined by fluorescence intensity of the labeled FisB ECD band on the gel using a Typhoon FLA 9500 (GE Healthcare). The dissociation constant *K_d_* was determined following ref. 102. Titration curves were fitted to:

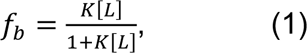

where *f_b_* is the fraction of bound protein and *K* the apparent association constant (*K* = 1/*K_d_*). Eq. (1) assumes that the total lipid concentration [*L*] is much larger than the concentration of bound protein, a condition satisfied in our experiments for [*L*] > 10^-7^ M.

### Image analysis

To estimate the fraction of cells that have undergone membrane fission at a particular time after sporulation was initiated by nutrient downshift, cells were labeled with TMA-DPH (see “Live-cell fluorescence microscopy of *B. subtilis*” above). The dye labels the forespore contours intensely before membrane fission, as it has access to three membranes in close proximity that cannot be resolved (forespore, engulfment, and mother cell membranes). After membrane fission, the dye dimly labels forespore contours (see S1 Appendix Fig. 1 for examples and quantification). Due to the clear separation between the two labeling patterns (S1 Appendix Fig. 1) cells can be scored visually, with 6-7% of cells having intermediate labeling that prevents categorization. Thus, we underestimate the percentage of cells that have undergone membrane fission by at most ∼7%.

For the analyses shown in S1 Appendix Figs. 4A,B,C,E, and 9, we calculated the total intensity (sum ox pixel values) inside the cell contour (indicated in yellow in Fig. S4A) using MicrobeJ^103^. Mean integrated auto-fluorescence (∼1300 a.u) was calculated by analyzing in the same way an equivalent number of individual wild-type cells, imaged under identical conditions.

For the analyses shown in Figure 2 and S1 Appendix Fig. 4D, FisB foci were semi-automatically selected using SpeckleTrackerJ^97^. For each spot the sum of pixel values in a 6 × 6 pixel (0.5 µm × 0.5 µm) box around the center of the spot were calculated. For each corresponding cell the same operation was performed at a membrane area where no clusters were present and subtracted from the FisB cluster intensity.

### Preparation of DNA Origami-based mEGFP standards

These standards were prepared and characterized as described in^44^. Briefly, DNA “rods” consisted of six-helix-bundle DNA origami nanotubes. Rods carried varying numbers of single stranded “handle” sequences for DNA-conjugated fluorophore hybridization. A long scaffold DNA (p7308^104^) was folded into the desired shape by self-assembly with a six-fold molar excess of designed “staple strands” by heating and cooling cycles over an 18-hour period in a thermocycler (Bio-Rad). Excess staples were removed by PEG precipitation^105^, and DNA-conjugated fluorophores were hybridized to the DNA origami nanotubes by coincubation for 2 hours at 37°C. Finally, excess fluorophore-DNA conjugates were removed by a second PEG precipitation^105^.

To estimate fluorophore labeling efficiency, standards designed to host 5 copies of Alexa Fluor 488 were similarly prepared. These standards were imaged on a TIRF microscope (Eclipse Ti, Nikon) until fully bleached. The photobleaching steps of the fluorescence traces were fit to a binomial function to estimate the labeling efficiency to be ∼80% (95% CI = 76%-84%).

### Quantitative Western Blot

mYFP was cloned into pVS001 (His6-Sumo-mYFP) and purified using affinity chromatography. For immunoblotting, cells in 100 ml sporulation medium were pelleted and the supernatant removed. The pellets were suspended in ice-cold lysis buffer (pH=7.5; 50 mM HEPES, 100 mM KCl, 3 mM MgCl_2_, 1 mM EGTA, 0.1% Triton X-100, 1 mM DTT, 1 mM PMSF, with one complete protease inhibitor tablet (Roche) to a final volume of 300 µl, and then we added 0.3 g acid-washed glass beads (425-600 µm, Sigma). After adding 150 µl boiling sample buffer (250 mM Tris-HCl, pH 6.8, 50% glycerol, 3.58 µM β-mercaptoethanol, 15% SDS, and 0.025% Bromophenol Blue), samples were incubated at 100°C for 5 min. Samples were centrifuged at 14,000 rpm in a desktop centrifuge at room temperature for 10 min and stored at -80°C. The blots were probed with peroxidase-conjugated anti-GFP antibody (ab13970). Images were scanned and quantified using ImageJ.

## Supporting information

S2 Appendix: Theoretical modeling

S1 Appendix: Supplementary Experiments

## ACKNOWLEDGEMENTS

We thank members of the Karatekin and Rudner laboratories for stimulating discussions. This work was supported by National Institute of General Medical Sciences and National Institute of Neurological Disorders and Stroke of the National Institutes of Health (NIH) under award numbers R01GM114513 and R01NS113236 (to EK), DP2GM114830 and R01GM132114 (to CL). The content is solely the responsibility of the authors and does not necessarily represent the official views of the National Institutes of Health. We thank Vladimir Polejaev and Jeorg Nikolaus (directors of the Yale West Campus Imaging Core), and Josh Lees (Yale Center for Cellular and Molecular Imaging Electron Microscopy Facility) for their help with imaging, Karin Reinisch in whose laboratory work by FH was carried out, Daniel R. Zeigler (Bacillus Genetic Stock Center) for helpful advice, and Alexander J. Meeske for some of the strains used in this study. NDW was supported by a NIH training grant (T32-EB09941). We gratefully acknowledge a Yale University Predoctoral Fellowship to MB. This work was supported in part by the National Science Foundation, through the Center for the Physics of Biological Function (PHY-1734030). AM-C acknowledges support from the Spanish MECD through the program Ayudas para la Formación de Profesorado Universitario (FPU), Grant No. FPU16/0256. We are grateful to Aurelien Roux (U. Genève) for the generous gift of Atto395-EndoA1.

## AUTHOR CONTRIBUTIONS

AL, MB, VS, and EK conceived the study. AL and MB performed experiments whose results are shown in the main figures. AA (Fig. S10), FH (Fig. 5, 7, S8), VS (Fig. S3) performed additional experiments. NDW and CL developed the DNA-origami fluorescence calibration method and contributed to the data in Fig. 2. CR, TD, and DR provided resources, training, and technical and conceptual input. They introduced EK, AL, MB and VS to *B. subtilis* and sporulation. EK and DR provided supervision and acquired funding. AMC and NW developed the model and wrote the corresponding sections. AL, MB, and EK and wrote the manuscript, with input from other co-authors. We thank Aurelien Roux for the generous gift of Atto395-EndoA1.

## REFERENCES

1. Haucke V, Kozlov MM. Membrane remodeling in clathrin-mediated endocytosis. J Cell Sci 131, (2018).

2. Campelo F, Malhotra V. Membrane fission: the biogenesis of transport carriers. Annu Rev Biochem 81, 407–427 (2012).

3. Ahmed I, Akram Z, Iqbal HMN, Munn AL. The regulation of Endosomal Sorting Complex Required for Transport and accessory proteins in multivesicular body sorting and enveloped viral budding - An overview. Int J Biol Macromol 127, 1–11 (2019).

4. Jaumouille V, Waterman CM. Physical Constraints and Forces Involved in Phagocytosis. Front Immunol 11, 1097 (2020).

5. Carlton JG, Jones H, Eggert US. Membrane and organelle dynamics during cell division. Nat Rev Mol Cell Biol 21, 151–166 (2020).

6. Errington J. Regulation of endospore formation in Bacillus subtilis. Nat Rev Microbiol 1, 117–126 (2003).

7. Higgins D, Dworkin J. Recent progress in Bacillus subtilis sporulation. FEMS Microbiol Rev 36, 131–148 (2012).

8. Tan IS, Ramamurthi KS. Spore formation in Bacillus subtilis. Environ Microbiol Rep 6, 212–225 (2014).

9. Rand RP, Parsegian VA. Mimicry and mechanism in phospholipid models of membrane fusion. Annu Rev Physiol 48, 201–212 (1986).

10. Wong JY, Park CK, Seitz M, Israelachvili J. Polymer-cushioned bilayers. II. An investigation of interaction forces and fusion using the surface forces apparatus. Biophys J 77, 1458–1468 (1999).

11. Kozlovsky Y, Kozlov MM. Membrane fission: model for intermediate structures. Biophys J 85, 85–96 (2003).

12. Bashkirov PV, Akimov SA, Evseev AI, Schmid SL, Zimmerberg J, Frolov VA. GTPase cycle of dynamin is coupled to membrane squeeze and release, leading to spontaneous fission. Cell 135, 1276–1286 (2008).

13. Kozlov MM, McMahon HT, Chernomordik LV. Protein-driven membrane stresses in fusion and fission. Trends Biochem Sci 35, 699–706 (2010).

14. Ferguson SM, De Camilli P. Dynamin, a membrane-remodelling GTPase. Nat Rev Mol Cell Biol 13, 75–88 (2012).

15. Schoneberg J, Lee IH, Iwasa JH, Hurley JH. Reverse-topology membrane scission by the ESCRT proteins. Nat Rev Mol Cell Biol 18, 5–17 (2017).

16. Simunovic M, et al. Friction Mediates Scission of Tubular Membranes Scaffolded by BAR Proteins. Cell 170, 172–184 e111 (2017).

17. Roux A, Cuvelier D, Nassoy P, Prost J, Bassereau P, Goud B. Role of curvature and phase transition in lipid sorting and fission of membrane tubules. EMBO J 24, 1537–1545 (2005).

18. Hatch AL, Gurel PS, Higgs HN. Novel roles for actin in mitochondrial fission. J Cell Sci 127, 4549–4560 (2014).

19. Yang C, Svitkina TM. Ultrastructure and dynamics of the actin-myosin II cytoskeleton during mitochondrial fission. Nat Cell Biol 21, 603–613 (2019).

20. Lacy MM, Ma R, Ravindra NG, Berro J. Molecular mechanisms of force production in clathrin-mediated endocytosis. FEBS Lett 592, 3586–3605 (2018).

21. Nickaeen M, Berro J, Pollard TD, Slepchenko BM. Actin assembly produces sufficient forces for endocytosis in yeast. Mol Biol Cell 30, 2014–2024 (2019).

22. Snead WT, et al. BAR scaffolds drive membrane fission by crowding disordered domains. J Cell Biol 218, 664–682 (2019).

23. Doan T, et al. FisB mediates membrane fission during sporulation in Bacillus subtilis. Genes Dev 27, 322–334 (2013).

24. Stragier P, Losick R. Molecular genetics of sporulation in Bacillus subtilis. Annu Rev Genet 30, 297–241 (1996).

25. Gest H, Mandelstam J. Longevity of microorganisms in natural environments. Microbiol Sci 4, 69–71 (1987).

26. Potts M. Desiccation tolerance of prokaryotes. Microbiol Rev 58, 755–805 (1994).

27. Ulrich N, Nagler K, Laue M, Cockell CS, Setlow P, Moeller R. Experimental studies addressing the longevity of Bacillus subtilis spores - The first data from a 500-year experiment. PLoS One 13, e0208425 (2018).

28. Brown JK, Hovmøller MS. Aerial dispersal of pathogens on the global and continental scales and its impact on plant disease. Science 297, 537–541 (2002).

29. Eichenberger P, et al. The sigmaE regulon and the identification of additional sporulation genes in Bacillus subtilis. J Mol Biol 327, 945–972 (2003).

30. Mileykovskaya E, Dowhan W. Visualization of phospholipid domains in Escherichia coli by using the cardiolipin-specific fluorescent dye 10-N-nonyl acridine orange. J Bacteriol 182, 1172–1175 (2000).

31. Kawai F, Shoda M, Harashima R, Sadaie Y, Hara H, Matsumoto K. Cardiolipin domains in Bacillus subtilis marburg membranes. J Bacteriol 186, 1475–1483 (2004).

32. Koppelman CM, Den Blaauwen T, Duursma MC, Heeren RM, Nanninga N. Escherichia coli minicell membranes are enriched in cardiolipin. J Bacteriol 183, 6144–6147 (2001).

33. Kawai F, Hara H, Takamatsu H, Watabe K, Matsumoto K. Cardiolipin enrichment in spore membranes and its involvement in germination of Bacillus subtilis Marburg. Genes Genet Syst 81, 69–76 (2006).

34. Lewis RN, McElhaney RN. The physicochemical properties of cardiolipin bilayers and cardiolipin-containing lipid membranes. Biochim Biophys Acta 1788, 2069–2079 (2009).

35. Haines TH. A new look at Cardiolipin. Biochim Biophys Acta 1788, 1997–2002 (2009).

36. Ortiz A, Killian JA, Verkleij AJ, Wilschut J. Membrane fusion and the lamellar-to-inverted-hexagonal phase transition in cardiolipin vesicle systems induced by divalent cations. Biophys J 77, 2003–2014 (1999).

37. Khalifat N, Puff N, Bonneau S, Fournier JB, Angelova MI. Membrane deformation under local pH gradient: mimicking mitochondrial cristae dynamics. Biophys J 95, 4924–4933 (2008).

38. Romantsov T, Helbig S, Culham DE, Gill C, Stalker L, Wood JM. Cardiolipin promotes polar localization of osmosensory transporter ProP in Escherichia coli. Mol Microbiol 64, 1455–1465 (2007).

39. Romantsov T, Stalker L, Culham DE, Wood JM. Cardiolipin controls the osmotic stress response and the subcellular location of transporter ProP in Escherichia coli. J Biol Chem 283, 12314–12323 (2008).

40. Lopez D, Koch G. Exploring functional membrane microdomains in bacteria: an overview. Curr Opin Microbiol 36, 76–84 (2017).

41. Sharp MD, Pogliano K. An in vivo membrane fusion assay implicates SpoIIIE in the final stages of engulfment during Bacillus subtilis sporulation. Proc Natl Acad Sci U S A 96, 14553–14558 (1999).

42. Meyer P, Gutierrez J, Pogliano K, Dworkin J. Cell wall synthesis is necessary for membrane dynamics during sporulation of Bacillus subtilis. Mol Microbiol 76, 956–970 (2010).

43. Doan T, et al. Novel secretion apparatus maintains spore integrity and developmental gene expression in Bacillus subtilis. PLoS Genet 5, e1000566 (2009).

44. Williams ND, et al. DNA-Origami-Based Fluorescence Brightness Standards for Convenient and Fast Protein Counting in Live Cells. Nano Lett 20, 8890–8896 (2020).

45. Guiziou S, et al. A part toolbox to tune genetic expression in Bacillus subtilis. Nucleic Acids Res 44, 7495–7508 (2016).

46. Donovan C, Bramkamp M. Characterization and subcellular localization of a bacterial flotillin homologue. Microbiology 155, 1786–1799 (2009).

47. Kunst F, et al. The complete genome sequence of the Gram-positive bacterium Bacillus subtilis. Nature 390, 249–256 (1997).

48. Chernomordik LV, Kozlov MM. Mechanics of membrane fusion. Nat Struct Mol Biol 15, 675–683 (2008).

49. Schmid SL, Frolov VA. Dynamin: functional design of a membrane fission catalyst. Annu Rev Cell Dev Biol 27, 79–105 (2011).

50. Nishibori A, Kusaka J, Hara H, Umeda M, Matsumoto K. Phosphatidylethanolamine Domains and Localization of Phospholipid Synthases in *Bacillus subtilis* Membranes. Journal of Bacteriology 187, 2163–2174 (2005).

51. Good MC, Zalatan JG, Lim WA. Scaffold proteins: hubs for controlling the flow of cellular information. Science 332, 680–686 (2011).

52. Langhorst MF, Reuter A, Stuermer CA. Scaffolding microdomains and beyond: the function of reggie/flotillin proteins. Cell Mol Life Sci 62, 2228–2240 (2005).

53. López D, Kolter R. Functional microdomains in bacterial membranes. Genes Dev 24, 1893–1902 (2010).

54. Sohlenkamp C, Geiger O. Bacterial membrane lipids: diversity in structures and pathways. FEMS Microbiology Reviews 40, 133–159 (2015).

55. Oliver PM, Crooks JA, Leidl M, Yoon EJ, Saghatelian A, Weibel DB. Localization of anionic phospholipids in Escherichia coli cells. J Bacteriol 196, 3386–3398 (2014).

56. Bogdanov M, Heacock PN, Dowhan W. Study of polytopic membrane protein topological organization as a function of membrane lipid composition. Methods Mol Biol 619, 79–101 (2010).

57. Ovchinnikov S, et al. Protein structure determination using metagenome sequence data. Science 355, 294–298 (2017).

58. den Kamp JA, Redai I, van Deenen LL. Phospholipid composition of Bacillus subtilis. J Bacteriol 99, 298–303 (1969).

59. Updegrove TB, Ramamurthi KS. Geometric protein localization cues in bacterial cells. Curr Opin Microbiol 36, 7–13 (2017).

60. Simunovic M, Voth GA, Callan-Jones A, Bassereau P. When Physics Takes Over: BAR Proteins and Membrane Curvature. Trends in Cell Biology 25, 780–792 (2015).

61. Antonny B. Mechanisms of membrane curvature sensing. Annu Rev Biochem 80, 101–123 (2011).

62. Baumgart T, Capraro BR, Zhu C, Das SL. Thermodynamics and mechanics of membrane curvature generation and sensing by proteins and lipids. Annu Rev Phys Chem 62, 483–506 (2011).

63. Sorre B, et al. Nature of curvature coupling of amphiphysin with membranes depends on its bound density. Proc Natl Acad Sci U S A 109, 173–178 (2012).

64. Kjaerulff O, Brodin L, Jung A. The structure and function of endophilin proteins. Cell Biochem Biophys 60, 137–154 (2011).

65. Chen Z, Zhu C, Kuo CJ, Robustelli J, Baumgart T. The N-Terminal Amphipathic Helix of Endophilin Does Not Contribute to Its Molecular Curvature Generation Capacity. J Am Chem Soc 138, 14616–14622 (2016).

66. Hohendahl A, et al. Structural inhibition of dynamin-mediated membrane fission by endophilin. Elife 6, (2017).

67. Shi Z, Baumgart T. Membrane tension and peripheral protein density mediate membrane shape transitions. Nat Commun 6, 5974 (2015).

68. Handler AA, Lim JE, Losick R. Peptide inhibitor of cytokinesis during sporulation in Bacillus subtilis. Mol Microbiol 68, 588–599 (2008).

69. Renner LD, Eswaramoorthy P, Ramamurthi KS, Weibel DB. Studying biomolecule localization by engineering bacterial cell wall curvature. PLoS One 8, e84143 (2013).

70. Canham PB. Minimum Energy of Bending as a Possible Explanation of Biconcave Shape of Human Red Blood Cell. J Theor Biol 26, 61-& (1970).

71. Helfrich W. Elastic Properties of Lipid Bilayers - Theory and Possible Experiments. Z Naturforsch C **C** 28, 693–703 (1973).

72. Zhongcan OY, Helfrich W. Instability and Deformation of a Spherical Vesicle by Pressure. Phys Rev Lett 59, 2486–2488 (1987).

73. Zhongcan OY, Helfrich W. Bending Energy of Vesicle Membranes - General Expressions for the 1st, 2nd, and 3rd Variation of the Shape Energy and Applications to Spheres and Cylinders. Phys Rev A 39, 5280–5288 (1989).

74. Seifert U, Langer SA. Viscous Modes of Fluid Bilayer-Membranes. Europhys Lett 23, 71–76 (1993).

75. Seifert U. Configurations of fluid membranes and vesicles. Adv Phys 46, 13–137 (1997).

76. Ojkic N, López-Garrido J, Pogliano K, Endres RG. Cell-wall remodeling drives engulfment during Bacillus subtilis sporulation. Elife 5, (2016).

77. Hernandez JM, Kreutzberger AJ, Kiessling V, Tamm LK, Jahn R. Variable cooperativity in SNARE-mediated membrane fusion. Proc Natl Acad Sci U S A 111, 12037–12042 (2014).

78. Mostafavi H, et al. Entropic forces drive self-organization and membrane fusion by SNARE proteins. Proc Natl Acad Sci U S A 114, 5455–5460 (2017).

79. Wu Z, et al. Dilation of fusion pores by crowding of SNARE proteins. Elife 6, (2017).

80. Stepanyants N, Macdonald PJ, Francy CA, Mears JA, Qi X, Ramachandran R. Cardiolipin’s propensity for phase transition and its reorganization by dynamin-related protein 1 form a basis for mitochondrial membrane fission. Mol Biol Cell 26, 3104–3116 (2015).

81. Chernomordik LV, Kozlov MM, Melikyan GB, Abidor IG, Markin VS, Chizmadzhev YA. The shape of lipid molecules and monolayer membrane fusion. Biochimica et Biophysica Acta (BBA) - Biomembranes 812, 643–655 (1985).

82. Cullis PR, de Kruijff B, Verkleij AJ, Hope MJ. Lipid polymorphism and membrane fusion. Biochem Soc Trans 14, 242–245 (1986).

83. Landajuela A, et al. Lipid Geometry and Bilayer Curvature Modulate LC3/GABARAP-Mediated Model Autophagosomal Elongation. Biophysical journal 110, 411–422 (2016).

84. Zeigler DR, et al. The origins of 168, W23, and other Bacillus subtilis legacy strains. J Bacteriol 190, 6983-6995 (2008).

85. Youngman PJ, Perkins JB, Losick R. Genetic transposition and insertional mutagenesis in Bacillus subtilis with Streptococcus faecalis transposon Tn917. Proc Natl Acad Sci U S A 80, 2305–2309 (1983).

86. Spizizen J. TRANSFORMATION OF BIOCHEMICALLY DEFICIENT STRAINS OF BACILLUS SUBTILIS BY DEOXYRIBONUCLEATE. Proc Natl Acad Sci U S A 44, 1072–1078 (1958).

87. Schaeffer P, Millet J, Aubert JP. Catabolic repression of bacterial sporulation. Proc Natl Acad Sci U S A 54, 704–711 (1965).

88. Sterlini JM, Mandelstam J. Commitment to sporulation in Bacillus subtilis and its relationship to development of actinomycin resistance. Biochem J 113, 29–37 (1969).

89. Koo BM, et al. Construction and Analysis of Two Genome-Scale Deletion Libraries for Bacillus subtilis. Cell Syst 4, 291–305.e297 (2017).

90. Schneider CA, Rasband WS, Eliceiri KW. NIH Image to ImageJ: 25 years of image analysis. Nat Methods 9, 671–675 (2012).

91. Bogdanov M, Zhang W, Xie J, Dowhan W. Transmembrane protein topology mapping by the substituted cysteine accessibility method (SCAM(TM)): application to lipid-specific membrane protein topogenesis. Methods 36, 148–171 (2005).

92. Stratton BS, et al. Cholesterol Increases the Openness of SNARE-Mediated Flickering Fusion Pores. Biophys J 110, 1538–1550 (2016).

93. Nikolaus J, Karatekin E. SNARE-mediated Fusion of Single Proteoliposomes with Tethered Supported Bilayers in a Microfluidic Flow Cell Monitored by Polarized TIRF Microscopy. J Vis Exp, (2016).

94. Tinevez JY, et al. TrackMate: An open and extensible platform for single-particle tracking. Methods 115, 80–90 (2017).

95. Tarantino N, et al. TNF and IL-1 exhibit distinct ubiquitin requirements for inducing NEMO-IKK supramolecular structures. J Cell Biol 204, 231–245 (2014).

96. Huet S, Karatekin E, Tran VS, Fanget I, Cribier S, Henry JP. Analysis of transient behavior in complex trajectories: application to secretory vesicle dynamics. Biophys J 91, 3542–3559 (2006).

97. Smith MB, Karatekin E, Gohlke A, Mizuno H, Watanabe N, Vavylonis D. Interactive, computer-assisted tracking of speckle trajectories in fluorescence microscopy: application to actin polymerization and membrane fusion. Biophys J 101, 1794–1804 (2011).

98. Lacombe C, Lubochinsky B. Specific extraction of bacterial cardiolipin from sporulating Bacillus subtilis. Biochim Biophys Acta 961, 183–187 (1988).

99. Angelova MI, Dimitrov DS. Liposome Electroformation. Faraday Discuss 81, 303-+ (1986).

100. Connell E, Scott P, Davletov B. Real-time assay for monitoring membrane association of lipid-binding domains. Anal Biochem 377, 83–88 (2008).

101. Sterlini JM, Mandelstam J. Commitment to Sporulation in Bacillus Subtilis and Its Relationship to Development of Actinomycin Resistance. Biochem J 113, 29-+ (1969).

102. Buser CA, Sigal CT, Resh MD, McLaughlin S. Membrane binding of myristylated peptides corresponding to the NH2 terminus of Src. Biochemistry 33, 13093–13101 (1994).

103. Ducret A, Quardokus EM, Brun YV. MicrobeJ, a tool for high throughput bacterial cell detection and quantitative analysis. Nat Microbiol 1, 16077 (2016).

104. Douglas SM, Dietz H, Liedl T, Högberg B, Graf F, Shih WM. Self-assembly of DNA into nanoscale three-dimensional shapes. Nature 459, 414–418 (2009).

105. Stahl E, Martin TG, Praetorius F, Dietz H. Facile and scalable preparation of pure and dense DNA origami solutions. Angew Chem Int Ed Engl 53, 12735–12740 (2014).

106. Chen ZM, Atefi E, Baumgart T. Membrane Shape Instability Induced by Protein Crowding. Biophysical Journal 111, 1823–1826 (2016).

107. Powers TR. Dynamics of filaments and membranes in a viscous fluid. Rev Mod Phys 82, (2010).

108. Ojkic N, Lopez-Garrido J, Pogliano K, Endres RG. Bistable Forespore Engulfment in Bacillus subtilis by a Zipper Mechanism in Absence of the Cell Wall. Plos Comput Biol 10, (2014).

