## Supplementary material for "FisB relies on homo-oligomerization and lipid-binding to catalyze membrane fission in bacteria": S2 Appendix: Theoretical modeling

### DERIVATION OF THE EQUILIBRIUM EQUATIONS

As explained in the main text, we consider the free energy $\mathscr{F}$ of an axisymmetric membrane neck of radius $R$ and length $L$ that connects two planar membrane sheets, corresponding to the local geometry depicted in **Fig A**, where the engulfment membrane meets the rest of the mother cell membrane. For simplicity, we assume the surface density $\phi$ of FisB proteins in the neck is uniform and reaches equilibrium with a surface density $\phi_0$ of FisB in the surrounding membranes. The energy functional consists of a term accounting for membrane bending and tension, $\mathscr{F}_{\mathrm{m}}$, and another term accounting for FisB protein-protein interactions, $\mathscr{F}_{\mathrm{p}}$. We employ the classical Helfrich-Canham theory [1–6] for the energy of the membrane, $\mathscr{F}_{\mathrm{m}}$, which reads

$$\mathscr{F}_{\mathrm{m}} = \int_{S_{\mathrm{n}}} \mathrm{d}S_{\mathrm{n}} \left[\frac{\kappa}{2}H^2 + \gamma\right] + \int_{S_{\mathrm{s}}} \mathrm{d}S_{\mathrm{s}}\gamma, \qquad (1)$$

where $S_{\mathrm{n}}$ and $S_{\mathrm{s}}$ are the surfaces of the membrane neck and sheets, $H$ is twice the mean curvature, $\kappa$ is the bending modulus, and $\gamma$ is the surface tension. The two membrane sheets are assumed to be planar, thus their only contribution to the energy comes from membrane tension.

Concerning FisB proteins, we include translational entropy, the energy of homo-oligomerization in trans between opposing membranes, and an energy that limits crowding. Additionally, as shown in the main text, FisB proteins do not exhibit curvature sensing, thus we do not include a term coupling FisB density to spontaneous curvature in Eq. (1). This results in the following expression for the protein free energy functional

$$\mathscr{F}_{\mathrm{p}} = \int_{S_{\mathrm{n}}} \mathrm{d}S_{\mathrm{n}} \left\{ k_{\mathrm{B}}T\phi \ln\left(\frac{\phi}{\phi_0}\right) + aV_{\mathrm{LJ}}(R)\phi^2 + U(\phi) \right\}. \qquad (2)$$

The first term accounts for translational entropy. The second term is an energy per unit area describing trans interactions of FisB, and which for simplicity is assumed to be proportional to the standard Lennard–Jones (LJ) potential accounting for a longer-range attraction and shorter-range repulsion:

$$V_{\mathrm{LJ}}(r) = \left(\frac{\sigma}{r}\right)^{12} - \left(\frac{\sigma}{r}\right)^{6}. \qquad (3)$$

The function $U(\phi)$ is an energy penalty for in-plane crowding that increases rapidly above a certain FisB concentration. To obtain $U(\phi)$ we assume a purely repulsive, truncated and shifted LJ potential between cis-neighboring FisB molecules, which we take to occupy a triangular lattice in order to relate density to nearest-neighbor distance. Therefore, $U(\phi) = \varepsilon[V_{\mathrm{LJ}}(r(\phi)) - V_{\mathrm{LJ}}(r_{\max})]$ when $r \le r_{\max}$ and 0 when $r > r_{\max}$, where we have chosen $r_{\max} = 2^{1/6}\sigma_{\mathrm{cis}}$, namely the minimum of the LJ potential with length scale $\sigma_{\mathrm{cis}}$. The result is

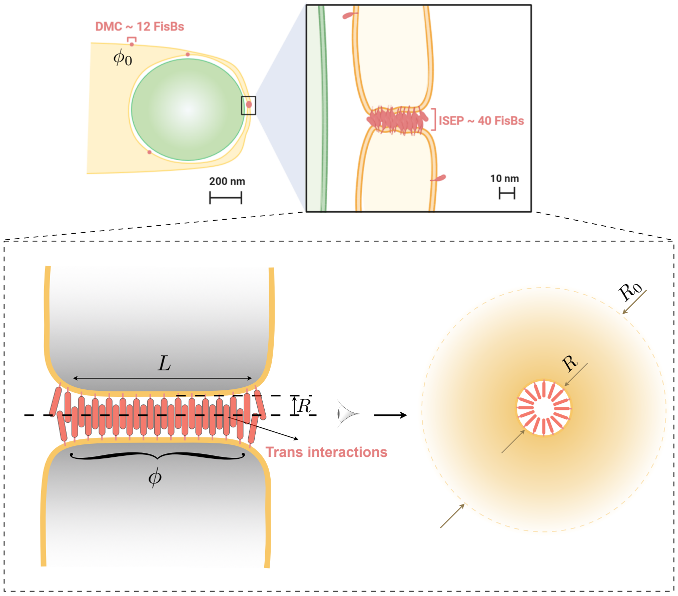


**Fig A**. Left: Sketch of the late stages of forespore engulfment, when a small membrane neck connects the engulfment membrane to the rest of the mother cell membrane. Right: Schematic of FisB accumulation at the fission site. FisB freely moves around the engulfment membrane and other regions of the mother cell membrane, forming clusters of up to $\sim$12 molecules. Bottom: Sketch of the modeled axisymmetric membrane neck of radius $R$ and length $L$ populated by a uniform surface density $\phi$ of FisB, and which connects two planar membrane sheets. The side view of the neck displays the idealized distance $R_0$ at which FisB surface density becomes $\phi_0$. (Reproduced from Panel A of Fig 9 of the main text.)

$$U(\phi) = \begin{cases} \varepsilon \dfrac{(\phi^3 - \phi^3_{r_{\max}})^2}{4\phi^6_{r_{\max}}} & \phi \ge \phi_{r_{\max}}, \\ 0 & \phi < \phi_{r_{\max}}, \end{cases} \qquad (4)$$

where $\phi_{r_{\max}} = 2^{2/3}/(3^{3/2}\sigma^2_{\mathrm{cis}}) = 2/(3^{3/2}r^2_{\max})$ is the FisB concentration corresponding to a nearest neighbor distance $r_{\max}$.

To obtain an equation for the equilibrium density of FisB proteins in the neck, we minimize $\mathscr{F} = \mathscr{F}_{\mathrm{m}} + \mathscr{F}_{\mathrm{p}}$ with respect to $\phi$,

$$k_{\mathrm{B}}T[1 + \ln(\phi/\phi_0)] + 2a\phi \left[\left(\frac{\sigma}{R}\right)^{12} - \left(\frac{\sigma}{R}\right)^{6}\right] + \partial_\phi U(\phi) = 0. \qquad (5)$$

Then, minimizing $\mathscr{F}$ with respect to $R$ yields an equation that determines the equilibrium radius of the neck

$$\gamma_{\mathrm{eff}} - \frac{\kappa}{2R^2} + a\phi^2\left(\frac{6\sigma^6}{R^6} - \frac{12\sigma^{12}}{R^{12}}\right) - \frac{2\gamma R}{L} = 0, \qquad (6)$$

where $\gamma_{\mathrm{eff}} = \gamma + k_{\mathrm{B}}T\phi \ln(\phi/\phi_0) + U(\phi)$.

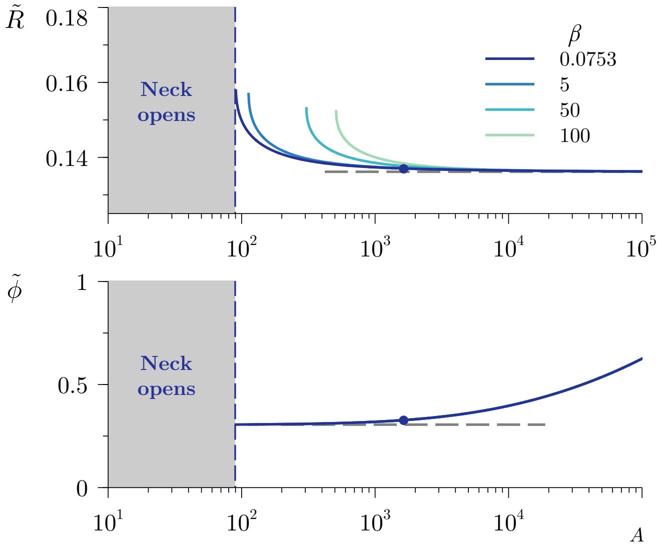


**Fig B**. Dimensionless equilibrium neck radius $R$ (top) and FisB surface density $\phi$ (bottom) as functions of the dimensionless FisB trans homo-oligomerization strength $A$, for $\Gamma = 0.149$, $\Lambda = 6.11 \times 10^{-4}$, $\bar{\varepsilon} = 200$, $\Sigma = 0.121$, and different values of $\beta$ indicated in the legend. Below a minimum interaction strength, FisB cannot stabilize the neck and the neck opens. The horizontal lines are the radius corresponding to the minimum of the potential describing the trans interaction, $R = 2^{1/6}\Sigma$, and the concentration of FisB at the onset of in-plane crowding $\phi_{r_{\max}} = 2^{2/3}/3^{3/2}$. The dot indicates the predicted equilibrium neck radius and the corresponding FisB surface density in the neck for the values of the dimensionless parameters obtained using the estimates given in **Table A**, namely $A \simeq 1.637 \times 10^3$ and $\beta \simeq 0.075$.

**Table A**. Estimates of the physical parameters

| Parameter | Value |
|---|---|
| $\kappa$ | $20 k_{\mathrm{B}}T$ [7] |
| $\phi_0$ | 100 FisB $\mu$m$^{-2}$ |
| $\gamma$ | $10^{-4}$ N m$^{-1}$ [8] |
| $\phi_{r_{\max}}$ | $5 \times 10^4$ FisB $\mu$m$^{-2}$ |
| $a$ | $10^4$ $k_{\mathrm{B}}T$ nm$^2$ FisB$^{-1}$ |
| $L$ | 40 nm |
| $\sigma \sim \sigma_{\mathrm{cis}}$ | 2.47 nm |
| $\varepsilon$ | 32.78 $k_{\mathrm{B}}T$ nm$^{-2}$ |
| $R_0$ | 1.5 $\mu$m |
| $D$ | 0.1-1 $\mu$m$^2$/s |

### DIMENSIONLESS PARAMETERS AND EQUATIONS

To reduce the parameters of the system, Eqs. (5) and (6) are non-dimensionalized by choosing $\phi_{\mathrm{c}} = \sigma_{\mathrm{cis}}^{-2}$ and $R_{\mathrm{c}} = \sqrt{\kappa/(2\gamma)}$ as the characteristic scales of FisB surface density at the neck, and the neck radius, respectively. In particular, using the estimates in **Table A**, $\phi_{\mathrm{c}} \simeq 1.637 \times 10^5$ FisB $\mu$m$^{-2}$ and $R_{\mathrm{c}} \simeq 20.28$ nm. The value of $\sigma_{\mathrm{cis}}$ is obtained from the relation $\phi_{r_{\max}} = 2^{2/3}/(3^{3/2}\sigma_{\mathrm{cis}}^2) = 2/(3^{3/2} r_{\max}^2)$. Introducing these scales into Eqs. (5) and (6) yields

$$1 + \ln\left(\frac{\tilde{\phi}}{\Lambda}\right) + 2A\tilde{\phi}\left[\left(\frac{\Sigma}{\tilde{R}}\right)^{12} - \left(\frac{\Sigma}{\tilde{R}}\right)^{6}\right] + \partial_{\tilde{\phi}}\tilde{U}(\tilde{\phi}) = 0, \tag{7a}$$

$$\Gamma\left(1 - \frac{1}{\tilde{R}^2}\right) + \tilde{\phi}\ln\left(\frac{\tilde{\phi}}{\Lambda}\right) + \tilde{U}(\tilde{\phi}) + A\tilde{\phi}^2\left[6\left(\frac{\Sigma}{\tilde{R}}\right)^{6} - 12\left(\frac{\Sigma}{\tilde{R}}\right)^{12}\right] - 2\beta\tilde{R} = 0, \tag{7b}$$

where

$$\tilde{U}(\tilde{\phi}) = \begin{cases} \dfrac{\tilde{\varepsilon}}{192}(4\sqrt{3} - 243\tilde{\phi}^3)^2 & \tilde{\phi} \geq \dfrac{2^{2/3}}{3^{3/2}}, \\ 0 & \tilde{\phi} < \dfrac{2^{2/3}}{3^{3/2}}, \end{cases} \tag{8}$$

where $\tilde{\phi}$ and $\tilde{R}$ denote the dimensionless versions of FisB surface density and neck radius, respectively. The dimensionless parameters in Eqs. (7a) and (7b) are

$$A = \frac{a\phi_{\mathrm{c}}}{k_{\mathrm{B}}T}, \quad \Sigma = \frac{\sigma}{\sqrt{\kappa/(2\gamma)}}, \quad \tilde{\varepsilon} = \frac{\varepsilon\sigma_{\mathrm{cis}}^2}{k_{\mathrm{B}}T}, \quad \Lambda = \frac{\phi_0}{\phi_{\mathrm{c}}},$$
$$\Gamma = \frac{\gamma}{k_{\mathrm{B}}T\phi_{\mathrm{c}}}, \quad \beta = \frac{\gamma\sqrt{\kappa/(2\gamma)}}{k_{\mathrm{B}}T\phi_{\mathrm{c}}L}. \tag{9}$$

Taking the values from **Table A**, $\Gamma \simeq 0.149$, $\Lambda \simeq 6.11 \times 10^{-4}$, $\beta \simeq 0.075$, $A \simeq 1.637 \times 10^3$, $\tilde{\varepsilon} \simeq 200$, and $\Sigma \simeq 0.121$.

### CONDITIONS FOR A STABLE NECK RADIUS IN THE PRESENCE AND IN THE ABSENCE OF FISB

**Fig B** shows $\tilde{R}$ and $\tilde{\phi}$ as functions of the dimensionless FisB trans homo-oligomerization strength $A$, for different values of $\beta$ indicated in the legend. In particular, fixing the material parameters given in **Table A**, $\beta$ is an inversely-proportional function of the neck length $L$ alone. Indeed, for a realistic estimate of the material parameter values (the dot), we find that FisB trans interactions are strong enough to stabilize the neck at $\tilde{R} \sim 0.14$, i.e. $R = 2.84$ nm, with a close-packed concentration of FisB in the neck $\tilde{\phi} \sim 2^{2/3}/3^{3/2}$, i.e. $\phi = 0.05$ FisB nm$^{-2}$.

We also show that there is a critical lower limit of $A$ below which the FisB trans interactions are too weak to stabilize the neck, thus the neck opens ($\tilde{R} \to \infty$) in our simplified model. This critical value of $A$ depends on $\beta$, namely the larger the value of $\beta$ (the shorter the neck), the stronger the trans interactions needed to stabilize the neck at a finite radius. This makes intuitive sense: the longer the neck, the more FisB can be present to hold the neck together in opposition to membrane tension, which tends to make the neck expand.

We note that even in the absence of FisB, for a long enough neck there is a metastable state of the neck at finite radius $R$. This reflects a balance between membrane bending energy,

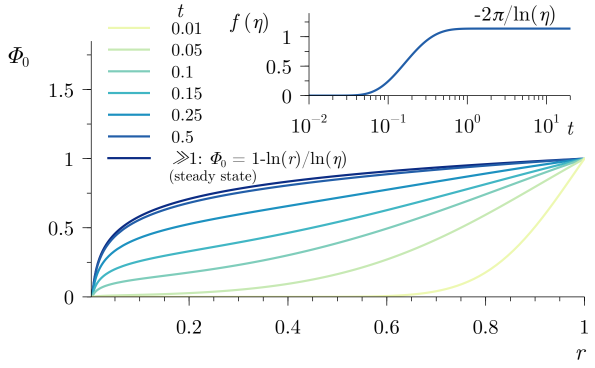


**Fig C**. Concentration of FisB, $\Phi_0(r)$ at the membrane sheet as a function of the radial coordinate $r$, for $\eta = 4\times10^{-3}$. The inset shows the evolution of the FisB flux at the neck radius $r=\eta$ over time.

which always favors larger $R$, and surface tension, which contributes non-mononotically to the membrane energy as a function of $R$. For small $R$ and large $L$, increasing the radius $R$ increases the total amount of membrane in the vicinity of the neck and so increasing $R$ is opposed by surface tension. By contrast, for large $R$, increasing $R$ removes more membrane from the parallel sheets than is added to the neck, since the former scales $\sim R^2$ while the latter scales $\sim LR$, so surface tension favors further increase of $R$. To illustrate this effect mathematically we analyze the membrane free energy functional in the absence of FisB proteins, which reads

$$\mathscr{F}_{\rm m} = R - \frac{R^2}{L/R_{\rm c}} + \frac{1}{R}. \tag{10}$$

For lengths below a critical value $L < 3^{3/2}R_{\rm c}$, the above energy functional does not have a real minimum and expanding the radius of the neck decreases the total energy of the system, implying the neck opens. By contrast, for $L > 3^{3/2}R_{\rm c}$, $\mathscr{F}_{\rm m}$ in Eq. (10) has a local minimum, implying a metastable finite neck radius. For the sake of conciseness the lengthy expression for the metastable radius is not shown here.

### MINIMUM STABLE NECK LENGTH

In this section we compute the critical $\beta$ (i.e. neck length) at which the stable neck represented by the dot in **Fig B** opens for realistic parameter values. For a value of $A \simeq 1.637\times10^3$, the opening of the neck occurs at $\beta \simeq 402$, which, taking the estimates from **Table A**, corresponds to a value of $L$ below 1 Å. This means that for realistic values of $L$ and the estimates given in **Table A**, our simple model always predicts the membrane neck can be stabilized by FisB trans interactions.

### DIFFUSION AND CAPTURE TIME ESTIMATE

While the above results suggest that an accumulation of FisB at the neck can be energetically stable, we now consider how long it might take to reach that state. Here, we first analyze the time needed for FisB monomers in the surrounding membranes to diffuse and be captured at the neck.

To estimate the time required for a certain number of FisB proteins to be absorbed by the neck, we consider purely-diffusive dynamics within a planar membrane connected to the neck. In particular, we assume an annular domain, $\mathscr{V}$, depicted in **Fig A** (bottom right), whose FisB concentration $\Phi_0(\boldsymbol{x},t)$ is given by the diffusion equation, which is made dimensionless employing

$$\phi_{0,\rm c} = \phi_0, \quad \ell = R_0, \quad t_{\rm c} = \frac{R_0^2}{D} \tag{11}$$

as characteristic scales for the areal FisB concentration, length, and time, respectively. Here $\phi_0$ is the concentration of FisB in the sheet, assumed to be held constant at a certain radius $R_0$ far from the neck, and $D$ is the diffusion coefficient of FisB. The dimensionless diffusion equation then reads

$$\partial_t \Phi_0 = \boldsymbol{\nabla}^2\Phi_0 \quad \boldsymbol{x}\in\mathscr{V}, \tag{12}$$

with $\boldsymbol{x} = (r,\theta)$ denoting the position vector. We impose an absorbing boundary condition at the neck radius $R$, and a fixed value of $\Phi_0$ at the outer radius $R_0$,

$$\Phi_0 = 0 \quad \text{at } r=\eta, \quad \text{and} \quad \Phi_0 = 1 \quad \text{at } r=1, \tag{13}$$

where $\eta = R/R_0$ is the inner-to-outer radii ratio. Concerning the initial condition, as a conservative choice we impose $\Phi_0(t=0)=0$ everywhere except at the outer radius, where $\Phi_0(t=0, r=1)=1$. The solutions can be found using, for example, a Fourier expansion involving Bessel functions (not shown here for the sake of brevity). The axisymmetric time-dependent solution is shown in **Fig C** at different times indicated in the legend. The inset displays the flux at $r=\eta$ as a function of time, from which we can obtain an estimate of the absorption time of a specified number of FisB proteins at the neck. At steady state, the concentration of FisB proteins in the annulus and the corresponding flux at the neck, $r=\eta$, read

$$\Phi_0(r, t\gg1) = 1 - \frac{\ln(r)}{\ln(\eta)}, \quad f(t\gg1) = -\frac{2\pi}{\ln(\eta)}. \tag{14}$$

Using the estimates given in **Table A**, where we have considered $R_0 \simeq 1.5$ $\mu$m, the characteristic diffusion time is $t_{\rm c} = R_0^2/D \simeq$ 2.25-22.5 s, and the ratio of radii is $\eta \simeq 4\times10^{-3}$. The steady-state flux then yields, $f \simeq 1.126$, i.e. 10 FisB s$^{-1}$. Using the time-dependent evolution of the flux, the time required for diffusion and capture ∼40 FisB proteins in the neck is ∼4 s, which is very short and thus not expected to be rate limiting.

### NUCLEATION TIME OF FISB CLUSTERS AT THE NECK

To obtain a simple estimate of the nucleation time of a stable cluster of FisB proteins at the neck for both low-expression and native-expression strains, we assume that FisB proteins diffuse independently on the entire membrane and that nucleation of stable a cluster in the neck occurs when $n$ proteins happen to be in the neck at the same time. To this end, we need to estimate the fraction of time there are $n$ or more FisB proteins in the neck, as well as the correlation time, that is the time between uncorrelated samples. Since we assume FisB proteins are independent, the number of proteins in the neck will be Poisson distributed, thus we only need to know the average number of FisB in the neck to obtain the complete distribution. The average can be estimated as the neck area, $2\pi RL$, times the background concentration, $\phi_0$. Furthermore, the correlation time is simply the time for a FisB to diffuse the length of the neck, $L^2/D$. Using the the values given in **Table A**, and taking the concentration $\phi_0 \simeq 20$ FisB $\mu\text{m}^{-2}$ for the low-expression strain, yields $\langle \text{FisB} \rangle \simeq 0.03$ in the neck. Assuming that the $\sim$1-hour delay in membrane fission during sporulation observed in the low-expression strain is due to the time for nucleation, we can infer that the number of FisB proteins required for nucleation is $n = 3$, which yield a nucleation time of $t_\text{N} = \tau/\text{Pois}(3, \langle \text{FisB} \rangle) \simeq 60$ min. If the native-expression strain also needs $n \approx 3$ FisB proteins to nucleate, we can estimate its corresponding nucleation time using the value given in **Table A**, $\phi_0 \simeq 100$ FisB $\mu\text{m}^{-2}$, which yields $\langle \text{FisB} \rangle = 0.151$ and a nucleation time of $t_\text{N} \simeq 30$ s.


[1] P. B. Canham, J. Theor. Bio. **26**, 61 (1970).
[2] W. Helfrich, Z. Naturforsch. C. **28**, 693 (1973).
[3] O. Y. Zhong-Can and W. Helfrich, Phys. Rev. Lett. **59**, 2486 (1987).
[4] O.-Y. Zhong-Can and W. Helfrich, Phys. Rev. A **39**, 5280 (1989).
[5] U. Seifert and S. A. Langer, EPL **23**, 71 (1993).
[6] U. Seifert, Adv. Phys. **46**, 13 (1997).
[7] T. R. Powers, Rev. Mod. Phys. **82**, 1607 (2010).
[8] N. Ojkic, J. López-Garrido, K. Pogliano, and R. G. Endres, PLoS Comput. Biol. **10**, e1003912 (2014).
