## Supplemental Movies for "FisB relies on homo-oligomerization and lipid-binding to catalyze membrane fission in bacteria": README.txt

Movies are presented in two separate formats:

1) As tiff stacks (with extension .tif). These are the raw, 16-bit images that can be opened using the freely available programs ImageJ (https://imagej.nih.gov/ij/) or Fiji (https://imagej.net/Fiji). 

2) As .avi movies. The .avi movies will play at 10 images/sec, 10-fold accelerateed compared to the original movies.
