## Supplementary figures and images for "FisB relies on homo-oligomerization and lipid-binding to catalyze membrane fission in bacteria"

### Movie1_FisB.tif

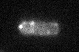

### Movie2_FisB.tif

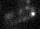

### Movie3_GFP_FisB.tif

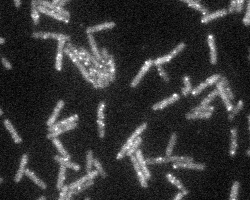

### Movie4_GFP_FisB.tif

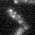

### Movie5_GFP_Mbl.tif

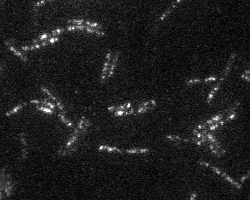

### Movie6_GFP_Mbl.tif

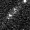

### Movie7_GFP_Mbl_fos.tif

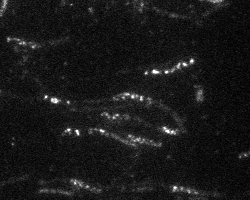

### Movie8_GFP_Mbl_fos.tif

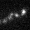

### Movie9_GFP_FisB_fos.tif

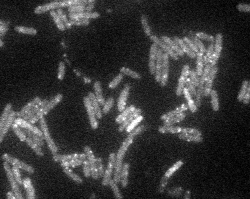

### Movie10_GFP_FisB_fos.tif

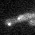
