## Supplementary material for "FisB relies on homo-oligomerization and lipid-binding to catalyze membrane fission in bacteria": S1 Appendix: Supplementary Experiments

### S1 Appendix: Supplementary Experimental Information

#### S1 APPENDIX: RESULTS

##### Quantification of FisB copy numbers using *B. subtilis* fluorescence standards and Western blotting.

We used two independent approaches to quantify FisB copy numbers in addition to the DNA-origami based quantification presented in the main text. First, we used *B. subtilis* calibration strains[1] to relate the total cell fluorescence (sum of pixel values) to copy numbers of super-folder GFP (sfGFP). Cells expressing known copy numbers of sfGFP were imaged in wide-field fluorescence and the total intensity, corrected for background, was plotted against sfGFP copy number (S1 Appendix Fig. 2A,B). We then imaged cells expressing an equivalent number of sfGFP (strain SG13) or mEGFP (strain BAL038) molecules and found the total fluorescence was 2.4±0.2-fold higher for sfGFP (S1 Appendix Fig. 2C). Using this correction factor, the calibration for sfGFP was converted to a calibration for mEGFP, the label used for imaging FisB (S1 Appendix Fig. 2D). We then computed the distribution of total cell fluorescence in *B. subtilis ΔfisB* cells expressing mEGFP-FisB (BAL001) at native levels at t=3 h into sporulation (S1 Appendix Fig. 2E). We also estimated the total fluorescence of DMC and the ISEP as a fraction of total cellular fluorescence intensity (S1 Appendix Fig. 2F). Together with the calibration of S1 Appendix Fig. 2D, the means of these distributions allowed us to estimate ~1300 total FisB copies per cell, of which ~50 are located at the ISEP. The DMCs contain ~16 copies of FisB on average by this estimate.

Second, we used quantitative Western blotting (WB) to relate the total number of FisB molecules per cell to fluorescence intensity. Purified recombinant mYFP was used to calibrate the average number of mGFP-FisB molecules per cell using WB, using anti-GFP antibodies (S1 Appendix Fig. 3). This average number per cell (~700) was then used to calculate mGFP-FisB copies at the ISEP (~30 copies) and DMC (~10 copies) from the fluorescence measurements in S1 Appendix Fig. 2F, the fraction of total cellular fluorescence signal found at ISEP or DMC.

##### Localization of FisB is not coupled to cell wall remodeling, the protonmotive force, or the membrane potential

The sub-cellular localization and motion of many cellular components depend on the cell-wall remodeling machinery[2-4], the protonmotive force and the membrane potential[5]. We tested if any of these influenced dynamics of FisB.

Cell wall synthesis and degradation drive engulfment during sporulation[2,6]. It was suggested that cell wall remodeling might also drive fission of the engulfing membrane at the end of engulfment. We wondered whether FisB dynamics could be coupled to cell wall remodeling. Since inhibition of cell wall synthesis leads to engulfment defects, we expressed mGFP-FisB from an inducible promoter during vegetative growth and investigated effects of inhibition of cell wall synthesis by fosfomycin on the motion of FisB clusters. As a control, we chose to image GFP-Mbl in parallel experiments. Mbl is an actin homologue that controls cell wall synthesis and cell shape[7]. Mbl forms filaments that are associated with the cell membrane and rotate around the cell circumference together with enzymes required for cell wall synthesis[8]. Mbl filaments stop moving and eventually disassemble upon treatment with fosfomycin[9] which inhibits the formation of N-acetylmuramic acid, a building block of the bacterial cell wall[10].

We imaged cells expressing either mGFP-FisB (BMB014) or GFP-Mbl (BDR2061) using total internal reflection fluorescence microscopy (TIRFM) in the absence and presence of fosfomycin (S1 Appendix Fig. 6). In this imaging mode, only fluorophores within ~100 nm of the glass-aqueous buffer interface are detected[11]. That is, only the spots near the substrate-proximal side a cell would be visible. Small mGFP-FisB spots, similar to the ones present during early stages of engulfment, moved around in the cell membrane seemingly randomly (S1 Appendix Movies 3 and 4). By contrast, GFP-Mbl spots moved along the short axis of the cell (S1 Appendix Movies 5 and 6), as previously reported[8,12]. To quantify these motions, we used four different approaches. First, we computed kymographs of mGFP-FisB or GFP-Mbl along the short and long axes of a cell, as shown in S1 Appendix Figure 6A, C. Before treatment, GFP-Mbl moved around the cell circumference, reflected by stripes across the cell in the maximum intensity projections (MIP) and spots that appear and disappear in the kymographs along the long axis (marked with a red frame). Spots also appear and disappear along the short axis as GFP-Mbl spots move in and out of the evanescent field as they move along the cell circumference (kymographs marked by blue frames). Addition of fosfomycin stopped the motion of GFP-Mbl (S1 Appendix Movies 7 and 8), resulting in small spots in the MIP and continuous lines in the kymographs. By contrast, mGFP-FisB MIPs and kymographs were not appreciably modified upon fosfomycin treatment (S1 Appendix Movies 9 and 10). Second, we tracked individual GFP-Mbl and mGFP-FisB spots and calculated the mean-squared displacement (MSD) (S1 Appendix Figure 6B,D). Addition of fosfomycin reduced the motility of GFP-Mbl filaments (S1 Appendix Figure 6B), whereas the motion of mGFP-FisB was unaffected (S1 Appendix Figure 6D). Third, the average total distance that GFP-Mbl filaments traveled within 3 s was reduced in the presence of fosfomycin, whereas no such effect was found for the distance traveled by mGFP-FisB spots (S1 Appendix Figure 6E). Finally, we computed the asymmetry[13] of the individual GFP-Mbl and mGFP-FisB trajectories, and computed averages (S1 Appendix Figure 6F). Asymmetry is a measure of the tendency for a persistently preferred direction of motion: asymmetry is zero for a perfectly symmetric trajectory, whereas it diverges for a straight-line trajectory. For simulated 2-dimensional Brownian trajectories, asymmetry rapidly converges to ~0.26 for a large number of steps and/or particles. The asymmetry of GFP-Mbl trajectories before fosfomycin treatment was high, equal to $0.58 \pm 0.06$ (mean ± SEM), consistent with previous reports of Mbl moving in linear tracks[8]. Upon treatment, GFP-Mbl spots stopped moving, and the asymmetry decreased to $0.30 \pm 0.07$. By contrast, the asymmetry of mGFP-FisB trajectories before ($0.22 \pm 0.03$) and after treatment ($0.23 \pm 0.06$) were similar (S1 Appendix Figure 6F). Thus, the motion of FisB clusters is independent of cell-wall synthesis.

The protonmotive force (PMF) is important for the localization of proteins that are involved in maintaining cell shape, such as MreB and Mbl, or cell division (e.g. FtsZ/FtsA)[5]. We tested whether the localization of FisB depends on the PMF by imaging mGFP-FisB (BAM003) during sporulation in the absence and presence of carbonyl cyanide m-chlorophenyl hydrazone (CCCP), a proton-ionophore that dissipates the membrane PMF. We found that the localization of GFP-FisB at t=3 h into sporulation is not affected by the PMF unlike the localization of Mbl (S1 Appendix Figure 6G).

Proteins whose localization depends on the PMF also require an intact membrane potential. To see if the membrane potential affected FisB dynamics, we imaged mGFP-

FisB (BAM003) in the presence and absence of valinomycin, an antibiotic that functions as a potassium carrier that depletes the transmembrane electric potential component of the PMF. We found that GFP-Mbl mislocalizes in the presence of valinomycin, whereas the localization of FisB is not affected (S1 Appendix Figure 6G).

Together, these results show that the dynamic movement of FisB is independent of cell wall remodeling, the PMF, and the membrane potential.

#### Topology of FisB

The FisB protein family is defined through a consensus region (residues 130-223, Figure 4A) identified in Pfam[14]. Most algorithms predict FisB to possess a single transmembrane domain (TMD) with a small N-terminal cytoplasmic domain and a larger (23-kDa) extracellular domain (ECD), as depicted Figure 4A. However, some algorithms assign a hydrophobic region around residues 130-150 as a second TMD, or even predict an inverted topology (S1 Appendix Figure 7B). If there were a second TMD (S1 Appendix Figure 8C), or if the topology were inverted, the C-terminus would face the cytosol rather than the extracytoplasm. To determine FisB's topology, we tested accessibility of cysteine residues introduced at various positions to a membrane impermeable, biotinylated, sulfhydryl-reactive reagent, 3-(N-maleimidylpropionyl) biocytin (MPB)[15]. We generated three Myc-tagged FisB mono-cysteine variants (Myc-tagged FisB G6C, FisB L137C and FisB A245C) and tested separately whether these cysteines were intra-or extracellular. When complementing ΔfisB cells, these mutations reduced the sporulation efficiency slightly (S1 Appendix Table 2), we assume without an effect on topology. We first tested cysteine accessibility to MPB in lysed protoplasts of sporulating cells. Membranes were solubilized with detergent and FisB was pulled down using a polyclonal anti-myc antibody. Biotinylation was probed by Western Blot using an HRP-conjugated-avidin antibody. This analysis showed that only residues 6 and 245 were accessible (S1 Appendix Figure 8D left panel, top row), suggesting residue 137 may be restricted by secondary/tertiary structures and/or the membrane. In intact protoplasts only residue 245 was labeled by MPB, indicating the C-terminus faces the extracellular space (S1 Appendix Figure 8D middle panel). In contrast, only residue 6 was biotinylated when extracellular cysteines were blocked by 4-acetamido-4′-maleimidylstilbene-2,2′-disulfonic acid (AMS) prior to cell lysis and incubation with MPB. That is, residue 6 faces the cytoplasm (S1 Appendix Figure 8D right panel). To confirm the presence of FisB, we stripped every Western Blot and probed it with anti-FisB antibody (S1 Appendix Figure 8D, bottom row). Together, these results are consistent with a topology in which the larger C-terminus of FisB is extracellular, the N-terminus faces the cytoplasm, and residue 137 is inaccessible to biotinylation, possibly residing inside a globular domain, shielded at the oligomerization interface, or by the membrane. Our attempts to determine the structure of FisB ECD were unsuccessful, but a computational model of FisB for residues 44 to 225, covering most of the ECD is available[16] and is shown in Figure 4B. The model predicts a curved ECD structure, with ~3 nm and ~5 nm for the inner and outer radii of curvatures. The overall topology of FisB, with the predicted ECD structure is schematically shown Figure 4B.

#### S1 APPENDIX FIGURE LEGENDS

**S1 Appendix Figure 1. Membrane fission assay. A**. Membrane fission as a function of time after downshift to sporulation medium. Aliquots were taken from sporulating suspensions of wild-type (PY79) or FisB null mutant cells (BDR1083) at the indicated times, labeled with the membrane dye TMA-DPH, mounted on agarose pads containing resuspension medium, and imaged using a wide-field fluorescence microscope. The dye only partially crosses the membrane, resulting in strong labeling of the forespore contour pre-fission and dim labeling post-fission. Bar,1 µm. **B.** The original membrane fission assay using the lipophilic membrane dye FM4-64 and expression of a forespore marker[17]. The dye is virtually non-fluorescent in the medium, and it cannot cross the cell membrane. Thus, before fission, FM4-64 labels the outer leaflet of both the mother cell and the forespore membranes. After fission, only the outer leaflet of the mother cell is labeled. Because post-fission cells and cells that never entered sporulation are labeled in the same manner, in addition to FM4-64, a fluorescent protein is expressed in the forespore under the control of the forespore-specific transcription factor $\sigma^F$ to distinguish between the two cell types[6]. This makes it challenging to monitor FisB dynamics simultaneously, which requires a third channel. FM4-64 was added to an aliquot from a sporulating culture of *B. subtilis* cells (BKM15), then cells were mounted onto an agarose pad and imaged. The cell in the top image completed engulfment, but not fission, because the dye had access to the space around the forespore (FS). For the cells in the middle and bottom, the dye did not have access to the space around the FS. This could be because a cell never entered sporulation, or because of successful membrane fission. Expression of a FS reporter (PspoIIQ-cfp, green) is required to distinguish between the two possibilities. For the cell in the middle row, the FS marker is present, indicating the cell successfully underwent membrane fission. The cell in the bottom row never entered sporulation. **C**. Kinetics of membrane fission during sporulation, monitored using either FM4-64 or TMA-DPH. The two dyes result in indistinguishable kinetics. **D-G**. Quantification of TMA-DPH intensities for categorization of cells. Forespore contours were detected using JFilament[18]. A MATLAB script was then used to determine the mean fluorescence intensity along individual contours (see Materials and Methods). **D**. A snapshot of PY79 cells labeled with TMA-DPH at t=2.5 h after downshifting into nutrient-poor medium. The same image is shown on the right, with detected forespore contours overlaid. **E.** Distribution of mean contour intensity values from the image in D. The distribution is bimodal. A fit to a sum of two Gaussian functions is shown, with means = 349, 459 a.u. and std. dev. = 35, 23, a.u. respectively. Cells having lower intensity values were classified as having undergone fission (red, 54/124=40%), while cells with higher intensity values (blue, 62/124=54%) were classified as "no fission". Eight cells (8/124=6%) displaying intermediate values were not classified. **F.** The same as in D, but cells were labeled at t=3 h after nutrient downshift. **G.** Distribution of mean contour intensity values from the image in F. A double Gaussian fit yielded means = 333, 461 a.u. and std. dev. = 18, 22, a.u. respectively. Out of the 133 cells in F, 70% (93/133) were classified as having undergone fission, 23% (31/133) as "no fission", and 7% (9/133) as intermediate. For the data shown in Figs. 1F, 3D, and 6D, the same analysis pipeline was used, but multiple view fields were analyzed from three independent experiments.

**S1 Appendix Figure 2. Calibration of fluorescence intensity as a function of GFP copy number using *B. subtilis* calibration strains. A.** Representative images of cells expressing known average copy numbers of sfGFP[1]. **B**. The total intensity (sum of pixel values) per cell, plotted against copy number, after selecting cell contours using MicrobeJ software and subtracting background. Background was defined as total intensity (sum of pixel values) in wildtype cells which did not express any fluorescent protein. Data is plotted on semi-logarithmic coordinates in the inset, to show low copy number intensities. **C**. *B. subtilis* cells expressing the same average number of mEGFP (BAL038) or sfGFP (SG13) were imaged to determine the correction factor for mEGFP-FisB copy number quantification. Top: representative images using the same acquisition and display settings, bottom: quantification. On a per molecule basis, sfGFP is ~2.4±0.2-fold brighter than mEGFP. Strains were made by transforming PY79 cells with either plasmid pECE321 (encoding amyE::(Pveg_R0_sfGFP_spec)) or its variant wherein sfGFP was replaced by mEGFP (pECE321_mEGFP). On average $3.00\times10^5$ GFP copies/cell are expressed[1]. **D**. The calibration in B, rescaled with the correction factor determined in C, such that the calibration now corresponds to the copy numbers for mEGFP (the best-fit line constrained to pass through the origin has slope $22.48$ ($R^2 = 0.93$). **E.** Distribution of background-corrected total fluorescence intensity (sum of pixel values) for cells expressing mEGFP-FisB (BAL001) that have undergone membrane fission at t=3h. Membrane fission was assessed as in Fig. 1 and S1 Appendix Fig. 1. Background was defined as in B. N=250 cells. **F**. The total intensity of ISEP and DMCs as a percentage of total cell FisB fluorescence. Using the calibration for whole-cell fluorescence shown in D, ~1300 copies of FisB per cell; ~50 FisB/ISEP and ~16 FisB/DMC are estimated. Scale bars in A, C, represent 1 µm.

**S1 Appendix Figure 3. FisB copies per cell, estimated from Western blot analysis**. **A.** Calibration of nanograms of YFP vs. WB band intensity. Top: purified mYFP was diluted in sporulation medium to indicated ng/lane, detected using chemiluminescence using an anti-GFP antibody (ab13970) and analyzed by densitometry. Bottom: amount of YFP (in ng) vs. integrated intensity of WB band (mean ± SEM of three independent experiments). Linear regression (dotted line, forced through the origin), with $y = 2783x$ ($R^2 = 0.98$). **B.** WB detection of mXFP-FisB (where X is Y or G) loaded from a known number of cells. Solubilized membranes of wild type (PY79), *ΔfisB* (BDR1083), *ΔfisB* expressing mYFP-FisB (BVS001) or mGFP-FisB (BAM003) at 3 h into sporulation, and standards of purified mYFP were separated by SDS-polyacrylamide gel electrophoresis, subjected to immunoblotting with an anti-GFP antibody and analyzed by densitometry. Each lane was loaded with $1.44 \times 10^7$ cells, determined by serial dilutions and counting under the microscope. The average intensity of the mGFP-FisB and mYFP-FisB bands from the membrane fraction corresponds to $0.6 \pm 0.05$ ng of mXFP-FisB in A, which in turn corresponds to $(6.61 \pm 0.49) \times 10^9$ molecules per lane. Thus, on average there are 459±34 mXFP-FisB molecules/cell. However, some cleaved XFP (on average 44% of the mXFP-FisB membrane fraction signal) is detected in the soluble fraction, likely reflecting degradation of mXFP-FisB either in cells or during sample processing. Correcting for this, we estimate 661±49 mXFP-FisB molecules/cell.

**S1 Appendix Figure 4. FisB copy numbers in the low expression strain. A.** Representative fluorescence images of cells expressing mEGFP-FisB under native (BAL001) or low (BAL004) expression levels. Cell contours (yellow outline) were calculated using MicrobeJ. Fission in each case was determined as in Fig. 1C and S1 Appendix Fig. 1. Due to the large difference in pixel values, images of native and low expression strains are displayed with different brightness settings. **B.** Comparison of total copies of FisB under native or low expression levels. Sum of pixel values (total mEGFP fluorescence) per cell was calculated with MicrobeJ within the cell contours as shown in A, after background correction. The calibration in Fig. 2D was used to estimate copies per cell. Low-expression cells have 122±51 copies of mEGFP-FisB on average, or ~8-fold lower than native levels. **C.** Same as in A, with ISEP circled. ISEP were detected semi-automatically using SpeckleTrackerJ[19]. **D.** Distributions of total fluorescence intensities (sum of pixel values) for ISEP for the native and low expression strains. For each spot the sum of all pixel values in a 6×6 pixel (0.5 $\mu$m × 0.5 $\mu$m) box around the center of the cluster detected as in C was integrated. The same operation was performed at a membrane area where no clusters were present, and this background value was subtracted from the FisB cluster intensity. The integrated, background-corrected intensity values at the ISEP were compared directly with the calibration in Fig. 2D. The distribution for native expression cells is copied from Fig. 2C for comparison. Low expression cells have ~6 FisB at ISEP at the time of fission. **E.** The total intensity of ISEP as a percentage of total cell fluorescence for native and low expression cells. **F**. Immunoblot analysis of whole-cell lysates from sporulating cells. FisB levels were analyzed using an anti-FisB antibody in sporulating cells from wild-type (strain PY79), *ΔfisB* (BDR1083) and FisB-null cells expressing mYFP-FisB in low levels (strain BAL002). Time (in hours) after initiation of sporulation is indicated. **G.** Summary of FisB copy number quantification. Top. Under native expression, on average, there are ~1,000 FisB molecules per cell at t=3 h. Each DMC contains ~12 FisB molecules, while the ISEP contains ~40 FisB copies. In the low-expression strain all the numbers are scaled down ~7-8-fold.

**S1 Appendix Figure 5. Sporulation efficiency of various *B. subtilis* strains.** Sporulation efficiency was assayed by measuring heat-resistant (80°C, 20 min) colony forming units (see Materials and Methods) and normalized to the wild-type level (% of WT) for the indicated strains. Results are shown as means ±SD for four replicates per condition.

**S1 Appendix Figure 6. Motion of FisB clusters is not coupled to cell wall synthesis, or pH and voltage gradients across the cell membrane. A.** Representative TIRFM images of cells expressing GFP-Mbl (BDR2061) before and after treatment with fosfomycin. Red and light blue lines indicate the directions along the long and short axes of the cell used to compute the kymographs on the right. Before treatment, GFP-Mbl moved around the cell circumference, reflected by stripes across the cell in the maximum intensity projections (MIP) and spots that appear and disappear in the kymographs along the long axis (marked with a red frame). Spots also appear and disappear along the short axis as GFP-Mbl spots move in and out of the evanescent field as they move along the cell circumference. Addition of fosfomycin stopped the motion of GFP-Mbl, reflected in small spots in the MIP and continuous lines in the kymographs. **B**. Mean-squared displacement (MSD) as a function of lag time for GFP-Mbl before (24 tracks) and after (20 tracks) fosfomycin treatment. Colored lines connect averaged points, whereas gray areas represent standard deviation and error bars represent the standard error of the mean (SEM). Movies were acquired at 1 frame/s. The short-time diffusion coefficient, estimated from a parabolic fit to the MSD, was $D_{Mbl} = 505\,nm^2/s$ (95% confidence interval CI=439-571 $nm^2/s$) and $D_{Mbl}^{fos} = 112\,nm^2/s$ (CI=79-146 $nm^2/s$) before and after fosfomycin treatment, respectively. **C**. Representative TIRFM images of cells expressing mGFP-FisB (BMB014). Motion of GFP-FisB was not affected by addition of fosfomycin. **D**. MSD as a function of lag time for GFP-FisB before (18 tracks) and after (12 tracks) fosfomycin treatment. Acquisition rate was 1 frame/s. The short-time diffusion coefficient was $D_{FisB} = 6270\,nm^2/s$ (95% confidence interval CI=5810-6740 $nm^2/s$) and $D_{FisB}^{fos} = 6370\,nm^2/s$ (CI=5580-7160 $nm^2/s$) before and after fosfomycin treatment, respectively. **E**. Average total distance traveled by GFP-Mbl and mGFP-FisB spots over 3 s in the presence and absence of fosfomycin. GFP-Mbl (20 tracks), GFP-Mbl + fosfomycin (24 tracks), mGFP-FisB(18 tracks) and mGFP-FisB + fosfomycin (12 tracks). Fosfomycin decreased the total distance traveled by Mbl filaments ($p = 0.024$, Student's t-test), whereas FisB was not affected ($p = 0.433$). **F**. Average asymmetry of the Mbl and FisB trajectories. Upon treatment with fosfomycin, GFP-Mbl filaments stop moving, which is reflected as a decrease in asymmetry ($p = 0.0044$), whereas mGFP-FisB's motion is unaffected ($p = 0.8655$). **G.** Localization of GFP-Mbl (BDR2061) during vegetative growth and mGFP-FisB (BAM003) at t=3h into sporulation in the presence or absence of 100 μM CCCP or 30 μM valinomycin. GFP-Mbl mislocalizes in the presence of either drug, whereas the localization of mGFP-FisB is unaffected. Scale bar is 3 μm.

**S1 Appendix Figure 7. Conservation and predicted topology of FisB**. **A**. Conservation of FisB amino acid sequences derived from alignment of 250 FisB sequences from the SwissProt database, using the program ConSurf version 3.0[20]. **B.** Membrane protein topology prediction from 10 different algorithms, and the consensus prediction by Constrained Consensus Topology Prediction Server (CCTOP[21]).

**S1 Appendix Figure 8**. **Domain structure and topology of *B. subtilis* FisB. A.** Predicted domain structure of FisB. Pfam[14] identifies a consensus region (residues 129-223) defining the FisB protein family. **B**. Kyte-Doolittle hydrophobicity profile of the FisB sequence, with a potential second TMD indicated **C**. Possible topologies of FisB. Left: a single TMD with a cytoplasmic N-terminus and extracellular C-terminus. Right: With two TMDs, both the N- and the C-termini should be cytoplasmic. Cysteine residues introduced at positions 6 or 245 are indicated. **D**. Accessibility of the cysteines at positions 6, 137, and 245 to a biotinylated, sulfhydryl-reactive compound, 3-(N-maleimidoypropionlyl) biocytin (MPB). Myc-tagged monocysteine FisB variants were produced in *ΔfisB* cells and reacted with MPB before or after blocking extracellular cysteines with 4-acetamido-4'-maleimidylstilbene-2,2'-disulfonic acid (AMS). FisB was pulled down using an anti-myc antibody and biotinylation was probed by Western blot using an HRP-conjugated avidin antibody. Lysed cells were probed to ensure accessibility of MPB to the cysteine labels. The results are consistent with the amino and carboxy termini being intra- and extracellular, respectively.

**S1 Appendix Figure 9. His6-FisB ECD forms soluble aggregates *in vitro* and binds acidic membranes mainly through electrostatic interactions. A.** Schematic domain structure of the 6His-tagged FisB construct comprising the soluble extracytoplasmic domain (ECD), generated for recombinant protein purification. **B.** Second and third elution fractions from the affinity column were analyzed by SDS-PAGE and stained with SpyroOrange. The red arrow indicates the monomeric form of His6-FisB (23 kDa) and the blue bracket highlights SDS-resistant $His_6$-FisB multimers. **C.** Gel filtration elution profile of His6-$FisB^{ECD}$ in Superose 6 Increase 10/300 GL column (top). Two fractions comprising the indicated peaks were re-injected in the same column under the same conditions, and eluted at the same volume as in the original sample. Elution volumes of molecular weight markers are indicated. **D.** Peaks labeled 1-3 in C were analyzed by Coomassie Blue-stained SDS-PAGE. Molecular weight markers are indicated on the left (in kDa). The band that corresponds to $His_6$-$FisB^{ECD}$ is indicated with a red arrow. The black asterisk indicates a chaperone that co-elutes with monomeric $His_6$-$FisB^{ECD}$. **E.** Representative electron micrographs of fractions comprising the 1st and 2nd peaks from C. Scale bar is 50nm. **F-H.** FisB ECD binding to liposomes is independent of calcium or pH, but decreases rapidly with increasing ionic strength. SUVs composed of 45 mole % CL and 55 mole % PC (40 nmol total lipid) were incubated with 200 pmol $His_6$-FisB ECD in buffers with the indicated [$Ca^{2+}$] (F), pH (G), or NaCl (H) for 1h and subjected to step-gradient isopycnic ultracentrifugation.

**S1 Appendix Figure 10. FisB mutants selectively deficient in membrane binding or oligomerization are expressed at similar levels as wild-type FisB. A.** Examples of cell contours detected using MicrobeJ. **B**. Distributions of background-corrected total fluorescence intensity per cell for *ΔfisB* cells expressing mYFP-$FisB^{WT}$ (BAL002), mYFP-$FisB^{KK}$ (BAL006), or mYFP-$FisB^{GIII}$ (BAL007) at low levels. The pixel values within the contours detected by MicrobeJ as in A were summed to define the total intensity per cell. This value was corrected for autofluorescence and background by subtracting the average total intensity per cell in cells (PY79) that did not express any fluorescent protein. The three distributions were indistinguishable, indicating that the mutants were expresses at the same level as the wild-type protein. **C**. Expression levels of mYFP-$FisB^{GIII}$ (BAL007) was similar to those of $FisB^{WT}$ (BAL002) using Western blotting, probed using an anti-FisB antibody. Time points into sporulation probed are indicated above the blot.

**S1 Appendix Figure 11. FisB mutants tested. A.** Mutations neutralizing 1-4 positively charged residues in the consensus region were introduced into FisB ECD, the mutants were expressed in *E. coli*, purified, and tested for binding to negatively charged liposomes using the flotation assay depicted in Fig. 4C. Neutralization of lysines around K170 produced the strongest reduction in binding. Liposomes were composed of 45 mole % CL and 55mole % PC**. B**. Other designed mutations targeted hydrophobic residues (black), inversion of positively (blue) or negatively (red) charged residues, or deletions. mYFP fusions of the mutated FisB were expressed at low levels in *ΔfisB* cells and tested for heat-resistant colony formation (C,D) and imaged for localization (E). **C.** Sporulation efficiency of cells expressing mYFP-FisB with deletion and hydrophobic residue mutations shown in B. **D**. Sporulation efficiency of cells expressing mYFP-FisB with charge inversion mutations shown in B. **E**. Images of sporulating cells (t=3 h) expressing mYFP-FisB bearing some of the mutations in C,D. In half the cases, the mYFP signal was cytosolic, suggesting the fusion protein was not inserted into the membrane and degraded (images boxed in red). In other cases, some mYFP signal was on the membrane and some was cytosolic (cyan-framed images). Cases in which mutants were located exclusively to the membrane were rare and included neutral mutations (images boxed in green) as well as $FisB^{KK}$ and $FisB^{GIII}$ shown in Fig. 6, at least under low expression levels.

**S1 Appendix Figure 12. *Alignment of B. subtilis and C. perfringens FisB sequences.*** *B. subtilis* (uniprot ID O32131, *YUNB_BACSU*) and *C. perfringens* (uniprot ID A0A0H2YVA3, *A0A0H2YVA3_CLOP1*) sequences were obtained from The Universal Protein Resource (UniProt) database (www.uniprot.org) and aligned using Clustal Omega[22] (https://www.ebi.ac.uk/Tools/msa/clustalo/).

#### SUPPLEMENTARY MOVIES

**Movie 1.** FisB forms dynamic foci in the mother-cell membranes. mGFP-FisB forms dynamic foci in the mother-cell membranes. The movie is a widefield microscopy time-lapse movie of strain BAM003, cropped to show a single cell (the parent stack consisted of 1024x1024 pixels) at t=2.5 hrs after nutrient downshift. For determining the MSD of DMCs we analyzed 60 different mobile FisB spots from 45 different bacteria. The focal plane was at the bottom surface of the cell. Images were acquired every 1 second. Scale bar 1 µm. Related to data in Fig 2E.

**Movie 2.** GFP-FisB foci are immobile at the cell pole at the time of membrane fission. The movie is a widefield microscopy time-lapse movie of strain BAM003, cropped to show a single cell (the parent stack consisted of 1024x1024 pixels) at t=3 hrs after nutrient downshift. For determining the MSD of ISEPs we analyzed 30 different FisB spots at the tip from 30 different bacteria at T3. The focal plane was at mid-cell. Images were acquired every second. Scale bar 1 µm. Related to data in Fig 2E.

**Movies 3 – 10.**

TIRFM movies were collected with an Olympus IX81 inverted microscope equipped with an Andor ixon-ultra-897 EM CCD camera. Cells of *B. subtilis* expressing either GFP-Mbl (strain BDR2061) or GFP-FisB (strain BMB14) were imaged on an 2% PBS agarose pad. GFP fusion proteins were excited with a 488 nm laser and exposures times were 50 ms for BDR2061 and 100 ms for BMB14. 1 frame/s. Scale bar 5 µm.

**Movie 3.** *B. subtilis* cells expressing mGFP-FisB (strain BM0B14). Related to data in S1 Appendix Fig. 6D-F.

**Movie 4.** A single cell from Movie 3 expressing mGFP-FisB. Related to data in S1 Appendix Fig. 6 C, top row.

**Movie 5**. *B. subtilis* cells expressing GFP-Mbl (strain BDR2061). Related to data in S1 Appendix Fig. 6B,E,F.

**Movie 6**. A single cell from Movie 5 expressing GFP-Mbl. Related to data in S1 Appendix Fig. 6 A, top row.

**Movie 7.** *B. subtilis* cells in the presence of 50 µg/ml fosfomycin expressing GFP-Mbl (strain BDR2061). Related to data in S1 Appendix Fig. 6B,E,F.

**Movie 8.** A single cell from Movie 7 in the presence of 50 µg/ml fosfomycin expressing GFP-Mbl. Related to data in S1 Appendix Fig. 6A, bottom row.

**Movie 9.** *B. subtilis* cells in the presence of 50 µg/ml fosfomycin expressing mGFP-FisB (strain BMB014). Related to data in S1 Appendix Fig. 6D-F.

**Movie 10.** A single cell from Movie 9 in the presence of 50 µg/ml fosfomycin expressing mGFP-FisB. Related to data in S1 Appendix Fig. 6C, bottom row.

**Table 1. Summary of FisB copy numbers per cell, ISEP and DMC in the native and low expression strains obtained using different calibrations**.

| Calibration method | *B. subtilis* standards | | DNA origami | | Quantitative WB | |
|---|---|---|---|---|---|---|
| Expression Level | low | native | low | native | low | native |
| Total copies of m(E)GFP-FisB per cell | 160 ±66[a] | 1300 ±500[c] | 122 ±51[f] | 963 ±376[h] | 83 ±6[n] | 661 ±49[k] |
| Copies at ISEP (t=3h) | 8±2[b] | 50±18[d] | 6±2[g] | 36±15[i] | 5±3[o] | 26±13[l] |
| Copies at DMC (t=2h) | n.d | 16±10[e] | n.d | 12±7[j] | n.d | 9±6[m] |

**a.** Total FisB copies per cell (mean±SD) were calculated from the distribution shown in Fig. S4B (red) and the calibration using the B. subtilis calibration strains ($y = 22.48x, R^2 = 0.93$, Fig. S2D). The error estimate is from the SD of the distribution in Fig. S4B. **b.** Calculated using mean±SD from the distribution shown in Fig. S4D (red) and the calibration with B. subtilis calibration strains, Fig. S2D. **c.** Calculated using mean±SD from the distribution shown in Figure S4B (green) and the calibration in Fig. S2D. **d**. Calculated using mean±SD of the distribution shown in Fig. S4D (green) and the calibration in Figure S2D. **e.** Calculated using mean±SD of the distribution shown in Fig. 2C (blue) and the calibration in Figure S2D. **f.** Calculated using mean±SD from the distribution shown in Fig. S4B (red) and the calibration obtained with DNA origami standards, Fig. 2D ($y = 29.56x; R^2 = 0.97$). **g**. Calculated using mean±SD of the distribution shown in Fig. S4D (red) and the calibration in Fig. 2D. **h**. Calculated using mean±SD of the distribution shown in Figure S4B (green) and the calibration in Fig. 2D. **i.** Calculated using mean±SD of the distribution shown in Fig. S4D (green) and the calibration in Fig. 2D. **j.** Calculated using mean±SD of the distribution shown in Fig. 2C (blue) and the calibration in Fig. 2D. **k**. Calculated from Western blot calibration using purified mYFP ($y = 2783, R^2 = 0.98$, Figure S3A) together with the Western blots of mXFP-FisB using a known number of cells (Figure S3B). mXFP-FisB from $1.44 \times 10^7$ B. subtilis cells loaded into the gel in Fig S3B produced a WB band intensity corresponding to $0.6 \pm 0.05$ ng ($6.61 \pm 0.49 \times 10^9$ molecules) in Fig. S3A. This was corrected for cleaved mXFP (~44%, Fig. S5B). **l**. Calculated using percentage of total cellular FisB located at ISEP estimated in Fig S2F and the total copies of mXFP-FisB per cell in k. **m**. Calculated as in l. **n.** Estimated by dividing FisB copies per cell (k) by 8. **o**. Calculated using mean±SD of the distribution of percentage cell o fluorescence located at ISEP in Figure S4E.

**Table 2.** ***Bacillus subtilis*** **strains used in this study.**

| Strain | Genotype | % spo[a] | Source |
|---|---|---|---|
| PY79 | *Prototrophic wild-type strain* | 100±8.36 | [23] |
| BDR1083 | *ΔfisB::tet* | 13.17 ± 1.97 | [24] |
| BKM15 | *amyE::PspoIIQ-cfp (spec)* | n.d | [25] |
| BAM003 | *ΔfisB::tet ycgO::PfisB-mGFP -fisB (cat)* | 90.17±12.65 | [25] |
| BVS001 | *ΔfisB::tet ycgO::PfisB-mYFP -fisB (cat)* | n.d | This work |
| BAL001 | *ΔfisB::tet ycgO::PfisB-mEGFP -fisB (cat)* | n.d | This work |
| BAL002 | *ΔfisB::tet, ycgO::PspoIID-RBSfisB(5n)-mYFP A206K-fisB (erm)* | 83.54 ± 4.93 | This work |
| BAL003 | *ΔfisB::tet, ycgO::PspoIID-RBSfisB(5n)-mGFP A206K-fisB (erm)* | n.d | This work |
| BAL004 | *ΔfisB::tet, ycgO::PspoIID-RBSfisB(5n)-mEGFP A206K-fisB (erm)* | n.d | This work |
| BAL005 | *ΔfisB::tet ycgO::PfisB-mEGFP-fisB Clostridum perfringens sp. (cat)* | 85 ± 5 | This work |
| BAL006 | *ΔfisB::tet, ycgO::PspoIID-RBSfisB(5n)-mYFP A206K-fisB* K168D K170D *(erm)* | 16.91±7.03 | This work |
| BAL007 | *ΔfisB::tet, ycgO::PspoIID-RBSfisB(5n)-mYFP A206K-fisB* G175A I176S I194T I195S *(erm)* | 3.03±2.51 | This work |
| BAL008 | *ΔfisB::tet, ycgO::PspoIID-RBSfisB(5n)-mYFP A206K-fisB Δ80-96 (erm)* | 13.04±2.71 | This work |
| BAL009 | *ΔfisB::tet, ycgO::PspoIID-RBSfisB(5n)-mYFP A206K-fisB Δ122-132 (erm)* | 99.42±15.03 | This work |
| BAL010 | *ΔfisB::tet, ycgO::PspoIID-RBSfisB(5n)-mYFP A206K-fisB Δ137-154 (erm)* | 14.09±6.33 | This work |
| BAL011 | *ΔfisB::tet, ycgO::PspoIID-RBSfisB(5n)-mYFP A206K-fisB Δ167-182 (erm)* | 16.70±6.99 | This work |
| BAL012 | *ΔfisB::tet, ycgO::PspoIID-RBSfisB(5n)-mYFP A206K-fisB Δ210-220 (erm)* | 15.30±3.32 | This work |
| BAL013 | *ΔfisB::tet, ycgO::PspoIID-RBSfisB(5n)-mYFP A206K-fisB* Δ132-222*(erm)* | 14.78±3.40 | This work |

| | | | |
|---|---|---|---|
| BAL014 | *ΔfisB::tet, ycgO::PspoIID-RBSfisB(5n)-mYFP A206K-fisB* L90T ( erm) | 104.31±0.00 | This work |
| BAL015 | *ΔfisB::tet, ycgO::PspoIID-RBSfisB(5n)-mYFP A206K-fisB L137S G138A (erm)* | 16.74±2.39 | This work |
| BAL016 | *ΔfisB::tet, ycgO::PspoIID-RBSfisB(5n)-mYFP A206K-fisB L145TL146S (erm)* | 16.52±2.60 | This work |
| BAL017 | *ΔfisB::tet, ycgO::PspoIID-RBSfisB(5n)-mYFP A206K-fisB G150A (erm)* | 23.91±3.04 | This work |
| BAL018 | *ΔfisB::tet, ycgO::PspoIID-RBSfisB(5n)-mYFP A206K-fisB* V215S G217A *(erm)* | 10.43±1.73 | This work |
| BAL019 | *ΔfisB::tet, ycgO::PspoIID-RBSfisB(5n)-mYFP A206K-fisB* V219T *(erm)* | 23.48±0.00 | This work |
| BAL020 | *ΔfisB::tet, ycgO::PspoIID-RBSfisB(5n)-mYFP A206K-fisB* G175A I176S *(erm)* | 9.91±5.14 | This work |
| BAL021 | *ΔfisB::tet, ycgO::PspoIID-RBSfisB(5n)-mYFP A206K-fisB* I194T I195S (erm) | 45.65±12.97 | This work |
| BAL022 | *ΔfisB::tet, ycgO::PspoIID-RBSfisB(5n)-mYFP A206K-fisB* R56E *(erm)* | 55.00±15.00 | This work |
| BAL023 | *ΔfisB::tet, ycgO::PspoIID-RBSfisB(5n)-mYFP A206K-fisB* E67R D68K *(erm)* | 37.30±12.70 | This work |
| BAL024 | *ΔfisB::tet, ycgO::PspoIID-RBSfisB(5n)-mYFP A206K-fisB* K106D K109D *(erm)* | 22.50±7.50 | This work |
| BAL025 | *ΔfisB::tet, ycgO::PspoIID-RBSfisB(5n)-mYFP A206K-fisB* K116D *(erm)* | 65.70±14.30 | This work |
| BAL026 | *ΔfisB::tet, ycgO::PspoIID-RBSfisB(5n)-mYFP A206K-fisB* E119R *(erm)* | 55.00±15.00 | This work |
| BAL027 | *ΔfisB::tet, ycgO::PspoIID-RBSfisB(5n)-mYFP A206K-fisB* R156E *(erm)* | 65.50±22.50 | This work |
| BAL028 | *ΔfisB::tet, ycgO::PspoIID-RBSfisB(5n)-mYFP A206K-fisB K170D K172DE* (erm) | 16.40±1.45 | This work |
| BAL029 | *ΔfisB::tet, ycgO::PspoIID-RBSfisB(5n)-mYFP A206K-fisB* K192D *(erm)* | 55.16±14.80 | This work |
| BAM234 | *ΔywnE::cat ΔywjE::kan ΔywiE::erm* | 25.88 ± 8.53 | [25] |
| BAM236 | *ΔfisB::tet ΔywnE::cat ΔywjE::kan ΔywiE::erm* | 1.07 ± 0.52 | [25] |

| | | | |
|---|---|---|---|
| BAL030 | *ΔywnE ΔywjE ΔywiE::kan ΔpssA::erm* | 25.64 ± 2.56 | This work |
| BAL031 | *ΔpssA::erm* | 85.10 ± 12.16 | This work |
| BAL032 | *ΔltaSA::erm* | 113.8 ± 27.24 | This work |
| BAL033 | *ΔugtP::erm* | 74.04 ± 22.12 | This work |
| BAL034 | *ΔmprF::erm* | 119.2± 47.44 | This work |
| BAL035 | *ΔfloA::erm* | 35.9 ±12.18 | This work |
| BAL036 | *ΔfloT::erm* | 103.8±11.54 | This work |
| BAL037 | *ΔfisB::tet ΔywnE ΔywjE ywiE::kan ycgO::PspoIID-RBSfisB(5n)-mYFP -fisB (erm)* | n.d | This work |
| BDR2061 | *amyE::PxylA-gfp-mbl (spec), mblΩpMUTIN4 (erm) trpC2* | n.d | [12] |
| BMB014 | *amyE::PxylA-mciZ(cat), ycgO::Pspank-gfp-fisB (spec)* | n.d | This work |
| BMB031 | *ycgO::PfisB-RBSfisB-Myc-fisB(erm)* | 101 ± 7 | This work |
| BMB032 | *ycgO::PfisB-RBSfisB-Myc-fisB(G6C) (erm)* | 68 ± 19 | This work |
| BMB033 | *ycgO::PfisB-RBSfisB-Myc-fisB(L137C) (erm)* | 53 ± 3 | This work |
| BMB034 | *ycgO::PfisB-RBSfisB-Myc-fisB(A245C) (erm)* | 83 ± 8 | This work |
| BS168 | *Wild-type strain trpC2* | n.d | BGSC[b] |
| 1A1246 | *amyE::(Pveg(+1/+8)_R1-18_sfGFP_spec) trpC2* | n.d | BGSC |
| 1A1243 | *amyE::(Pveg(+1/+8)_R1-15_sfGFP_spec) trpC2* | n.d | BGSC |
| 1A1227 | *amyE::(Pveg(+1/+8)_R0-16_sfGFP_spec) trpC2* | n.d | BGSC |
| 1A1241 | *amyE::(Pveg(+1/+8)_R1-13_sfGFP_spec) trpC2* | n.d | BGSC |
| 1A1239 | *amyE::(Pveg(+1/+8)_R1-11_sfGFP_spec) trpC2* | n.d | BGSC |

| | | | |
|---|---|---|---|
| 1A1237 | *amyE::(Pveg(+1/+8)_R1-9_sfGFP_spec) trpC2* | n.d | BGSC |
| 1A1220 | *amyE::(Pveg(+1/+8)_R0-9_sfGFP_spec) trpC2* | n.d | BGSC |
| SG13 | *amyE::(Pveg_R0_sfGFP_spec)* | n.d | BGSC |
| BAL038 | *amyE::(Pveg_R0_mEGFP_spec)* | n.d | This work |

**a:** Sporulation efficiency (% of WT spores at 24h after the onset of sporulation) for each indicated strain. Results are shown as means ± SD for four replicates per condition.

**b:** Bacillus Genetic Stock Center (www.bgsc.org)

**Table 3. Plasmids used in this study.**

| Plasmid | Genotype | E.Coli | Source |
|---|---|---|---|
| pAM002 | *ΔfisB::tet ycgO::PfisB-mGFP -fisB (cat)* | amp | [25] |
| pVS001 | *ΔfisB::tet ycgO::PfisB-mYFP A206K -fisB (cat)* | amp | This work |
| pAL001 | *ycgO::PfisB-mEGFP-fisB B.subtilis.(cat)* | amp | This work |
| pAL002 | *ycgO::PspoIID-RBSfisB(5n)-mEGFP-fisB (erm)* | amp | This work |
| pAL003 | *ycgO::PfisB-mEGFP-fisB Clostridium Perfringens sp.(cat)* | amp | This work |
| pAL004 | *ycgO::PspoIID-RBSfisB(5n)-mYFP A206K-fisB (erm)* | amp | This work |
| pAL005 | *ycgO::PspoIID-RBSfisB(5n)-mYFP A206K-fisB K168DK170D (erm)* | amp | This work |
| pAL006 | *ycgO::PspoIID-RBSfisB(5n)-mYFP A206K-fisB G175A I176S I194TI195S (erm)* | amp | This work |
| pKM110 | *his6-fisB$^{ECD}$ WT* | amp | [25] |
| pAL007 | *his6-fisB$^{ECD}$ WT* G123C | amp | This work |
| pDT390 | *his6-fisB$^{ECD}$ G175A I176S I194T I195S* | amp | [25] |
| pAL008 | *his6-fisB$^{ECD}$ G175A I176S I194T I195S G13C* | amp | This work |
| pAL009 | *his6-fisB$^{ECD}$* K168D K170D | amp | This work |
| pAL010 | *his6-fisB$^{ECD}$* K168D K170D G123C | amp | This work |
| pAL011 | *ycgO::PspoIID-RBSfisB(5n)-mYFP A206K-fisB Δ80-96 (erm)* | amp | This work |
| pAL012 | *ycgO::PspoIID-RBSfisB(5n)-mYFP A206K-fisB Δ122-132 (erm)* | amp | This work |
| pAL013 | *ycgO::PspoIID-RBSfisB(5n)-mYFP A206K-fisB Δ137-154 (erm)* | amp | This work |
| pAL014 | *ycgO::PspoIID-RBSfisB(5n)-mYFP A206K-fisB Δ167-182 (erm)* | amp | This work |
| pAL015 | *ycgO::PspoIID-RBSfisB(5n)-mYFP A206K-fisB Δ210-220 (erm)* | amp | This work |

| | | | |
|---|---|---|---|
| pAL016 | *ycgO::PspoIID-RBSfisB(5n)-mYFP A206K-fisB Δ132-222 (erm)* | amp | This work |
| pAL017 | *ycgO::PspoIID-RBSfisB(5n)-mYFP A206K-fisB L90T (erm)* | amp | This work |
| pAL018 | *ycgO::PspoIID-RBSfisB(5n)-mYFP A206K-fisB L137SG138A (erm)* | amp | This work |
| pAL019 | *ycgO::PspoIID-RBSfisB(5n)-mYFP A206K-fisB L145TL146S (erm)* | amp | This work |
| pAL020 | *ycgO::PspoIID-RBSfisB(5n)-mYFP A206K-fisB G150A (erm)* | amp | This work |
| pAL021 | *ycgO::PspoIID-RBSfisB(5n)-mYFP A206K-fisB V215SG217A (erm)* | amp | This work |
| pAL022 | *ycgO::PspoIID-RBSfisB(5n)-mYFP A206K-fisB V219T (erm)* | amp | This work |
| pAL023 | *ycgO::PspoIID-RBSfisB(5n)-mYFP A206K-fisB G175AI176S (erm)* | amp | This work |
| pAL024 | *ycgO::PspoIID-RBSfisB(5n)-mYFP A206K-fisB I194TI195S (erm)* | amp | This work |
| pAL025 | *ycgO::PspoIID-RBSfisB(5n)-mYFP A206K-fisB* R56E *(erm)* | amp | This work |
| pAL026 | *ycgO::PspoIID-RBSfisB(5n)-mYFP A206K-fisB E67RD68K (erm)* | amp | This work |
| pAL027 | *ycgO::PspoIID-RBSfisB(5n)-mYFP A206K-fisB K106D K109D (erm)* | amp | This work |
| pAL028 | *ycgO::PspoIID-RBSfisB(5n)-mYFP A206K-fisB K116D (erm)* | amp | This work |
| pAL029 | *ycgO::PspoIID-RBSfisB(5n)-mYFP A206K-fisB E119R (erm)* | amp | This work |
| pAL030 | *ycgO::PspoIID-RBSfisB(5n)-mYFP A206K-fisB R156E (erm)* | amp | This work |
| pAL031 | *ycgO::PspoIID-RBSfisB(5n)-mYFP A206K-fisB K170D K172DE (erm)* | amp | This work |
| pDR244 | PPA-cre-ori(ts) spec | amp | BGSC |
| pMB062 | *ycgO::PfisB-RBSfisB-Myc-fisB(erm)* | amp | This work |

| | | | |
|---|---|---|---|
| pMB064 | *ycgO::PfisB-RBSfisB-Myc-fisB(G6C) (erm)* | amp | This work |
| pMB065 | *ycgO::PfisB-RBSfisB-Myc-fisB(L137C) (erm)* | amp | This work |
| pMB066 | *ycgO::PfisB-RBSfisB-Myc-fisB(A245C) (erm)* | amp | This work |
| pVS002 | his6-SUMO-mYFP | kan | This work |
| pECE321 | Pveg_R0_sfGFP | amp | BGSC |
| pECE321_mEGFP | Pveg_R0_mEGFP | amp | This work |

**REFERENCES**


1. Guiziou S*, et al.* A part toolbox to tune genetic expression in Bacillus subtilis. *Nucleic Acids Res* **44**, 7495-7508 (2016).

2. Ojkic N, López-Garrido J, Pogliano K, Endres RG. Cell-wall remodeling drives engulfment during Bacillus subtilis sporulation. *Elife* **5**, (2016).

3. Carballido-Lopez R, Formstone A, Li Y, Ehrlich SD, Noirot P, Errington J. Actin homolog MreBH governs cell morphogenesis by localization of the cell wall hydrolase LytE. *Dev Cell* **11**, 399-409 (2006).

4. Carballido-Lopez R, Formstone A. Shape determination in Bacillus subtilis. *Curr Opin Microbiol* **10**, 611-616 (2007).

5. Strahl H, Hamoen LW. Membrane potential is important for bacterial cell division. *Proc Natl Acad Sci U S A* **107**, 12281-12286 (2010).

6. Meyer P, Gutierrez J, Pogliano K, Dworkin J. Cell wall synthesis is necessary for membrane dynamics during sporulation of Bacillus subtilis. *Mol Microbiol* **76**, 956-970 (2010).

7. Shaevitz JW, Gitai Z. The structure and function of bacterial actin homologs. *Cold Spring Harb Perspect Biol* **2**, a000364 (2010).

8. Garner EC, Bernard R, Wang W, Zhuang X, Rudner DZ, Mitchison T. Coupled, circumferential motions of the cell wall synthesis machinery and MreB filaments in B. subtilis. *Science* **333**, 222-225 (2011).

9. Schirner K*, et al.* Lipid-linked cell wall precursors regulate membrane association of bacterial actin MreB. *Nat Chem Biol* **11**, 38-45 (2015).

10. Silver LL. Fosfomycin: Mechanism and Resistance. *Cold Spring Harb Perspect Med* **7**, (2017).

11. Axelrod D. Total internal reflection fluorescence microscopy in cell biology. *Traffic* **2**, 764-774 (2001).

12. Carballido-López R, Errington J. The bacterial cytoskeleton: in vivo dynamics of the actin-like protein Mbl of Bacillus subtilis. *Dev Cell* **4**, 19-28 (2003).

13. Huet S, Karatekin E, Tran VS, Fanget I, Cribier S, Henry JP. Analysis of transient behavior in complex trajectories: application to secretory vesicle dynamics. *Biophys J* **91**, 3542-3559 (2006).

14. El-Gebali S*, et al.* The Pfam protein families database in 2019. *Nucleic Acids Res* **47**, D427-D432 (2019).

15. Bogdanov M, Heacock PN, Dowhan W. Study of polytopic membrane protein topological organization as a function of membrane lipid composition. *Methods Mol Biol* **619**, 79-101 (2010).

16. Ovchinnikov S*, et al.* Protein structure determination using metagenome sequence data. *Science* **355**, 294-298 (2017).

17. Sharp MD, Pogliano K. An in vivo membrane fusion assay implicates SpoIIIE in the final stages of engulfment during Bacillus subtilis sporulation. *Proc Natl Acad Sci U S A* **96**, 14553-14558 (1999).

18. Smith MB, Li H, Shen T, Huang X, Yusuf E, Vavylonis D. Segmentation and tracking of cytoskeletal filaments using open active contours. *Cytoskeleton (Hoboken)* **67**, 693-705 (2010).

19. Smith MB, Karatekin E, Gohlke A, Mizuno H, Watanabe N, Vavylonis D. Interactive, computer-assisted tracking of speckle trajectories in fluorescence microscopy: application to actin polymerization and membrane fusion. *Biophys J* **101**, 1794-1804 (2011).

20. Landau M*, et al.* ConSurf 2005: the projection of evolutionary conservation scores of residues on protein structures. *Nucleic Acids Res* **33**, W299-302 (2005).

21. Dobson L, Reményi I, Tusnády GE. CCTOP: a Consensus Constrained TOPology prediction web server. *Nucleic Acids Res* **43**, W408-412 (2015).

22. Sievers F*, et al.* Fast, scalable generation of high-quality protein multiple sequence alignments using Clustal Omega. *Mol Syst Biol* **7**,  (2011).

23. Youngman PJ, Perkins JB, Losick R. Genetic transposition and insertional mutagenesis in Bacillus subtilis with Streptococcus faecalis transposon Tn917. *Proc Natl Acad Sci U S A* **80**, 2305-2309 (1983).

24. Eichenberger P*, et al.* The sigmaE regulon and the identification of additional sporulation genes in Bacillus subtilis. *J Mol Biol* **327**, 945-972 (2003).

25. Doan T*, et al.* FisB mediates membrane fission during sporulation in Bacillus subtilis. *Genes Dev* **27**, 322-334 (2013).

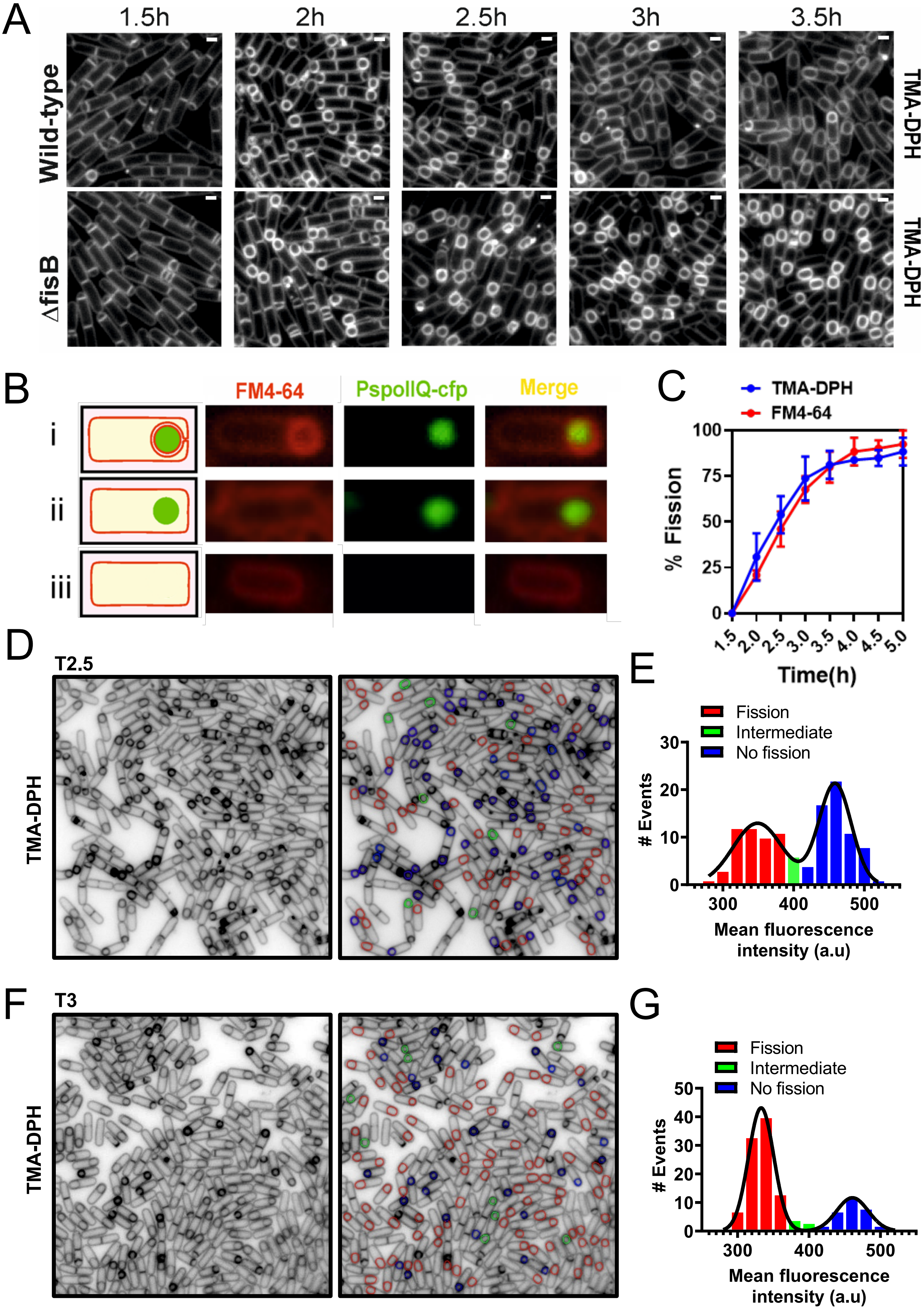


**S1 APPENDIX FIGURE 1**

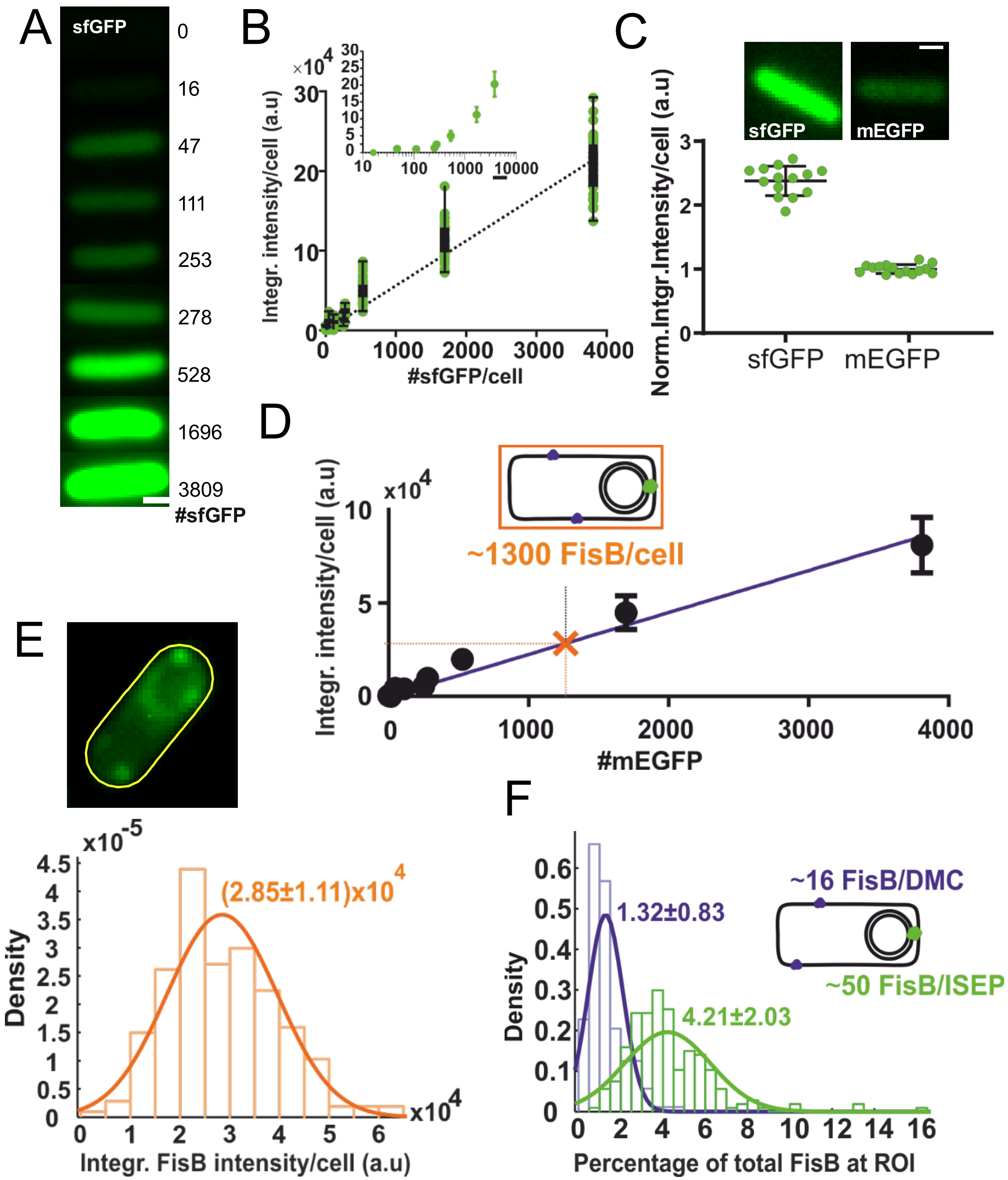


**S1 APPENDIX FIGURE 2**

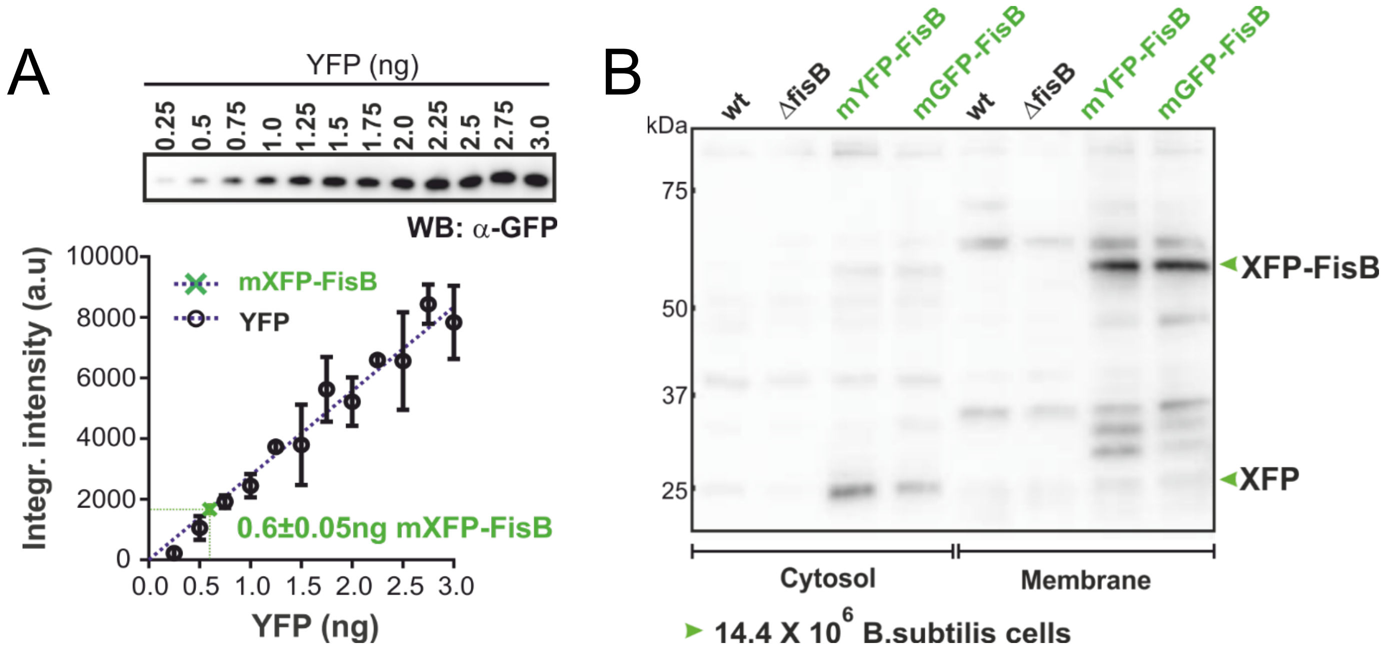


### S1 APPENDIX FIGURE 3

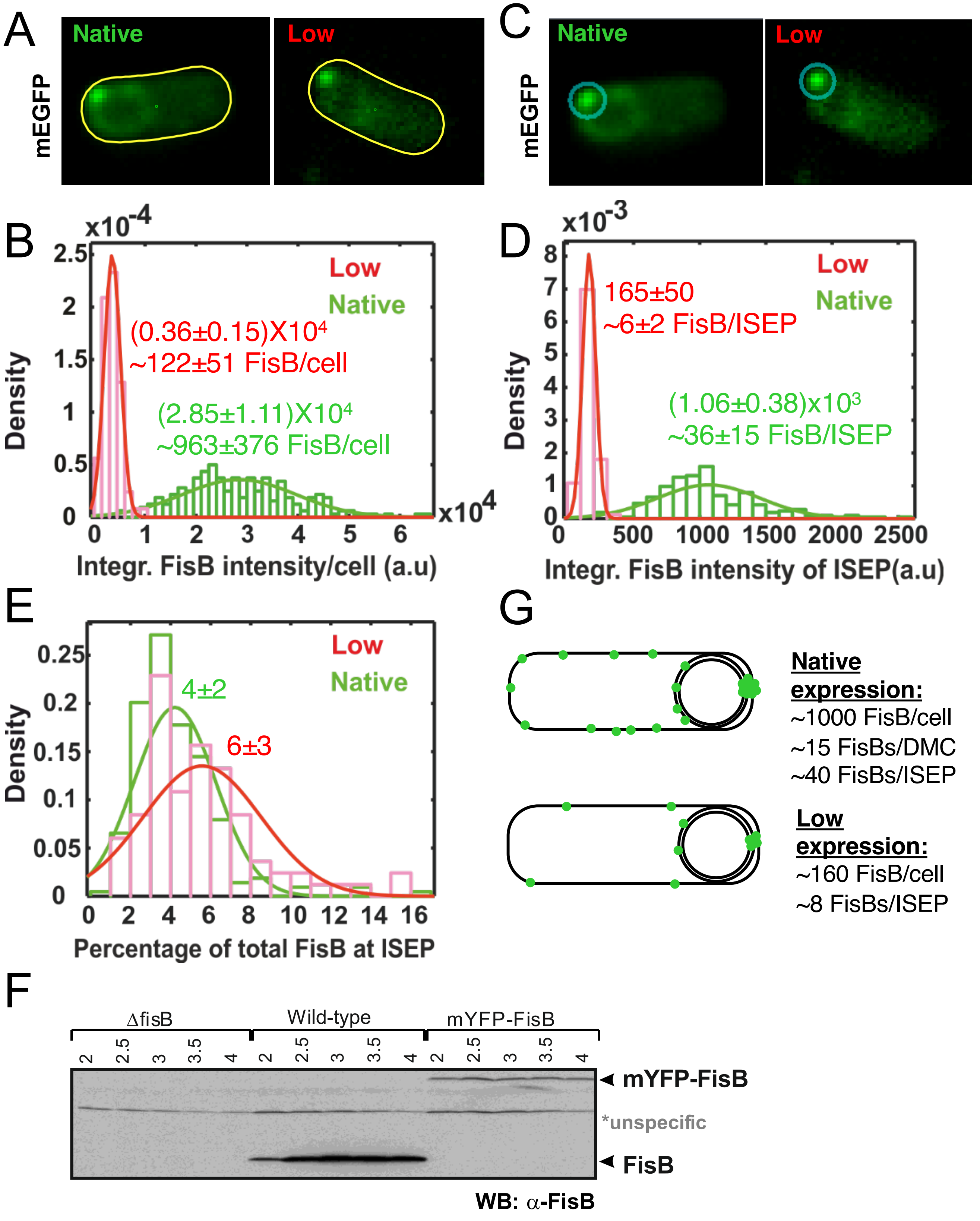


### S1 APPENDIX FIGURE 4

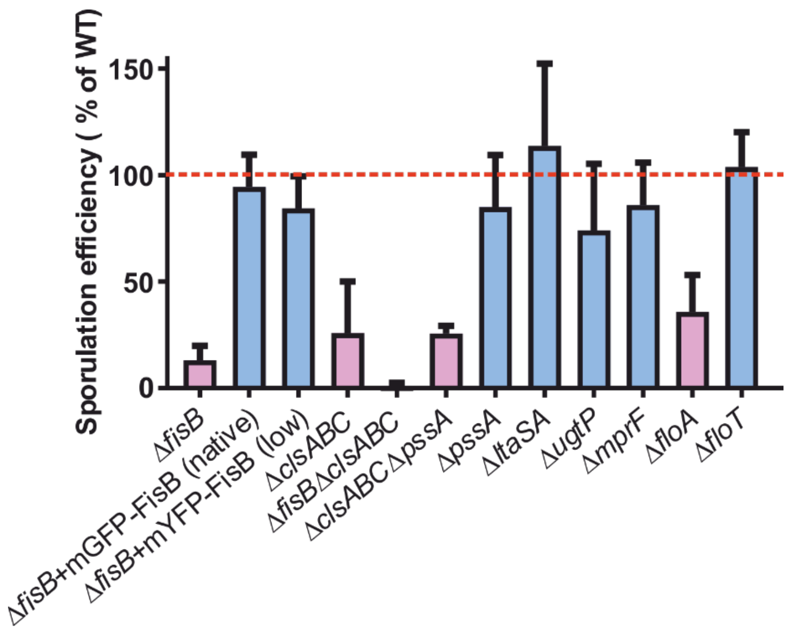


**S1 APPENDIX FIGURE 5**

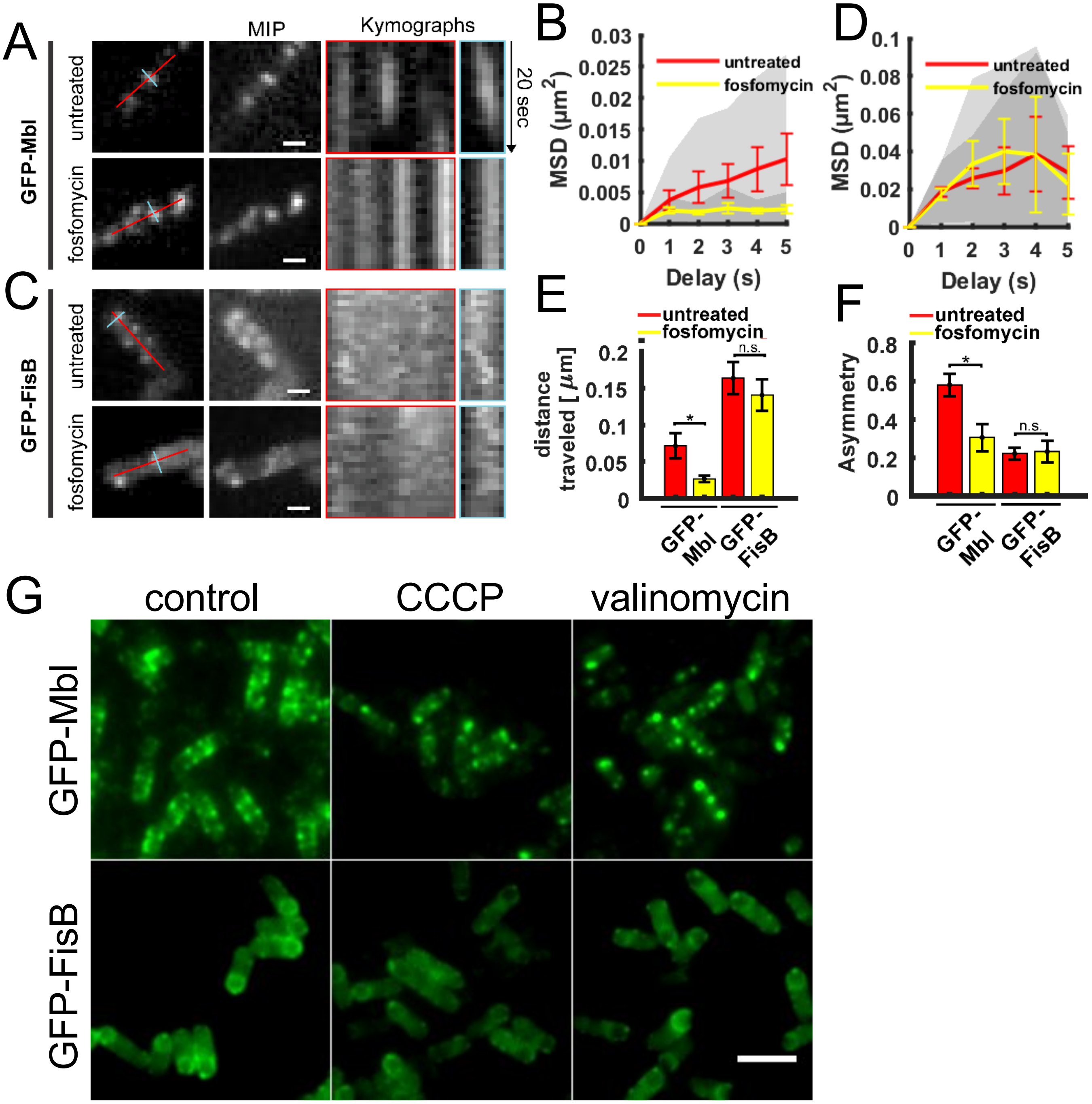


**S1 APPENDIX FIGURE 6**

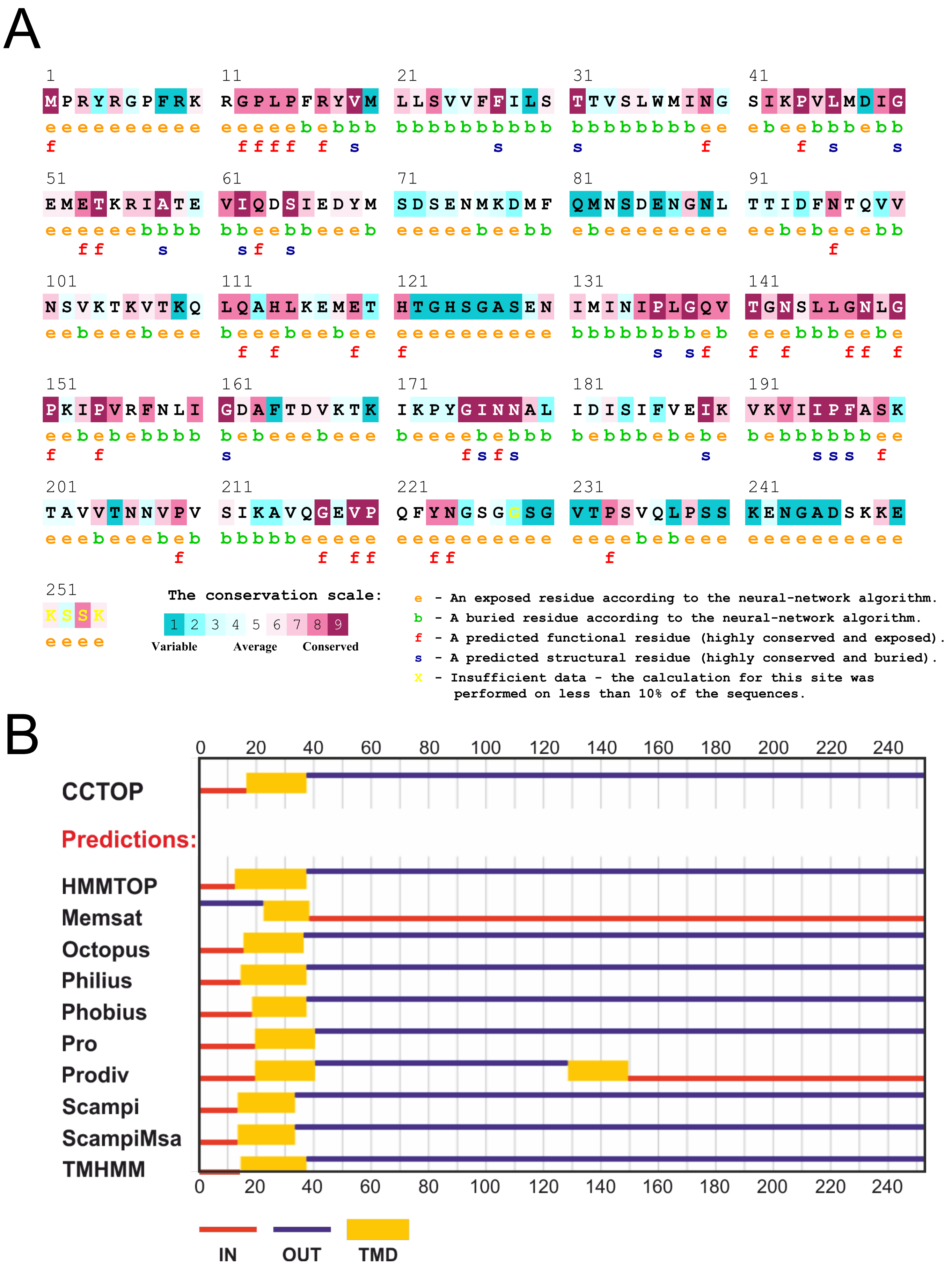


S1 APPENDIX FIGURE 7

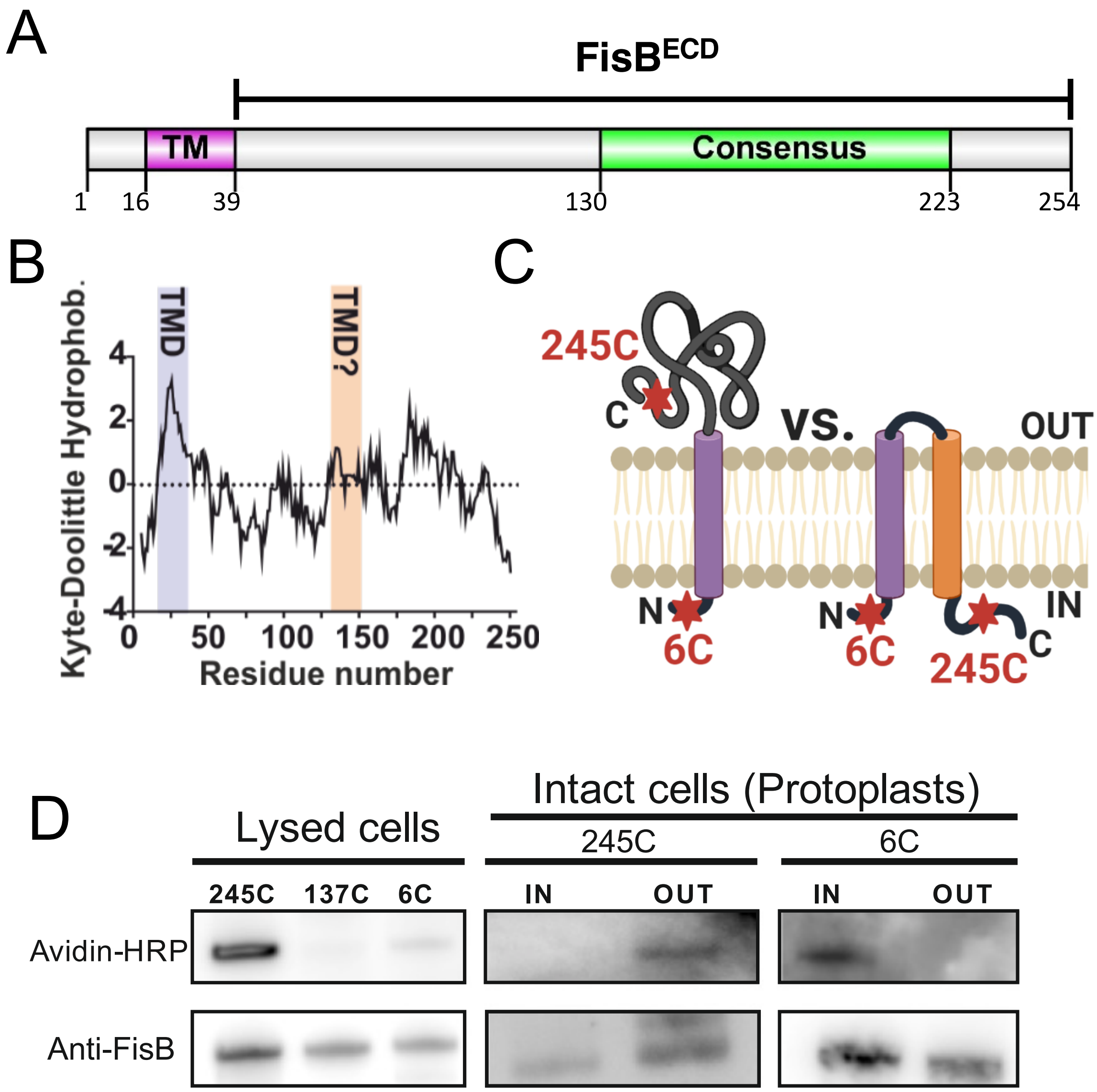


**S1 APPENDIX FIGURE 8**

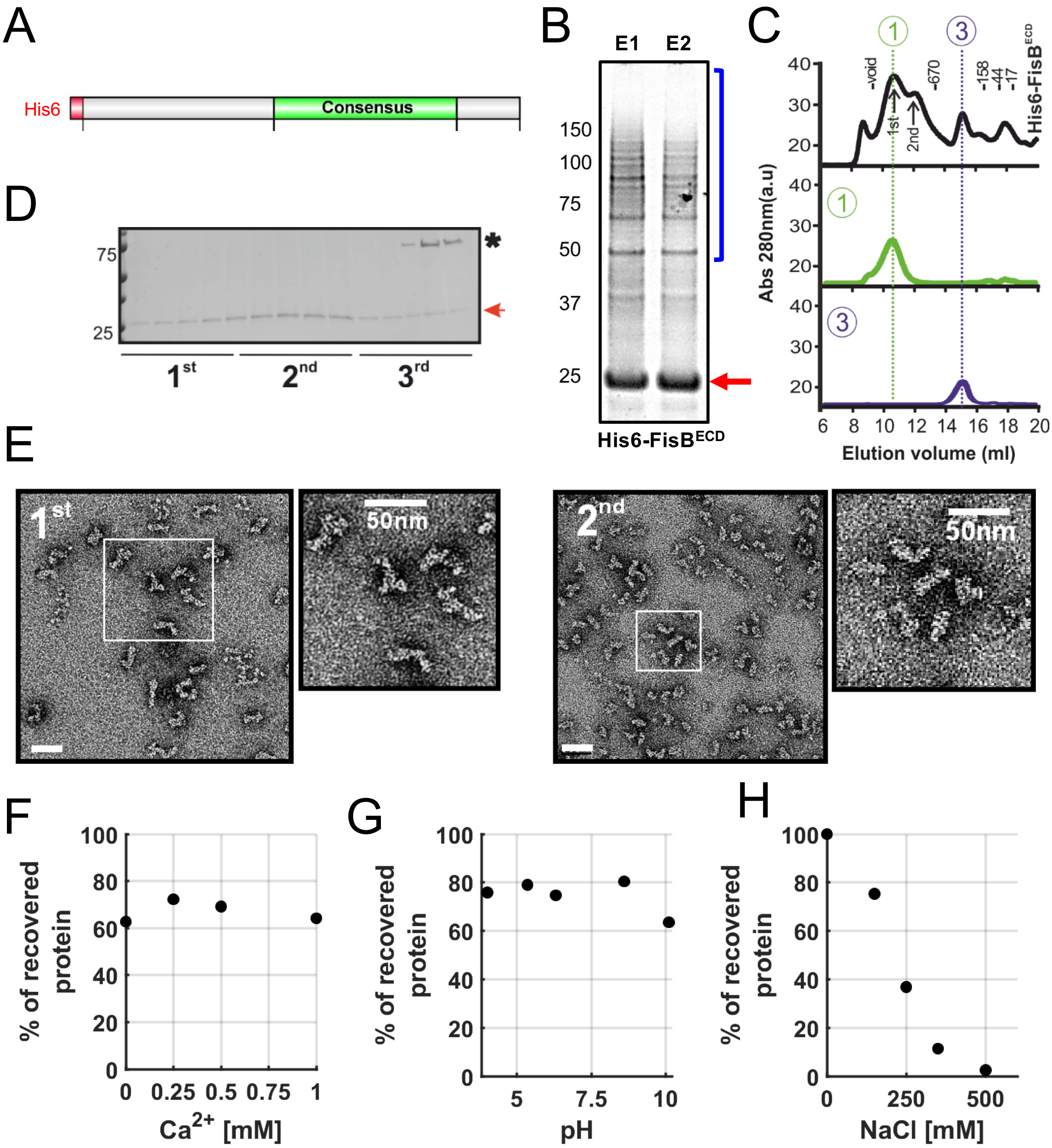


**S1 APPENDIX FIGURE 9**

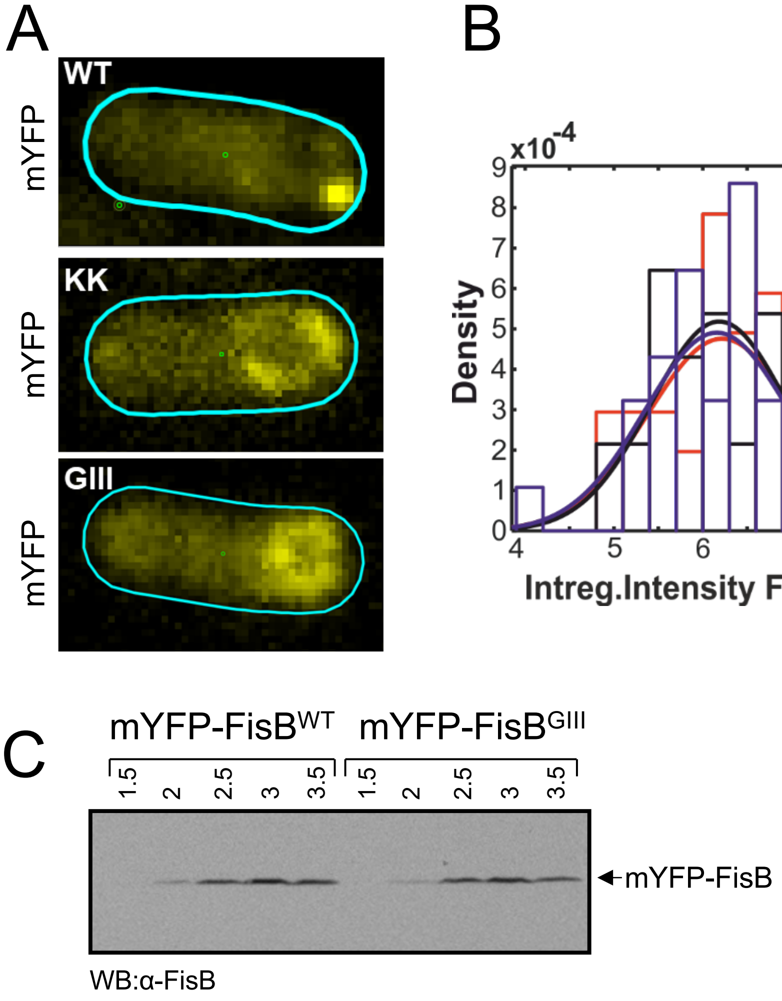


### S1 APPENDIX FIGURE 10

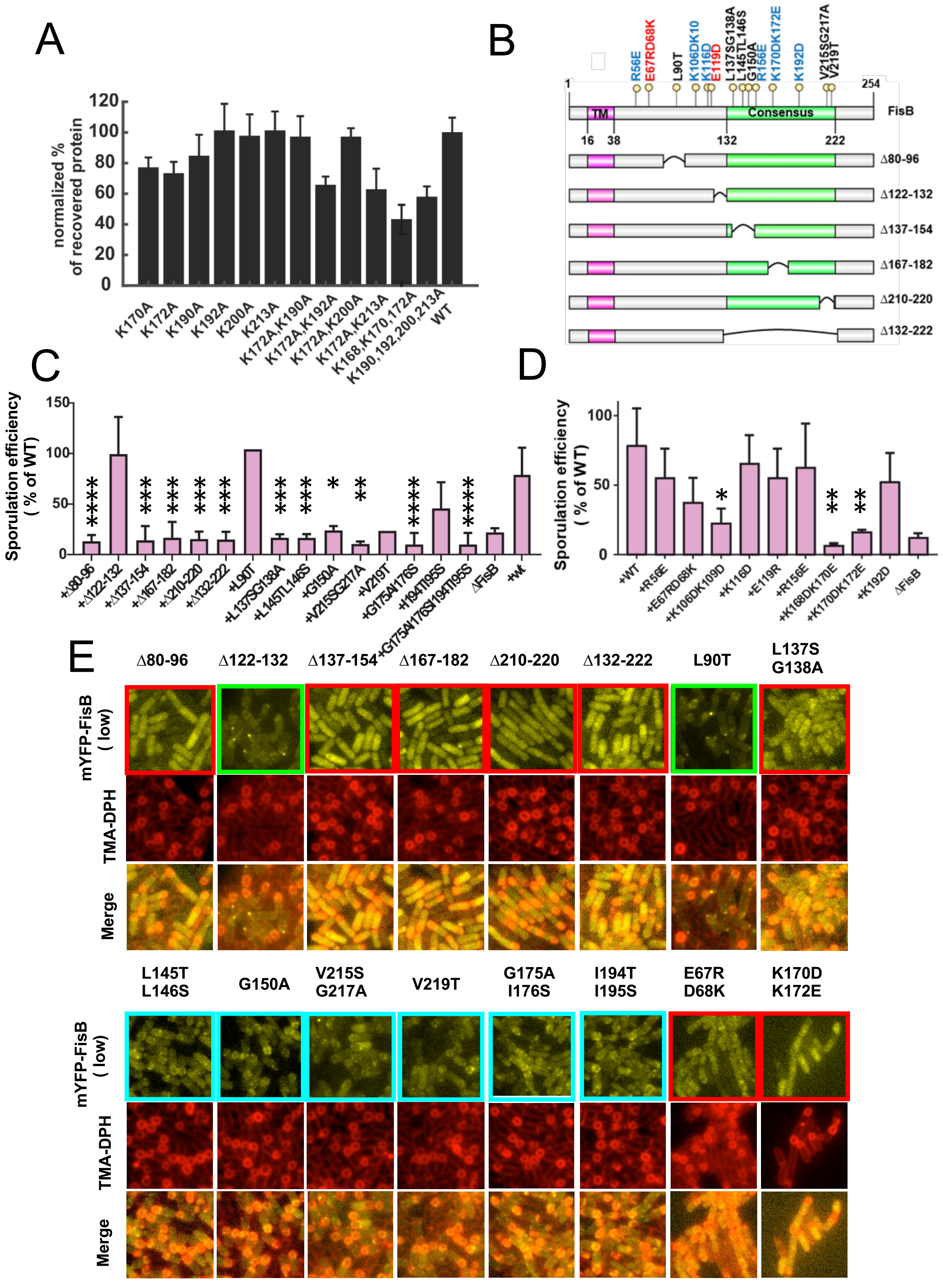


### S1 APPENDIX FIGURE 11

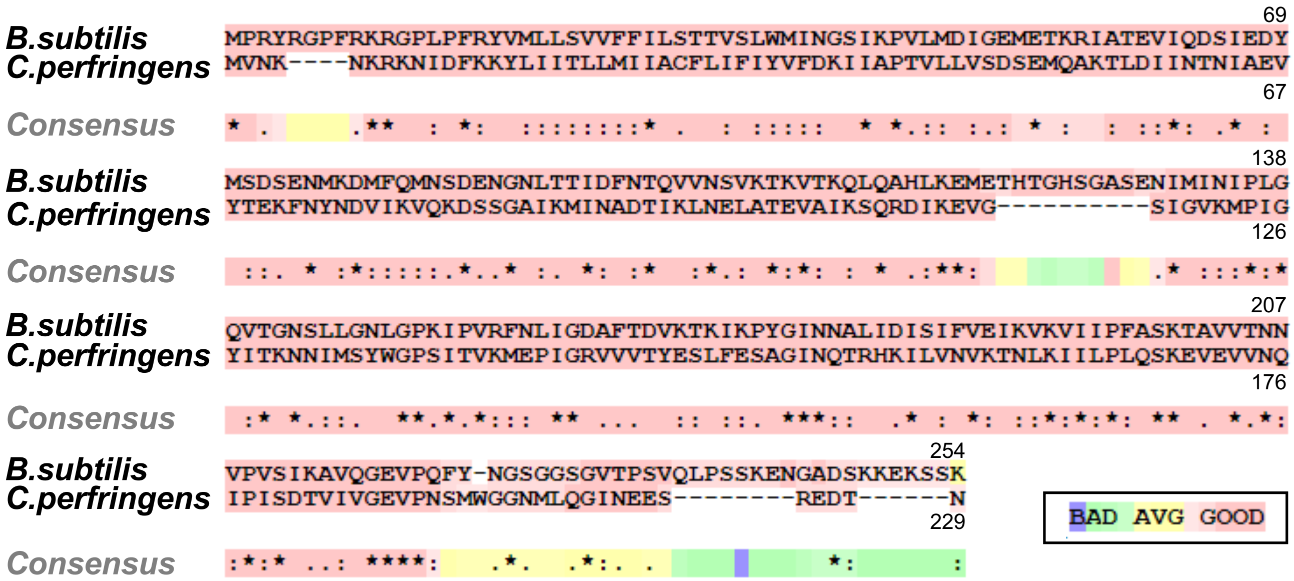


| | %Identity | Net Charge | Hydropathicity (GRAVY) |
|---|---|---|---|
| *Clostridium perfringens* | 23.50% | + | 0.055 |
| *Bacillus subtilis* | 100% | + | -0.160 |

### S1 APPENDIX FIGURE 12
